# Large-scale analyses reveal the contribution of adaptive evolution in pathogenic and non-pathogenic fungal species

**DOI:** 10.1101/2023.08.28.555124

**Authors:** Danilo Pereira, Melvin D Bolton, Timothy L Friesen, Wolfgang Stephan, Julien Y Dutheil, Eva H Stukenbrock

## Abstract

Genome studies of fungal pathogens have presented evidence for exceptionally high rates of evolution. It has been proposed that rapid adaptation is a hallmark of pathogen evolution that facilitates the invasion of new host niches and the overcoming of intervention strategies such as fungicide applications and drug treatments. To which extent high levels of genetic variation within and between species correlate with adaptive protein evolution in fungi more generally has so far not been explored. In this study, we addressed the contribution of adaptive evolution relative to genetic drift in 20 fungal species, hereby exploring genetic variation in 2,478 fungal genomes. We reannotated positions of protein-coding genes to obtain a high-quality dataset of 234,427 full-length core gene and 25,612 accessory gene alignments. We applied an extension of the McDonald-Kreitman test that models the distributions of fitness effects to infer the rate of adaptive (ω_A_) and non-adaptive (ω_NA_) non-synonymous substitutions in protein-coding genes. To explore the relevance of recombination on local adaptation rates, we inferred the population genomic recombination rate for all 20 species. Our analyses reveal extensive variation in rates of adaptation and show that high rates of adaptation are not a hallmark of a pathogenic lifestyle. Up to 83% of non-synonymous substitutions are adaptive in the species *Parastagonospora nodorum*. However, non-synonymous substitutions in other species, including the prominent rice-infecting pathogen *Magnaporthe oryzae*, are predominantly non-adaptive (neutral or slightly deleterious). Correlating adaptation measures with effective population size and recombination rate, we show that effective population size is a primary determinant of adaptive evolution in fungi. At the genome scale, recombination rate variation explains variation in both ω_A_ and ω_NA_. Finally, we demonstrate the robustness of our estimates using simulations. We underline the value of population genetic principles in studies of fungal evolution, and we highlight the importance of demographic processes in adaptive evolution of pathogenic and non-pathogenic species.

## Introduction

Standing genetic variation and new mutations are central to species’ long-term survival. Non-synonymous mutations in protein-coding genes are an important contributor to the adaptive potential of any species given the functional roles of proteins. Furthermore, the universal properties of the genetic code permit the development of models of codon sequence evolution. Consequently, the statistical inference of the dynamic of fixation of this class of mutations is unparalleled and allows to unravel the general dynamics of adaptation at the molecular level.

During evolution, the load of mutations segregating within populations varies quantitatively depending on the fitness changes they cause (1). Adaptive evolution occurs when mutations with beneficial effects on fitness increase in frequency towards fixation by the action of natural selection (2). By modulating genetic drift, demographic processes impact the frequency of genetic variants in populations. Hereby, a reduced population size dilutes the influence of natural selection, slowing down the fixation of advantageous mutations and maintaining slightly deleterious mutations in higher frequencies (3). This dynamic of adaptive and non-adaptive allele frequency changes significantly impacts the antagonistic co-evolution of pathogens and hosts. For fungal pathogens, a direct consequence of pathogen adaptation is often reported as outbreaks threatening human health, conservation efforts and food production (4). Human action in the natural environment and agro-ecosystems lead to changes in selective pressures imposed on pathogen populations and frequent fluctuation in pathogen population size (5). Yet, plant pathogens often feature among species with high adaptive potential, and it has been proposed that their ability to adapt exceeds that of other eukaryotic organisms (5–7).

However, despite the consequences of fungal pathogen adaptation, efforts to quantify the contribution of adaptive evolution and genetic drift to molecular evolution have mainly focused on individual species and reduced sample sizes (8,9). Hence, a large-scale multispecies approach is needed to elucidate general patterns that govern adaptive evolution and the mechanisms that shape rates of adaptation in fungal pathogens.

Molecular adaptive evolution can be measured at the level of the protein sequence composition. The McDonald-Kreitman (MK) test is based on comparing within and between species variation at non-synonymous and synonymous sites in protein-coding sequences (10). The rationale of the test is that adaptive evolution is reflected as a higher proportion of non-synonymous to synonymous fixed substitutions compared to the proportion of non-synonymous and synonymous polymorphisms segregating within the species. Developments of the MK test consider that not all substitutions at non-synonymous sites are adaptive. Some may be neutral or even slightly deleterious (11). MK-based statistics distinguish the proportion of non-synonymous mutations fixed by positive selection (α) and the rates of adaptive (ω_A_) and non-adaptive (ω_NA_) non-synonymous substitutions (12,13,3,14). Several studies have investigated the prevalence of positive selection across different taxa, primarily focusing on Hominids (3,15) and Drosophila (16–18, but more recently also Arabidopsis (18), non-model animals (14), bacteria (19) and individual species of fungi (8,9,20). A common determinant of adaptive evolution identified across very different species is effective population size (Ne) (3,14,21,22). In higher-Ne species, natural selection underlies the fixation of beneficial alleles concurrent with the removal of deleterious alleles, reducing the influence of genetic drift. However, the finding that some high-Ne species show no evidence of adaptive evolution indicates the contribution of other factors (i.e., *Zea mays* in Gossmann et al., 2012). Genome-wide features were also shown to influence rates of adaptive evolution. Recombination increases the probability of fixing beneficial mutations and facilitates the purging of deleterious changes by mitigating the effect of linked selection (24,25,19). Furthermore, some genes may locate in genomic regions where more mutations arise. Finally, the distribution of fitness effects, in particular the proportion of advantageous mutations, may differ greatly between genes, leading to differential substitution rates between functional categories. High rates of positively-selected mutations accumulating in genes involved in immune response (26,27) and pathogen-host interactions (so-called effectors; Stergiopoulos and de Wit, 2009; Stukenbrock et al., 2011) have been reported in several species.

Fungal plant pathogens pose as excellent models for the study of adaptive evolution, as these organisms are exposed to highly distinct selective pressures. Prominent examples of crop pathogenic fungi have been disseminated across continents with agriculture. They can successfully cause disease across very distant climatic and environmental scales and overcome various control measures, including crop resistance and fungicide applications (40–43. Several species display drastic variation in both Ne and frequency of sexual recombination (32–36. Alternative strategies for adaptive evolution include scattered regions across the genome with a higher accumulation of mutations (6,37). These regions often harbor genes related to environment and host adaptation, which display hallmarks of strong positive selection and relaxed purifying selection (8,9). Another characteristic of fungi that fosters fast adaptation is a larger proportion of accessory genes across the genome (38). At the species level, genes located on accessory regions would be dispensable under unfavorable conditions, decreasing adaptive constraints for the species (38,39).

In this study, we detect and quantify the contribution of adaptive molecular evolution across 20 pairs of fungal species represented by complete genome information from over 2,400 individuals. We aimed to quantify rates of evolution across a variety of different fungal species and to test the widely claimed hypothesis that the rate of adaption is faster in pathogenic species, while elucidating the roles of effective population size, recombination, functional categories and accessory genes on adaptive rates.

A common pitfall of comparing adaptive evolution across taxa stems from methods using varying assumptions for the distribution of the fitness effect of mutations, often over-simplistically (11). Additionally, early methods were strongly affected by the demographic history and impact of population structure, which biased estimations of adaptive evolution. Therefore, it becomes particularly challenging to incorporate extensive population genomic data from global samples without introducing biasing effects due to different population structure and demographic history (3,30). To address this challenge, we employed state-of-the-art methods that account for the distribution of fitness effect (DFE) of new mutations and the confounding factors stemming from demography or population structure (23,14,31). We further used simulations to demonstrate the robustness of our conclusions. By incorporating genetic information from multiple populations of different species, we provide an unprecedented atlas of adaptive molecular evolution in fungi.

## Results

### Extensive variation in patterns of diversity and recombination among fungal species

To establish a comprehensive picture of the rates of protein evolution in fungi, notably species infecting plants, we analyzed 2,478 complete genomes of individuals collected from natural or agricultural environments in North America, Europe, Asia, South America, Africa, and Oceania (Figure 1A). All genome analyses, including read mapping, SNP calling, and annotation were conducted *de novo* with raw sequence reads in order to obtain a high-quality dataset for comparative analyses (see Materials and Methods). We included a total of 20 target fungal species, reflecting a broad taxonomic diversity in the largest fungal phylum (Ascomycota) and representing pathogenic and non-pathogenic species (Figure 1B-C). Among the pathogenic species, we included samples infecting a broad range of host plants, such as economically important crops and cash crops, edible fruits, and ornamental species (Figure 1C). For all target species included, isolates were sampled from hosts or environments in more than one country, incorporating higher genetic variation stemming from adaptation to different hosts, microbial interactions, and climatic conditions. The number of individuals per focal species varied from 18 (*Magnaporthe oryzae* pathotype *Triticum*) to 485 (*Zymoseptoria tritici),* and being above 100 for ten of the species (Figure 1C).

**Figure 1.**
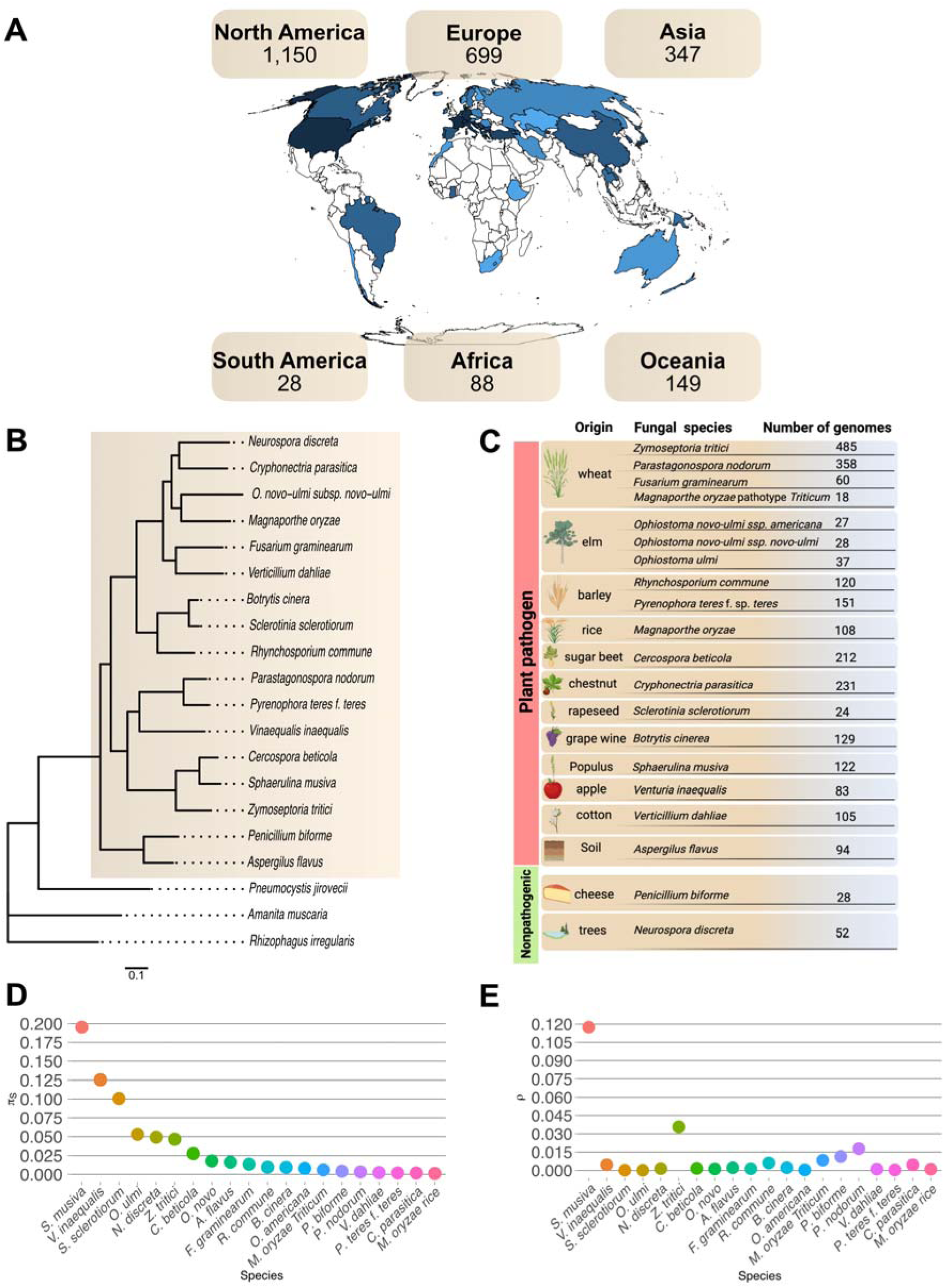
Geographic origin and evolutionary relationship of individual samples and fungal species. (A) World map showing the geographical distribution of the 2,461 isolates. Isolates were obtained from countries highlighted in blue. (B) Maximum-likelihood phylogenomic tree generated from a concatenated set of protein sequences. The 18 species used as established refences are highlighted in the square background. A total of 1,065 single-copy orthologs present in all species was used. The tree was estimated using the PROTGTR model, and was rooted using *Rhizophagus irregularis* as an outgroup. All clades have 100% support (Felsenstein’ bootstrap procedure with 100 re-samples). (C) List of host/environment of origin for each fungal species and total number of genomes per species. The first 18 species represent obligated plant pathogens, while the two last represent fungal species that are not pathogenic to plants or animals. (D) neutral genetic diversity π_S_, and (E) population recombination rate per species, ρ.

We extracted and aligned sequences of protein-coding genes along the genomes to infer rates of adaptive non-synonymous substitutions. First, we focused on core genes and applied a strict filtering scheme to include, for each species, only genes for which we were able to cover complete gene sequences from all individuals. The total number of core protein-coding genes defined in this way varied from 3,307 in *Botrytis cinerea* to 7,727 in *Fusarium graminearum*, with a median of 5,460 genes across the 20 species (Figure 2A, Supplementary Table S5).

**Figure 2.**
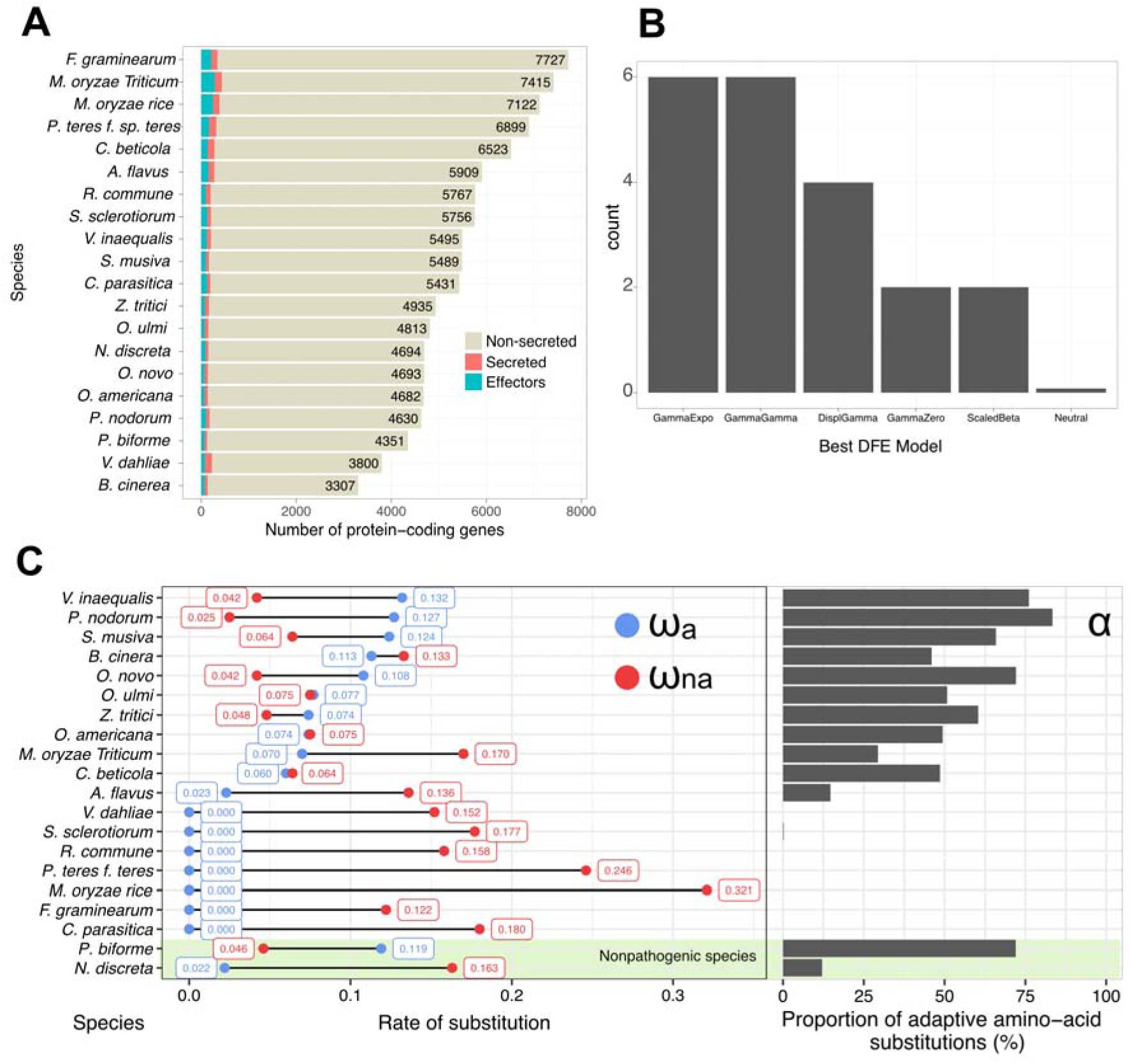
Landscape of adaptive and non-adaptive protein evolution across species. (A) Proportion of non-secreted, secreted and predicted effector proteins per species. (B) Best models for the distribution of fitness effects (DFE) of mutations. (C) Estimations for the adaptive nonsynonymous substitutions rate (ω_A_) and non-adaptive non-synonymous substitutions rate (ω_NA_), in blue and red, respectively. Bar-plots represent α (proportion of amino-acid substitutions that are adaptive). ω_A_, ω_NA_, and α estimations are from the entire core gene set of each species, calculated after model averaging over all models tested. Species are in decrescent order based on ω_A_ within plant pathogen and non-pathogenic categories.

Next, we considered genetic variation across the genes of each species (Supplementary Table S3). Our inference of adaptation relied on intra-specific polymorphism data, from which we built unfolded synonymous and non-synonymous site frequency spectra (SFS), using an outgroup species for each focal species to reconstruct ancestral alleles, and compute non-synonymous (d_N_) and synonymous divergence parameters (d_S_). The resulting SFS, together with the divergence parameters, were used to fit models of distributions of fitness effects with a component of positively selected mutations, allowing us to infer the proportion of adaptive mutations. We found that the median rate of synonymous substitutions per site (d_S_) across all pairs of species was 0.19, ranging between 0.02 in *Magnaporthe oryzae* rice and its outgroup species *M. oryzae* (*Setaria* host) to 0.63 between *Parastagonospora nodorum* and its outgroup species *P. jasniorum* (Supplementary Table 8).

The effective population size and recombination rate are key factors that can influence the potential of a species to respond to natural selection. To address the relevance of these two parameters, we estimated π_S_ reflecting the neutral genetic variation in the population and here considered a relative proxy for the effective population size, and we computed the population recombination rate parameter ρ (Figure 1). Across species, π_S_ varied up to a factor of 162, with the pathogenic species *M. oryzae* rice (π_S_=0.0012) and *S. musiva* (π_S_=0.1951) having the lowest and the highest π_S_, respectively (Figure 1D). The population recombination rate varied across species by a factor up to 2050, with *O. ulmi* (ρ=5.71 × 10^-5^) and *S. musiva* (ρ=1.17 × 10^-1^) showing the lowest and highest values, respectively (Figure 1E). We note that the population recombination rate ρ is the product of the effective population size and the recombination rate per generation per nucleotide r (ρ=2Ner, for haploid organisms) and that the extensive variation in π_S_ and ρ might reflect the interplay of different evolutionary forces across species. Hence, acquiring quantitative measures of the adaptive and non-adaptive rate of protein evolution is crucial to understanding the contribution of natural selection and neutral evolution in shaping a species’ evolutionary history.

### Extensive variation in rates of adaptive evolution among pathogenic and non-pathogenic fungal species

New mutations will segregate in a population with neutral, advantageous or slightly deleterious effects on the fitness of the organisms. To account for different fitness effects of polymorphisms and substitutions, we inferred the distribution of fitness effects (DFE) of mutations across species using six models (Figure 2B; Supplementary table S9). These included Neutral, GammaZero, GammaGamma, GammaExpo, DisplGamma, and ScaledBeta. We used Akaike’s Information Criterion (AIC) to select the best model per species. The Neutral model was not favored in any species. Instead, among the 20 species, we found that the DFE is best explained by the GammaGamma and GammaExpo models for twelve species in total (Figure 2B; Supplementary table S10). Both models assume the segregation of mutations with a weakly negative effect on fitness modeled as a (negative) Gamma distribution. However, the former assumes a Gamma model incorporating mutations with weakly advantageous effects, while the latter assumes an exponential distribution. Other models that include the distribution of both mutations with positive and negative effects were favored among the remaining six species, such as DisplGamma and ScaledBeta. Finally, the GammaZero model was selected in two species, indicating the predominance of mutations with slightly negative effects on fitness (Figure 2B).

We next sought to quantify and unveil the patterns of adaptive evolution at a genome-wide scale. We calculated the adaptive nonsynonymous substitutions rate (ω_A_), non-adaptive non-synonymous substitutions rate (ω_NA_) and the proportion of adaptive amino-acid substitutions (α) based on genome-wide alignments of core genes (Figure 2B). We first found striking differences in the signatures of adaptive evolution across fungal species. α ranges between 0 for several species to 83% for *P. nodorum* (Figure 2C). Seven species had a α value above 50%, indicating a predominance of fixed adaptive substitutions that contribute to protein evolution compared to non-adaptive substitutions. Concordantly, we also observe considerable differences in rates of adaptive and non-adaptive evolution. For ω_A,_ we observed a variation between 0 (in several species) and 0.132 (*V. inaequalis*), while ω_NA_ varied between 0.025 (in *P. nodorum*) and 0.321 (in *M. oryzae* Rice) (Figure 2C; Supplementary table S9). Interestingly, *P. biforme*, a non-pathogenic species, featured relatively high levels of ω_A_ and α as compared to plant pathogenic species. This suggests that rapid adaptative evolution is not a unique hallmark of pathogenic fungi but also occurs in non-pathogenic species. Overall, our findings indicate that the variation in α is driven by a combination of higher ω_A_ and lower ω_NA_, allowing us to further disentangle the effects of adaptive and non-adaptive evolution at the genome-wide scale.

### Model-based estimations of **ω**_NA_, **ω**_A_ and **α** are robust to the influence of demography and population structure

Next, we address the influence of demography and population structure on our estimates of adaptive evolution. Hereby, we demonstrate that our estimations of ω_NA_, ω_A_, and α are robust to the effects of the species’ demographic history and population structure when using a model-based approach. Under the basic and FWW models (see Materials and Methods), we find that α is underestimated in all scenarios, with an increased underestimation when the proportion of positively selected mutations (*p*) is lower (Supplementary material). A similar pattern is observed for ω_A_, which is underestimated across all scenarios for both basic and FWW models, but the underestimation increases with an increase in *p*. For the last estimator, ω_NA_, we observed a consistent overestimation under all scenarios, which increases under higher *p*. The trend of higher ω_NA,_ when *p* is higher, is likely due to the linkage effect between more deleterious mutations and adaptive mutations that are fixed. Indeed, in our simulations, we used low recombination rates and short intergenic regions. On the other hand, the model-based inferences using GammaExpo model are highly robust to demography and population structure (Supplementary material). Notably, the model-based inference reports a lower range for ω_A_ and α in cases of strong departure of a panmictic, constant population, leading to conservative estimates. Overall, the model-based inference results in more accurate estimates of ω_NA_, ω_A_, and α across both panmictic and non-panmictic population dynamics.

### A larger effective population size reduces the fixation rate of slightly deleterious mutations in fungi

To expand our understanding of factors globally affecting the variation in adaptive protein evolution across fungi, we first correlated our genome-wide estimates of adaptive evolution with two parameters that we consider as proxies for the effective population size (π_S_)=2.Ne.u (where u is the genome average mutation rate) and the genome average recombination rate (ρ=2.Ne.r, where r is the genome average recombination rate) computed for the different fungal species. Theory predicts that if adaptive evolution is limited by the supply of new mutations, species with a larger effective population size will have a higher rate of adaptive evolution and a stronger efficacy of purifying selection (23). Moreover, in addition to population size also recombination matters for the fixation of beneficial mutations and the removal of deleterious ones as frequent sexual recombination will mitigate the effects of linked selection (17). Hence, we hypothesize that species with larger effective population size and frequent sexual recombination will show higher rates of adaptive evolution while maintaining lower rates of non-adaptive substitutions.

Our estimates of adaptive evolution across a diversity of fungal species allow us to correlate rates of adaptation with both the effective population size and the recombination rate. We observed a positive correlation between log(π_S_) and *a* = *ω_A_*⁄(*ω_A_* + *ω_NA_*), and a negative correlation between log(π_S_) and dN/dS (*ω* = *ω_A_* + *ω_NA_*) (Figure 3). This is explained by distinct effect on the rate of adaptive and non-adaptive non-synonymous, log(π_S_) is positively correlated with ω_A_ and negatively correlated with ω_NA_. Similar patterns were observed for the correlations using ρ. A positive trend was observed between ρ and both α and ω_A_, while a negative correlation was found between ρ and both ω and ω_NA_. These correlations indicate that more adaptive substitutions are fixed in species with either higher effective population sizes or higher recombination rates, while the action of purifying selection more efficiently removes deleterious substitutions.

**Figure 3.**
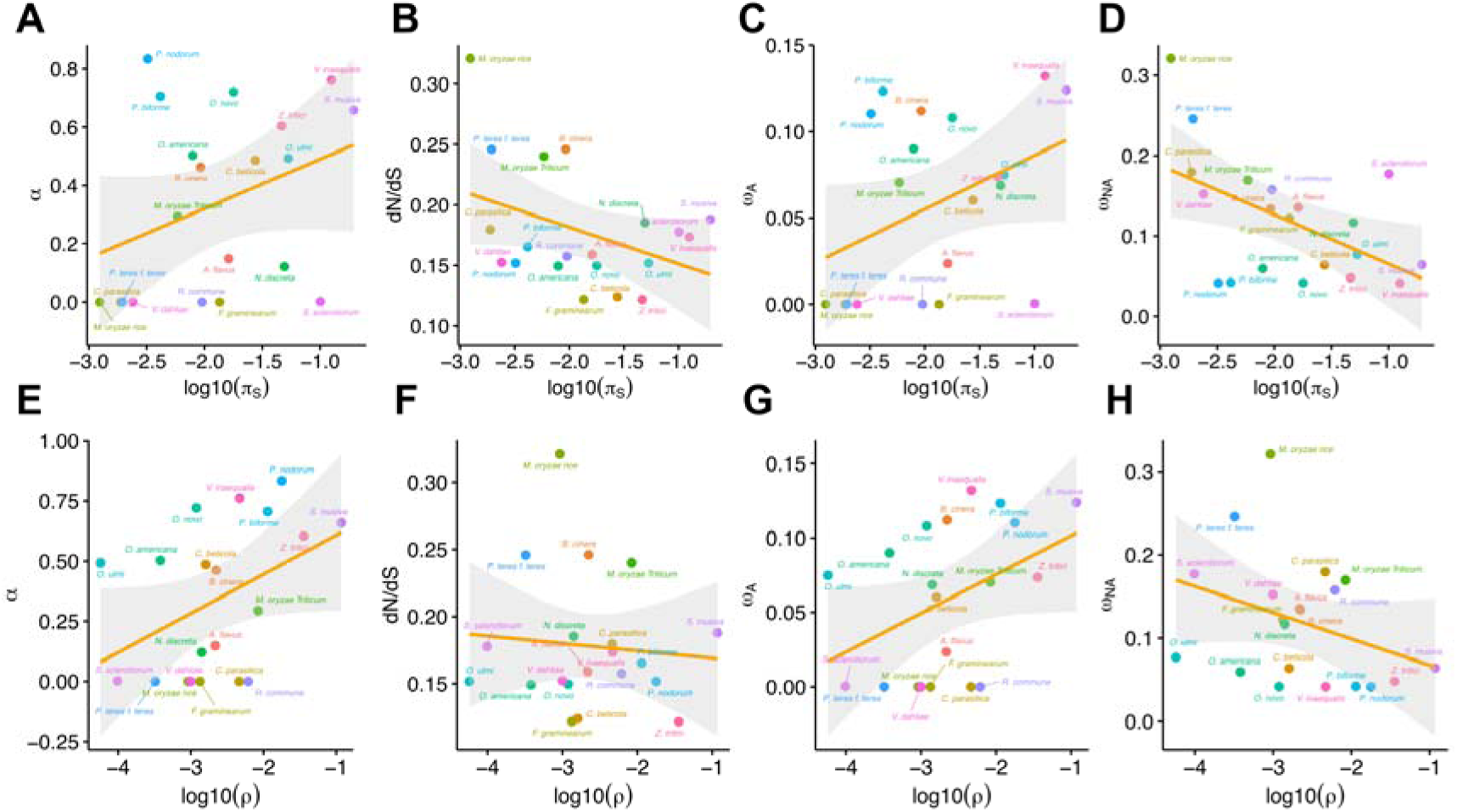
Species wide correlations between neutral genetic variation (π_S_) (A-D) and population recombination rate (ρ) (E-H) with ω_NA_, ω_A_, and α and dN/dS. Log transformation was applied to π_S_ and ρ. Each dot represents one of 20 target species. The significance of correlations is not shown, as it was determined using a generalized least squares method as described in the main text.

We used a phylogenetic generalized least square (PGLS) approach (44) to assess the statistical significance of the observed correlations across the 20 species. As variation in α can be driven by both variations in ω_A_ and ω_NA_, we focused our comparisons on these two rates to disentangle adaptive and non-adaptive evolution. We fitted two distinct models, using either ω_A_ or ω_NA_ as response variables, and recombination (as log(p)) and population size (as log(π_S_)) as explanatory variables. These PGLS analyses revealed a similar positive trend of the two variables, yet only marginally significant (*P* = 0.068 and *P* = 0.083, respectively, Table 1). When considering the non-adaptive rate of evolution (ω_NA_), we found a marginally significant negative correlation with recombination (*P* = 0.059), but a significant negative correlation with population size (*P* = 0.029, Table 1). Our results corroborate the importance of effective population size in shaping adaptive evolution in fungi. Additionally, our study also highlights the reduced influence of the accumulation of non-adaptive non-synonymous substitutions in species with larger effective population size. A similar observation was described in a comparative study of adaptive evolution in 44 vertebrate species. Across the large number of vertebrate species, effective population size was negatively correlated with ω_NA_ (14), indicating less fixation of slightly deleterious mutations in species with larger effective population size. Furthermore, due to the impact of recombination on both ω_A_ and ω_NA_ we postulate that recombination indeed acts on adaptive evolution by alleviating the effects of linked selection. However, the effect of recombination estimated in the linear models is only marginally significant (PGLS models 1 and 2). This weak effect may result from intra-genomic variation of substitution rates, which may vary in orders of magnitude similar to between-species variation (11). In the following, we further investigate the intra-genomic distribution of substitution rates.

**Table 1.**
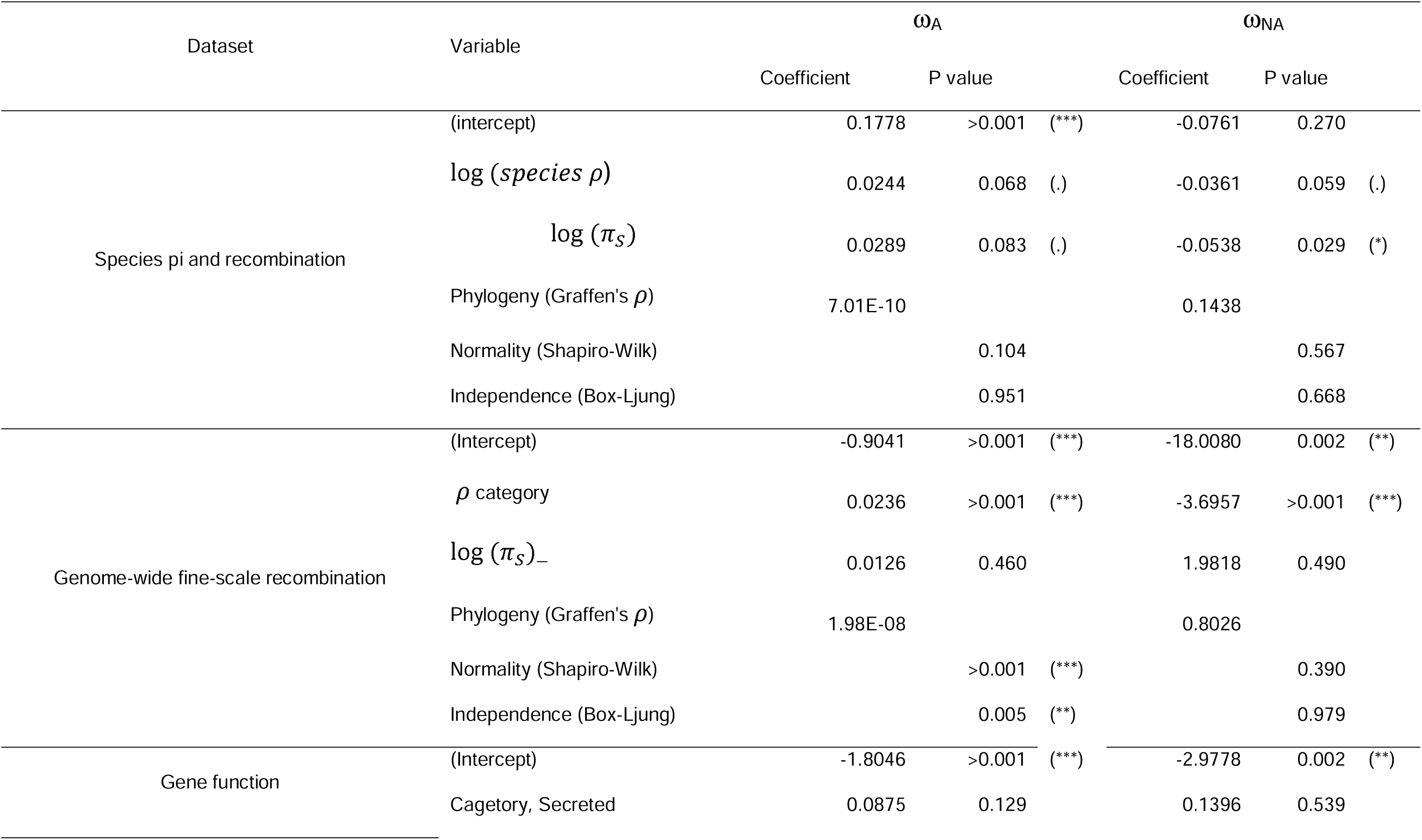
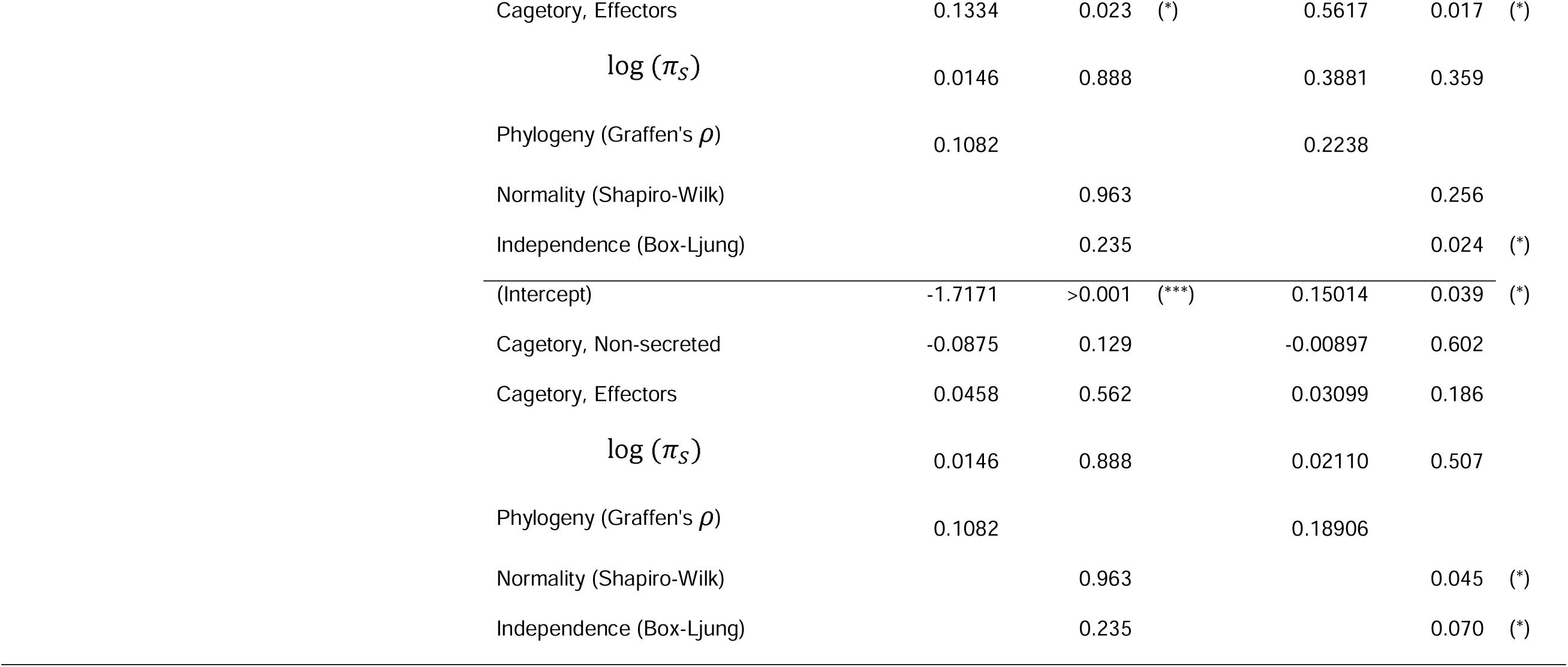
Generalized least squares (GLS) linear modeling to assess the effects of recombination p, effective population size (π_S_), and gene function across species.

| Dataset | Variable | $\omega_A$ | | | $\omega_{NA}$ | | |
| --- | --- | --- | --- | --- | --- | --- | --- |
|  |  | Coefficient | P value |  | Coefficient | P value |  |
| Species pi and recombination | (intercept) | 0.1778 | >0.001 | (***) | -0.0761 | 0.270 |  |
| | $\log(\text{species } \rho)$ | 0.0244 | 0.068 | (.) | -0.0361 | 0.059 | (.) |
| | $\log(\pi_S)$ | 0.0289 | 0.083 | (.) | -0.0538 | 0.029 | (*) |
| | Phylogeny (Graffen's $\rho$) | 7.01E-10 | | | 0.1438 | | |
|  | Normality (Shapiro-Wilk) |  | 0.104 |  |  | 0.567 |  |
|  | Independence (Box-Ljung) |  | 0.951 |  |  | 0.668 |  |
| Genome-wide fine-scale recombination | (Intercept) | -0.9041 | >0.001 | (***) | -18.0080 | 0.002 | (**) |
| | $\rho$ category | 0.0236 | >0.001 | (***) | -3.6957 | >0.001 | (***) |
| | $\log(\pi_S)_-$ | 0.0126 | 0.460 | | 1.9818 | 0.490 | |
| | Phylogeny (Graffen's $\rho$) | 1.98E-08 | | | 0.8026 | | |
|  | Normality (Shapiro-Wilk) |  | >0.001 | (***) |  | 0.390 |  |
|  | Independence (Box-Ljung) |  | 0.005 | (**) |  | 0.979 |  |
| Gene function | (Intercept) | -1.8046 | >0.001 | (***) | -2.9778 | 0.002 | (**) |
|  | Cagatory, Secreted | 0.0875 | 0.129 |  | 0.1396 | 0.539 |  |

|  |  |  |  |  |
| --- | --- | --- | --- | --- |
| Cagetory, Effectors | 0.1334 | 0.023 (*) | 0.5617 | 0.017 (*) |
| log ($\pi_s$) | 0.0146 | 0.888 | 0.3881 | 0.359 |
| Phylogeny (Graffen's $\rho$) | 0.1082 | | 0.2238 | |
| Normality (Shapiro-Wilk) |  | 0.963 |  | 0.256 |
| Independence (Box-Ljung) |  | 0.235 |  | 0.024 (*) |
| (Intercept) | -1.7171 | >0.001 (***) | 0.15014 | 0.039 (*) |
| Cagetory, Non-secreted | -0.0875 | 0.129 | -0.00897 | 0.602 |
| Cagetory, Effectors | 0.0458 | 0.562 | 0.03099 | 0.186 |
| log ($\pi_s$) | 0.0146 | 0.888 | 0.02110 | 0.507 |
| Phylogeny (Graffen's $\rho$) | 0.1082 | | 0.18906 | |
| Normality (Shapiro-Wilk) |  | 0.963 |  | 0.045 (*) |
| Independence (Box-Ljung) |  | 0.235 |  | 0.070 (*) |

### Fine-scale recombination landscape reveals the strong impact of recombination on adaptive and non-adaptive rates of evolution

To further explore the effects of recombination on the rates of adaptation we classified genes according to the average recombination rate in their genomic region. We then compared estimates of ω_A_ and ω_NA_ for each “recombination rate category”. This fine-scale method revealed two contrasting species profiles of genome-wide recombination: (i) species with broader quantitative variation in recombination across the genome (i.e., *S*. *musiva*) and (ii) species with lower variation in genome-wide recombination (i.e., *C. parasitica*; (Supplementary figure 1 and 2).

To account for the heterogenous recombination landscape, we down-sampled our dataset to a subset of six species belonging to the first recombination profile. This down-sampling enabled us to obtain a more robust assessment of the effects of fine-scale recombination on both ω_A_ and ω_NA_ for each p category (Supplementary figures 3 and 4). First, we found a positive correlation between p and ω_A_ across genomic regions for all six species, which was highly significant for five of them (Supplementary figure 3). The opposite pattern was observed with ω_NA_, where we find a negative correlation with the recombination rate (Supplementary figure 4).

Next, we performed GLS modeling using the same estimations of ω_A_ and ω_NA_ across fine-scale recombination categories and included the log(π_S_) of each species as a co-factor. The models confirmed the pervasive effect of recombination across the genome, where genes in regions with more recombination had significantly higher rates of adaptive evolution and lower rates of non-adaptive evolution (*P* < 0.001 in both models, Table 1). These results confirm the previously postulated effect of recombination on adaptive evolution via its effect on linked selection. However, it also indicates that recombination is effective only when the sexual cycle is recurrent in the species. The subset of six species selected displays evident levels of recombination across the genome, while predominantly asexual species were not included due to estimations of ω_A_ being zero (second recombination profile previously described).

### Increased accumulation of adaptive protein changes in genes encoding effector proteins

We next addressed the impact of gene function on the rates of adaptation. We hypothesized that the rate of adaptation occurs faster in genes encoding secreted proteins, as these are exposed to the very different environments inhabited by the fungi studied here. Moreover, of the secreted proteins, we considered “effectors” to be the proteins with the highest rate of adaptation, as these are predicted to act in host-pathogen interactions.

We predicted proteins targeted for secretion in all species, and our pipeline furthermore also identified effector proteins in all species, including the two non-pathogenic fungi. Effector proteins are identified by criteria including protein length and the composition of amino acids hereunder the cysteine content. Non-pathogenic fungi also produce and secrete proteins with these characteristics, suggesting that these proteins have a broader function beyond plant-pathogen interactions.

The number of secreted proteins ranged from 434 in *M. oryzae Triticum* to 124 in *P. biforme*, while the number of effectors varied from 281 in *M. oryzae Triticum* to 65 in *P. biforme* (Supplementary table S4). Comparing the total number of secreted proteins and effectors among fungal species, we find slightly more genes encoding secreted proteins and effectors in the pathogenic species relative to the non-pathogenic species. However, as we only included two non-pathogenic species in our analysis, the comparison does not allow us to make general conclusions about gene content in pathogenic and non-pathogenic fungal species. In summary, we find a median of 203 and 137 secreted proteins in the pathogenic and non-pathogenic species, respectively, and a median of 108 and 78 effectors in the pathogenic and non-pathogenic species, respectively (Supplementary table S3).

Several studies have demonstrated a high genetic variability in genes encoding predicted effectors (45,8,9). While this variation might reflect positive selection of new mutations, it might also reflect a relaxation of purifying selection. To disentangle these two scenarios, we compared both ω_A_ and ω_NA_ among gene categories. When we compare the rate of adaptive evolution in effector proteins and non-secreted proteins, we find a significantly higher value of ω_A_ in effector proteins in eight out of 14 species. The two non-pathogenic species *N. discrete* and *P. biforme* also exhibit a higher rate of adaptation in genes predicted to encode effector proteins (Figure 4; ω_A_ for all species in supplementary figure 5; Table S11). Comparing non-secreted and secreted proteins, we find that ω_A_ is significantly higher for genes encoding secreted proteins in only five species, all pathogenic (Figure 4; ω_A_ for all species in supplementary figure 5; Table S11). When we compare only genes encoding secreted proteins and effectors, ω_A_ is significantly higher in effector proteins in four species (*V. inaequalis*, *S. musiva*, *O. ulmi*, and *S. sclerotiorum*), while secreted proteins have higher ω_A_ than effectors in only a single species (*O. novo*), all pathogenic (Figure 4; ω_A_ for all species in supplementary figure 5; Table S11).

**Figure 4.**
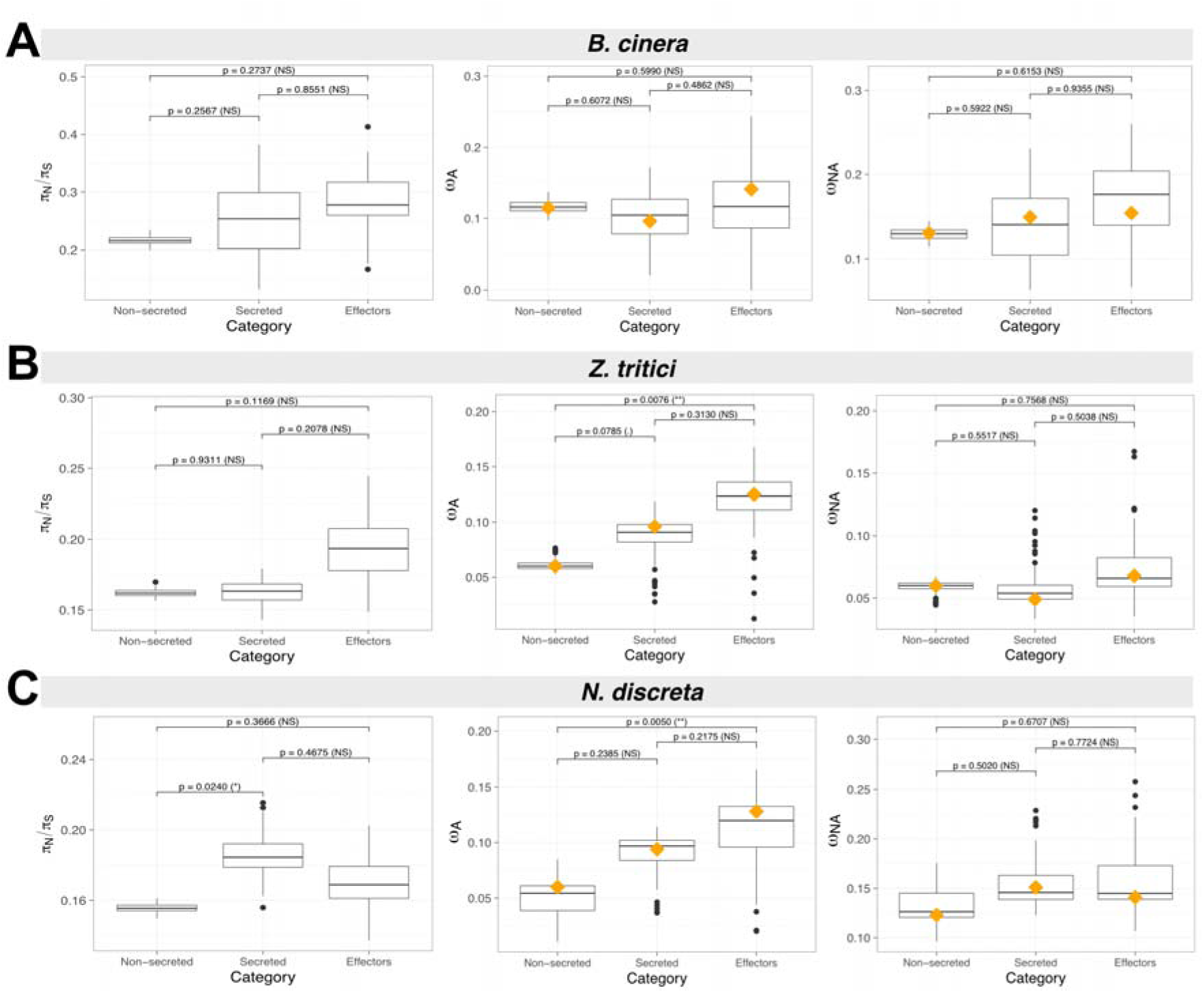
Three landscapes of the strength of purifying selection (π_S_/π_N_) and rates of adaptation (ω_A_ and ω_NA_) among non-secreted, secreted and effector gene categories. (A) *Botrytis cinerea,* (B) *Zymoseptoria tritici* and (C) *Neurospora discreta*. Box-plot distribution was generated using 100 bootstrap replicates performed using the best model for each species according to Akaike’s information criterion. The orange diamond represents the mean value calculated after averaging over all models of distribution of fitness effects. The significance of differences was determined based on 1000 permutations. Complete panel for all 20 species in the supplementary figures 6 and 7.

Intriguingly, for the parameter ω_NA_, expressing the rate of fixation of non-adaptive non-synonymous mutations, in general was slightly higher in genes encoding secreted proteins and effectors although in most comparison not significantly higher (Figure 4; ω_NA_ for all species in supplementary figure 6; Table S11). Together, the results show that genes encoding non-secreted proteins are generally more conserved than genes encoding secreted proteins and effectors. Moreover, we also show that effector proteins typically have higher rates of adaptation.

We further tested these observations using a linear model. Hereby we obtain further support that effectors generally have significantly higher ω_A_ and ω_NA_ than non-secreted proteins across species (*P* = 0.023 and *P =* 0.017, respectively; Table 1). However, values of ω_A_ and ω_NA_ in effector proteins are not significantly different from those in secreted proteins. We also observed a higher standard deviation in ω_A_ and ω_NA_ for secreted and effector proteins compared to non-secreted proteins (Figure 4 and supp figures 5, 6 and 7). This pattern likely reflects a higher level of conservation in housekeeping genes such as essential enzymes, while other proteins secreted to interact with the host or environment evolve at faster rates. Both median π_N_/π_S_ and d_N_/d_S_ support a less stringent level of purifying selection in effector proteins than non-secreted proteins in most species (Supplementary figures 8, 9, 10, 11, 12, 13). Overall, we show that effector proteins have higher rates of mutations fixed by positive selection. However, a relaxation of purifying selection also contributes to maintaining higher genetic variation in these genes.

**Figure 5.**
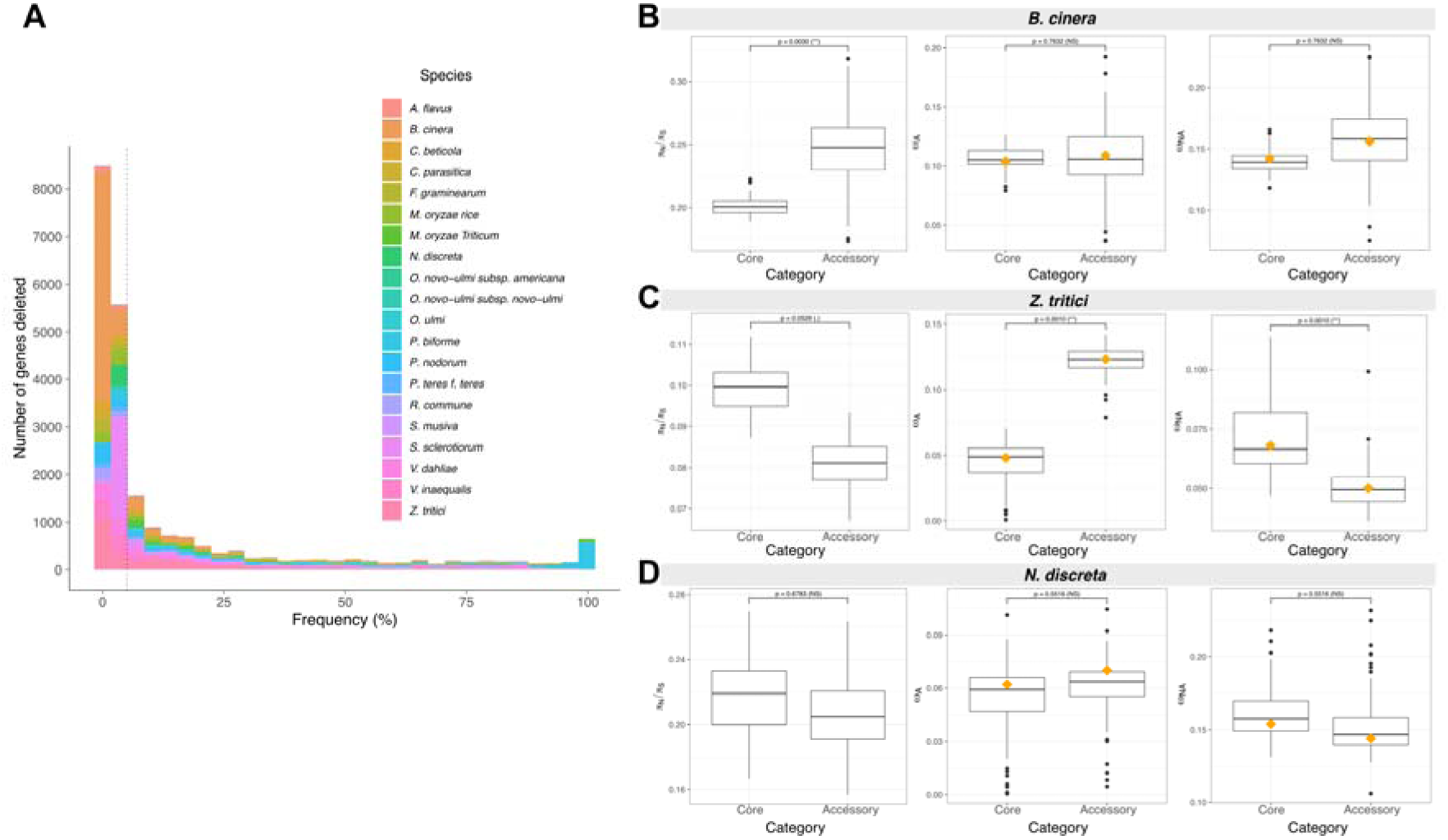
Variation of the strength of purifying selection (π_S_/π_N_) and rates of adaptation (ω_A_ and ω_NA_) between core and accessory gene datasets. (A) Number of gene deletions across isolates. The grey dotted line represents the 5% threshold used as maximum deletion frequency for the accessory dataset. (B) *Botrytis cinerea* with 3,524 genes and 47 isolates, (C) *Zymoseptoria tritici* with 503 genes and 36 isolates, and (D) *Neurospora discreta* with 128 genes and 22 isolates. The box-plot distribution was generated using 100 bootstrap replicates performed using the best model according to Akaike’s information criterion. The orange diamond represents the mean value calculated after averaging over all models of distribution of fitness effects. The significance of differences was determined based on 1000 permutations. Complete panel for 14 species in the supplementary figures 23 and 24.

### Accessory protein-coding genes accumulate higher adaptive changes

Genes present in accessory regions have been shown to play a role in pathogen adaptation in several microbial species (38,39). In some species, these regions encode genes involved in virulence and host specificity. Accessory regions are often found in specific compartments enriched with transposable elements and with a faster rate of evolution (37). Mutations can, to a larger extent, accumulate in these regions without disrupting essential metabolic functions encoded by “core compartments” of the genome. We used the population genomic datasets to identify accessory regions and genes across the twenty fungal species. Hereby, we found an extensive variation in the number of accessory genes, ranging from 67 in *O. novo-ulmi subsp. americana* to 6,570 in *B. cinera* (Figure 5A; Supplementary table S12). The frequency of gene deletions per species was skewed towards lower frequencies, indicating that genes are generally missing in a small portion of isolates, and widespread deletions are rare (Supplementary Figure 14).

We next determined the contribution of adaptive and non-adaptive substitutions to the evolution of core and accessory protein-coding genes. We used datasets of accessory genes missing in up to 5% of isolates per species (Figure 5A; Supplementary table S13). The proportion of adaptive amino-acid substitutions varied among species, being higher for seven out of nine species for which the analysis was successfully performed (Supplementary Figure 15). When disentangling the adaptive portion of ω, we found a predominance of higher ω_A_ across the same previous nine species, with significant statistical differences in five species (Figure 5; Supplementary Figure 16). The higher ω_A_ found in accessory genes indicates a strong contribution of positive selection among species. On the other hand, the non-adaptive non-synonymous substitutions rate (ω_NA_) was higher in the core genes for five species out of a total of nine that could be analysed (Figure 5; Supplementary Figure 17). ω_NA_ indicates that the core gene portion of *P. nodorum* and *S. sclerotiorum* is subject to higher selective constraints, while in *Z. tritici*, *C. beticola*, *A. flavus* there is an opposite trend. Additionally, the results from π_N_/π_S_ comparisons indicate that purifying selection is less stringent on accessory genes in only a few species (i.e., *S. sclerotiorum*; Supplementary figure 18,19,20,21,22,23). Together, the observed patterns of ω_A_ and ω_NA_ indicate that genes located in accessory regions are subject to positive selection and adapt at higher rates than core genes. Differences in the patterns of genetic variation within and among species however, indicate that the impact of natural selection on accessory gene evolution vary substantially.

## Discussion

Empirical data of animal and plant model species has been widely explored to test general principles in population genetics (2,46). How eukaryotic microorganisms with highly distinct life history traits adhere to prevailing models of population genetics has so far only been addressed for individual species (e.g., 18,9,46) but not across large evolutionary scales. The growing amount of population genetic data now allows us to quantify and compare rates of evolution across diverse microbial species and thereby test general hypotheses related to adaptive evolution. In our study, we conducted a comparative analysis of 20 fungal species, representing a diverse panel of ascomycete lineages. We demonstrate extensive variation in rates of adaptive evolution, hereby underlining that rapid adaptation is not a general premise of pathogenic lifestyles. We highlight the important contribution of factors such as effective population size, recombination rate, and gene function in shaping molecular evolution in fungi. Using simulations, we demonstrate the robustness of our estimations to confounding effects from demography and population structure.

### Effective population size and recombination are major determinants of adaptive and non-adaptive evolution in fungi

Many tests of positive selection consider that all substitutions occurring at non-synonymous sites in protein-coding genes are adaptive. However, non-synonymous substitutions with neutral or slightly deleterious fitness effects can also be fixed by chance events, genetic drift, or linked selection (47). In our study, we explicitly distinguished ω_A_ and ω_NA_ and thereby show a pervasive effect of non-adaptive substitutions on fungal evolution as indicated by α below 0.5 and ω_A_ < ω_NA_ for several species. Despite the occurrence of non-adaptive evolution and slightly deleterious mutations segregating within species, we also identified a subset of species where adaptive evolution is predominant. In this subset, comprising both pathogenic and non-pathogenic species (Figure 2C), over 50% of amino-acid differences were fixed by positive selection, and ω_A_ was higher than ω_NA_.

Estimates of adaptive evolution across a large panel of plant species, revealed that plants overall undergo little adaptive evolution in protein-coding genes. Most of the genetic variants which are fixed at non-synonymous sites are either neutral or slightly deleterious (48,23). The low rates of adaptation in plants have been attributed to low effective population sizes although some species with larger effective population sizes also showed little evidence of adaptation. However, there is a negative correlation between π_N_/π_S_ and effective population size in plants implying that the efficacy of purifying selection against slightly deleterious mutations is determined by the effective population size; a correlation also observed in our study of fungi (Figure 3C). Contrary to observations in plants, adaptive processes appear to dominate in vertebrate animal species (14). Interestingly, the relevance of effective population size in adaptation rates was not evident across 44 animal species.

Our data provide evidence that the effective population size plays a determining role in fungal evolution, both with respect to adaptive and non-adaptive evolution. We observe a positive correlation of πS (considered here as a proxy for the effective population size) and ω_A_ supporting that adaptation occurs faster in fungal species with large effective positive sizes. Moreover, we show a negative correlation of π_S_ and ω_NA_ consistent with a stronger efficacy of purifying selection, and thereby more efficient removal of non-adaptive mutations, in species with larger effective population sizes. Intriguingly, across the 20 fungal species, we find highly different values of π_S_ suggesting a large variation in effective population sizes. We speculate that variation in effective population sizes reflects different species histories as well as different reproduction strategies ranging from highly outcrossing to clonal.

In addition to the effective population size, we show that also the recombination rate is a determining factor in adaptation. We show this by correlating genome-wide estimates of recombination with ω_A_ and ω_NA_. The positive correlation of ρ and ω_A_ is consistent with the expected role of recombination in accelerating adaptive evolution. Essentially, recombination reduces the amount of Hill–Robertson interference on linked mutations and thereby facilitates the fixation of adaptive variants (24). Likewise, the removal of slightly deleterious mutations is facilitated by recombination. Recombination has been reported in plants (18), insects (49), bacteria (19), and individual species of fungi (8) to indeed mitigate the effects of linked selection.

Among the 20 fungal species studied here, we find large variability in the recombination rate parameter ρ. This is not surprising given the variability in reproductive mode among species. Contemporary populations associated with crops like *Verticilium dahliae* and *Botrytis cinera* are considered to be predominantly clonal (50,51), while species like *Zymoseptoria tritici* and *Parastagonospora nodorum* undergo extensive sexual reproduction (34,42). Comparative genome studies and population genomic analyses have been used to demonstrate that clonal pathogenic fungi can also adapt to changing and constraining conditions (e.g. in presence of antifungals) by the acquisition of structural variants (52), underlining the relevance of other types of adaptive changes in these organisms.

### Function and location in accessory regions are major contributors of genes molecular evolution in fungi

Several fungal species have been shown to comprise highly dynamic regions across the genome that accumulate mutations at higher rates. These regions often harbor genes related to host interaction and genes segregating as presence absence variation within species (so called accessory genes) (53–55,37). Often genes encoding virulence determinants, so called effectors, are enriched in these variable genomic compartments.

Across the 20 fungal species that we studied, we find a prevalent pattern where genes predicted to encode effectors show increased rates of adaptive evolution. However, in several species the accumulation of non-adaptive substitutions measured as ω_NA_ is also increased in these genes. This observation may reflect a lower efficacy of purifying selection in evolution of these genes. A general assumption is that genetic variation per se is favored in effector-encoding genes to facilitate arms-race evolution with the host (28,54). Interestingly, the two non-pathogenic species also exhibit higher rates of adaptation in genes predicted to encode effector proteins. In this case, we hypothesize that these small secreted proteins have a functional relevance in the interaction of the fungi with other microorganisms. The fact that these also exhibit high rates of adaptation may suggest that not only host-fungus interactions, but also microbial interactions are drivers of rapid adaptation in fungi (56).

In some cases, we note that ω_A_ is higher in genes encoding non-secreted proteins, indicating a high degree of conservation of effector proteins. One of these species is the wheat blast fungus *M. oryzae Triticum*. This species emerged as a new pathogen of wheat very recently. The overall low signature of adaptation in this species likely reflects a bottleneck associated with the recent speciation of the pathogen (within the past few decades). *M. oryzae Triticum* recently emerged as a pathogen of wheat in South America through a host jump from a pasture grass (57,58). We speculate that pathogen emergence via host jumps entails loss of diversity in the founding population, and consider that the lower rate of adaptive substitutions in *M. oryzae Triticum* reflects its recent speciation history.

Genome studies across diverse fungal species have demonstrated extensive variation in genome composition. Hereby fungi exhibit an impressive tolerance regarding presence-absence variation and chromosome rearrangements (52,59). Accessory genes are often considered to have functional relevance in host-pathogen interactions (60,41,37). In the plant pathogenic species *Fusarium oxysporum* and *Alternaria alternata*, genes encoding virulence determinants locate on specific accessory chromosomes or lineage-specific genome compartments (61,62). Hereby the presence or absence of a particular gene determines the ability to infect a particular host species.

We conducted a detailed comparison of the accessory gene repertoires across the 20 fungal species and provide for the first-time quantitative and comparable measures of adaptive evolution in protein-coding genes across accessory and core genome compartments. We observe that the frequency of gene deletions is skewed toward lower frequencies in most species, suggesting that most accessory genes are present in the majority of isolates (Figure 5A). This pattern is similar to the observed across different populations of the fungal plant pathogen *Z. tritici* (38). We demonstrate that accessory genes display a substitution pattern similar to that of effector-encoding genes, with a generally higher rate of both adaptive and non-adaptive non-synonymous substitutions, reflecting extensive positive selection and lower efficacy of purifying selection. Owing to their presence in only a fraction of individuals, accessory genes have a lower effective population size than core genes present in all individuals. Combined with a potentially lower recombination rate, these effects may hinder the impact of purifying selection on accessory genes. We note that the comparison of core and accessory genes included exclusively genes encoding non-secreted and non-effector genes; the measured effect is then independent of the possible additional effect of the gene category (10,8,18). The impact of gene “essentiality” on protein evolution was also investigated by comparing d_N_/d_S_ ratios in other organisms, including mice and yeast (63–65. In these previous studies, genes encoding dispensable proteins were found to evolve at faster rates, congruent with our findings for most fungal species.

To conclude, we have provided extensive analyses of protein evolution across 20 fungal species representing a broad diversity of ascomycete taxa. In general, we find a prevailing effect of adaptation on fungal evolution. High rates of adaptive protein evolution correlate with high effective population sizes, a relationship that is expected if the rate of adaptation is limited by the supply of new beneficial mutations. Pathogens may undergo recurrent population bottlenecks in response to fluctuating host densities, and we speculate that such demographic events explain the considerable variation in effective population sizes among the species investigated here. At a local scale, a primary determinant of adaptive evolution is recombination. Hereby variation in recombination rate correlates with rates of adaptation as expected under Hill-Robertson interference. Fungi may serve as excellent model systems to test population genetics principles including the effects of past and recent demographic events as well as the relative impact of asexual and sexual reproduction on species evolution.

## Materials and Methods

### Whole-genome sequence reads and quality filtering

We compiled a dataset of 2,478 whole-genome sequence reads comprising population genomic information from 37 haploid fungal species from repositories described in Supplementary Table 1. Among the 37 species, 20 are herein referred as focal species, and 17 herein referred to as outgroup species Supplementary Table 1. We submitted Illumina pair-end raw reads to the same filtering procedure using Trimmomatic (v0.35) to remove adaptor sequences and low-quality reads with the following filter settings: ILLUMINACLIP:TruSeq3-PE-2.fa:2:30:10, leading = 10, trailing = 10, sliding-window = 5:10 and minlen = 50 (66).

### Reads mapping and SNP calling for focal species

For each of the 20 focal species, we mapped the trimmed reads on the pre-established reference genomes listed in Supplementary Table 2. We used bowtie2 (v2.4.5) with the --very-sensitive-local option to map the reads (67). The generated SAM files were sorted and converted into BAM files using samtools (v1.9) (68). SNP calling was performed following the Genome Analysis Toolkit (GATK) pipeline (69). First, we used HaplotypeCaller to generate genome VCF files with -ploidy set to 1 and --emit-ref-confidence GVCF. Next, genome VCF files were combined using CombineGVCFs. The combined gVCF file was then genotyped using GenotypeGVCFs with option --max-alternate-alleles 2, and variants selected using SelectVariants with option --select-type-to-include SNP. Finally, SNPs were removed if flagged with any of the following filter criteria: QD < 5; QUAL < 1000; MQ < 20; - 2.0 > ReadPosRankSum > 2.0; -2.0 > MQRankSum > 2.0; -2.0 BaseQRankSum > 2.0. We used vcftools (v0.1.14) to remove SNPs failing the latter filters with -- remove-filtered-all (70).

### Genome assembly, gene predictions, and functional characterization for outgroup species

For each of the 17 outgroup species, we performed de novo assemblies using previously trimmed reads. We used spades assembly toolkit (v3.14.1) with option -- careful to reduce the number of mismatches and short indels (71). We used a range of pre-determined k-mer sizes per outgroup species according to the read length obtained. The quality of each genome assembly was assessed with quast (v4.6.3) (72).

Annotations of protein coding genes were performed for each of the previously assembled 17 outgroup species. Since transcriptome information is unavailable for all outgroup species included in the present study, we used a common approach based on a large protein database with BRAKER2 (v2.1.6) --epmode pipeline using the option --fungus (73,74). Briefly, we downloaded a fungal protein database from OrthoDB (75) and added additional protein sequences from the respective focal species. This merged protein database was used for training Augustus (v3.4.0), which generates protein hints of introns, start, stop, and coding sequence parts. After gene training, Augustus was used to annotate gene predictions in all outgroup species genomes with --alternatives-from-evidence=false.

### Gene orthology identification and gene alignment

After hard filtering, we kept only bi-allelic SNPs without missing information across isolates of each focal species Supplementary Table 3. A consensus fasta file with SNP variables was extracted from each isolate using the consensus function from bcftools (v1.3.1) (76). We used the program MafFilter (77) to extract the coding DNA sequence (CDS) regions from the consensus fasta file of each isolate. CDS coordinates were obtained from the gff file, and parts of the same gene were concatenated using a custom python script (https://github.com/DaniloASP/RatesOfAdaptation). Gene orthology relationships were established using protein sequences for each pair of species with OrthoFinder (v2.5.2), and only single-copy orthologues were selected. Orthogroups were aligned using Mafft (v4.745) (78) with the option -add for incorporating the outgroup species sequence and --keeplength. In the next step, each gene alignment was refined using the codon sequence aligner MACSE (v2.0.5) (79). The resulting protein-coding gene alignments were checked for the presence of start codons, absence of frameshifts, absence of terminal gaps, and a threshold of 5% gaps.

### Maximum-likelihood tree reconstruction

The protein sequences of each reference genome used to map the target species were used to establish the phylogenetic relationship among species. We included three outgroup species *Amanita muscaria* (PRJNA207684), *Pneumocystis jirovecii* (PRJNA223510), and *Rhizophagus irregularis* (PRJNA208392). Protein datasets were used for pairwise BLAST searches and ortholog reconstruction using OrthoFinder (v2.5.2) and default settings. A second run of OrthoFinder was performed to generate multiple sequence alignments (MSA) of single-copy orthologues using the Mafft aligner. A concatenated fasta file was retrieved of all single-copy orthogroups (*n*=1,065) with 100% of species having genes in any outgroup. Maximum-likelihood tree searches were performed in RAxML (v8.2.12) using a PROTGTR model, a randomized parsimony starting tree, and 100 Felsenstein bootstraps. The final tree was generated using ggtree in R programming language with *R. irregularis* set as root.

### Identification of DNA sequence deletions and accessory genes

We used information on genomic deletions from our focal species dataset to identify accessory genes. We first identified deletion events across the genome of each isolate using CNVnator (v0.4.1) on each bam file (80–82. CNVnator predicts deletion events based on read depth thresholds of mapping reads against a reference genome. We used bins of 100 bp to examine the genomes for deletions. We retained deletion events according to the following cut-offs: e_val1 (or t-test statistics) < 0.05, q0 (fraction of reads mapped with q0 quality) < 0.05, and normalized read depth < 0.4.

Next, we overlapped the prediction of deletion events with protein-coding gene coordinates. Using "read.gff" from the R package ape (v5.6-2) (83) we imported the coordinates of the protein-coding gene of each focal species using the gff file provided with the reference genome. We converted both deletion and gene coordinates into "Genomic Range" object using the R package GenomicRanges (v3.15) (84) and extracted genes overlapping with deleted regions with the mergeByOverlaps and "type = ’within’" function. Only genes completely overlapping with deleted regions were retained (i.e., partial gene deletions were not considered).

We constructed a dataset comprising: Core genes (genes present in all isolates in our dataset for a species), and accessory genes (genes missing from at least one isolate for a species). First, we selected accessory genes at a missingness frequency ranging between 0.1 and 5%. Next, we selected the isolates harboring the identified genes, from which we extracted sequence information for the accessory gene group. Finally, from the same isolates, we randomly chose an equal number of core genes to compose the core gene group. As proteins with an extracellular function are under strong selective pressure in some organisms (18), only genes encoding non-secreted proteins were considered.

### Prediction of gene function

Functional characterization was performed for the protein sequence of all reference genomes used to map the focal species with interproscan (v5.48-83.0) (Supplementary Table 4). We included the identification of signal peptides with SignalP (v4.1), the prediction of transmembrane helices in proteins with TMHMM (v2.0), and a look-up table (option -iprlookup) (85–87). We defined secreted proteins as genes predicted to have a signal peptide but lacking a transmembrane domain. Finally, using the genes classified as secreted proteins we used EffectorP (v3.0) to predict effector candidates (88,89). From the gene prediction, we divided genes across species into three datasets: non-secreted (protein-coding genes predicted as not secreted), secreted (protein-coding genes predicted as secreted, but not effectors), and effectors. Genes are unique to each category and only core genes were considered (Supplementary Table 5).

### Population recombination rate and definition of recombination (**p)** categories per species

To assess the relevance of recombination in adaptative evolution, we used the population genomic datasets to generate maps of the population recombination rate for all 20 focal species. As the individuals of all species included here are haploid, we calculated the population recombination rate (ρ) as ρ=2Ne*r, where Ne is the effective population size and r is the recombination rate per generation per nucleotide. Firstly, we estimated Watterson’s θ in 10 kb windows using ANGSD (90) and the bam file of each available isolate within each focal species. Watterson’s θ provides an estimation of the population mutation rate parameter. Second, we used LDhat (https://github.com/auton1/LDhat) to infer ρ. LDhat uses a pre-computed likelihood table for a given number of individuals and population mutation rate parameter. Due to computational limitations in calculating a likelihood table per species, we chose the one with a theta value closest to our estimation of Watterson’s θ for the given species. Also, for species with more than 100 individuals, we performed nine random subsampling of 100 individuals each and ran LDhat nine times to calculate a median value of ρ. If a species has a number of isolates lower than that available for the pre-computed likelihood tables, we used the option "lkgen" to reduce the pre-computed likelihood table and calculated the median value of ρ.

We further used the estimations of ρ across the genome of all species to gather genes in regions with similar recombination levels. We used the cut2 function in R, with levels.mean=TRUE. We varied the number of categories per species aiming to have an equivalent number of genes per ρ category (Supplementary Table 6).

### Estimation of transition/transversion rate, distribution of fitness effects, and rates of adaptation

The previously generated gene datasets were used to infer the synonymous and non-synonymous unfolded site frequency spectra (SFS), synonymous (Lps) and non-synonymous (Lpn) polymorphic sites, as well as synonymous (*d*_S_) and non-synonymous (*d*_N_) substitution rates. First, we used the program BppPopStats to estimate the transition over transversion rate (kappa, Supplementary Table 5) per gene in a maximum likelihood approach with a model of codon evolution (91). Next, a second run of BppPopStats was performed by fixing kappa to the median value obtained across all genes of each species, determining the expected number of polymorphisms and substitutions at synonymous and non-synonymous sites for each gene.

The distribution of fitness effects (DFE) was determined based on the SFS information of synonymous and non-synonymous sites of concatenated genes using the GRAPES software (3,14). GRAPES was run with the -no_div_param option, which estimates adaptive and non-adaptive rates from the fitted DFE. The divergence counts are only used to estimate the DFE, an approach that mitigates the influence of past demographic fluctuations (14,31,92,93). Six models were used to estimate the DFE: Neutral, GammaZero, GammaGamma, GammaExpo, Displaced Gamma, and Scaled Beta (Supplementary Table 7). The DFE models were compared using Akaike’s Information Criterion (94). A model averaging procedure was performed (95) when comparing the rates of adaptation among species, because several models had similar AIC values. Posterior to model fitting, the rates of evolution were estimated through ω_NA_ (rate of non-adaptive non-synonymous substitutions), ω_A_ (rate of adaptive non-synonymous substitutions), and α (proportion of amino-acid substitutions that are adaptive).

### Single-species statistical analyses

We used bootstraps to estimate the variation in each gene set’s adaptation rates. We performed 100 re-sampling in each gene set. The rates of adaptation were estimated for each bootstrap replicate using the model with the best fit for the species, and the estimates’ distribution was used to visualize confidence intervals.

A permutation test was implemented to assess the significance of differences in the adaptation rates between gene sets, shuffling all genes among predicted function (e.g., in pairwise combinations of non-secreted, secreted and effector gene sets) and between core and accessory 1,000 times. *P*-values were computed as by the formula: (Σ*i*[l*Si*| ≥ |*Sobs*|] + 1) / (*n* + 1), where Si represents the difference in parameter estimates for replicate I and [] denote the Iverson bracket, so that Σ*i*[|S*i*| ≥ |*Sobs*|] is the number of replicates for which the absolute difference is greater than or equal to the observed difference, and n is the number of replicates for which parameters could be successfully calculated (9).

### Generalized least square model and correlations

We performed statistical inferences using generalized least square models (GLS) implemented using the package nlme in R (R Core Team, 2022; Pinheiro et al., 2023).

We used both ω_A_ and ω_NA_ as response variables. As explanatory variables, we used *π_S_* and *ρ* averaged for each species. In another set of models, we further split genes per functional and recombination rate categories. Because substitution rates cannot be computed for each single genes but for a set of genes, we could only assess categorical factors independently of each other in separate models. We applied a log transformation for the *ρ* and *π_S_* variables to improve linearity. The variable “recombination category” was centered by subtracting the mean p for the species from each p category value. We assessed the occurrence of collinearity in the models and removed redundant factors using estimations of the variance inflation factor (VIF) (96,97). Factors were removed from the model if VIF > 3. To account for the non-independence of the residuals stemming from the phylogenetic signal (i.e., species displaying similar rates of adaptation simply due to more recent phylogenetical ancestry) we included a phylogenetic model of covariance (98). After fitting the models, we checked for the normality of residuals through visual inspection of residuals’ distribution and formal testing using the Shapiro-Wilk test (99,100). Residuals variance homogeneity (homoskedasticity) was also inspected, and the residual’s independence was assessed with a Ljung-Box test (101). Based on the previously described sanity checks, we performed a Box-Cox transformation on the response variable when requirements of normality and homoskedasticity were not fulfilled (102). The resulting following model structures were considered:

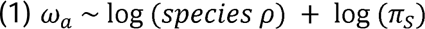

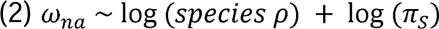

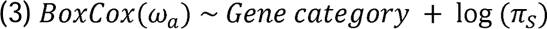

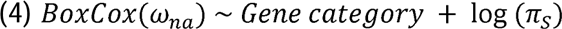

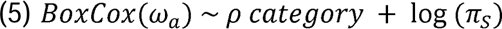

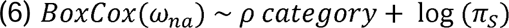

### Inference of demography and the effects of population structure on the rates of adaptation

To assess the robustness of ω_NA_, ω_A_, and α estimations against the effects of demography and population structure we performed simulations using different population dynamics and re-estimated the rates of adaptation (see supplementary file 1 for details). In brief, we used SLiM (v4.0.1) to simulate populations of 10,000 individuals evolving for 100,000 generations, with the first 10,000 generations discarded as burn-in (103). The simulated genome is constructed based on 1,500 genes of 500 bp each, separated by 500 bp of noncoding spacers. The recombination rate is uniform along the sequence and set to 1e-7 Mbp (10cM/Mb). The mutation rate, also uniform, is set to 2.2e-7 per bp. Only mutations within “genes” are recorded and divided into three semi-dominant mutations: neutral, deleterious, and advantageous. Neutral mutations were used to derive the neutral "synonymous" SFS, while deleterious and beneficial mutations were used to generate the "non-synonymous". We ran Grapes based on the SFS dataset of the following simulated scenarios: (i) panmictic population, (ii) panmictic population with exponential growth/shrinkage, (iii) population structure with two demes, (iv) population structure with two merging populations. Across all scenarios, estimations of the rates of adaptation were performed using the classical estimation (Basic, where α = 1 − pN/pS, ω_A_ = d_N_/d_S_ − p_N_/p_S_, ω_NA_ = p_N_/p_S_), Fay-Wyckoff-Wu (FWW) estimation (104), and a model based estimation that fits a GammaExpo DFE in Grapes (14). We compared the estimated ω_NA_, ω_A_, and α to the corresponding true values.

## Supporting information

Simulations

## Acknowledgments

The authors are thankful to Nicolas Galtier and Marjolaine Rousselle for helpful discussions and for sharing simulation scripts, as well as members of the Environmental Genomics group in Kiel for fruitful discussions. This research was funded by the DFG priority program SPP1819 (grant ID HO 4435/1-2) attributed to EHS and WS. This work was supported in part by the U.S. Department of Agriculture, Agricultural Research Service through USDA projects 3060-22000-051- 000D (Friesen) and 3060-21000-044-000-D (Bolton). Mention of trade names or commercial products in this publication is solely for the purpose of providing specific information and does not imply recommendation or endorsement by the U.S. Department of Agriculture._ USDA is an equal opportunity provider and employer.

## Data availability

All sequencing data is available from the NCBI Sequence Read Archive. Individual accession numbers are provided in the Supplementary Table 1.

## Code availability

All custom scripts used for analysis and data representation are available from https://github.com/DaniloASP/RatesOfAdaptation (DOI: 10.5281/zenodo.8289768). Guided steps, downstream processing and visualization of the data using R, Bash and Python are provided as R markdown tutorials.

