## Supplementary material for "Large-scale analyses reveal the contribution of adaptive evolution in pathogenic and non-pathogenic fungal species": RatesOfAdaptation_Pereira_Supplementary.pdf

**Table S1**. Detailed information on genome sources of focal and outgroup species and genome number per species. All species are haploid.

| Class | Focal or Outgroup | Species name | Bioproject | Number of genomes | Reference |
|---|---|---|---|---|---|
| Plant pathogen | Focal | *Aspergilus flavus* | PRJNA639008 | 94 | (Drott et al., 2020) |
| | Outgroup | Aspergillus caelatus (CBS 763.97 strain) | PRJNA333899 | 1 | |
| | Focal | *Botrytis cinerea* | PRJNA624742 | 32 | (Atwell et al., 2018; Mercier et al., 2021) |
| | | | PRJNA525902 | 97 | |
| | Outgroup | Botrytis fabae (33 strain) | PRJNA624742 | 1 | |
| | Focal | *Cercospora beticola* | PRJNA673877 | 190 | (Spanner et al., 2021) and this publication |
| | | | 10.5281/zenodo.8207821 | 22 | |
| | Outgroup | *Cercospora sojina (*DH strain) | PRJNA508859 | 1 | |
| | Focal | *Cryphonectria parasitica* | PRJNA604575 | 188 | (Stauber et al., 2020) |
| | | | PRJNA644891 | 43 | |
| | Outgroup | *Cryphonectria naterciae* (F2_M3664 strain) | PRJNA644891 | 1 | |
| | Focal | *Fusarium graminearum* | PRJNA397890 | 60 | (Kelly and Ward, 2018) |
| | Outgroup | *Fusarium pseudograminearum* (NRRL 28062 strain) | PRJNA397890 | 1 | |
| | Focal | *Magnaporthe oryzae* (rice host) | PRJNA354675 | 83 | (Islam et al., 2016; Zhong et al., 2018) |
| | | | PRJNA324432 | 25 | |
| | Outgroup | *Magnaporthe oryzae* (GFSI1-7-2, Setaria italica host) | PRJDB4326 | 1 | |
| | Focal | *Magnaporthe oryzae (triticum host)* | PRJNA324432 | 18 | (Islam et al., 2016; Zhong et al., 2018) |
| | Outgroup | *Magnaporthe oryzae* (GFSI1-7-2, Setaria italica host) | PRJDB4326 | 1 | |
| | Focal | *Ophiostoma novo-ulmi* subsp. americana | PRJNA566197 | 27 | (Comeau et al., 2015) |
| | Outgroup | *Ophiostoma piceae* (CB001 strain) | PRJNA182071 | 1 | |
| | Focal | *Ophiostoma novo-ulmi* subsp. novo-ulmi | PRJNA566197 | 28 | (Comeau et al., 2015) |
| | Outgroup | *Ophiostoma piceae* (CB001 strain) | PRJNA182071 | 1 | |

| | | | | | |
|---|---|---|---|---|---|
| | Focal | *Ophiostoma ulmi* | PRJNA566197 | 37 | (Comeau et al., 2015) |
| | Outgroup | *Ophiostoma piceae* (CB001 strain) | PRJNA182071 | 1 | |
| | Focal | *Parastagonospora nodorum* | PRJNA606320 | 177 | (Pereira et al., 2021; Richards et al., 2019) |
| | | | PRJNA398070 | 186 | |
| | Outgroup | *Parastagonospora jasniorum* (pnod27 strain) | PRJNA606320 | 1 | |
| | Focal | *Pyrenophora teres f. sp. teres* | 10.5281/zenodo.8183372 | 151 | This publication |
| | Outgroup | *Pyrenophora teres f. maculata* (G106_Pt124 strain) | | 1 | |
| | Focal | *Rhynchosporium commune* | PRJNA327656 | 120 | (Mohd-Assaad et al., 2016) |
| | Outgroup | *Rhynchobrunnera orthospora* (BA2_1_1 strain) | PRJNA419548 | 1 | |
| | Focal | *Sclerotinia sclerotiorum* | PRJNA516948 | 24 | (Derbyshire et al., 2019) |
| | Outgroup | Sclerotinia minor (SsChi strain) | PRJNA516948 | 1 | |
| | Focal | *Sphaerulina musiva* | PRJNA543887 | 122 | (Tabima et al., 2020)t |
| | Outgroup | *Mycosphaerella populicola* (p0202b strain) | PRJNA277053 | 1 | |
| | Focal | *Venturia inaequalis* | PRJNA407103 | 83 | (Le Cam et al., 2019) |
| | Outgroup | *Venturia aucupariae* (2371 strain) | PRJNA407103 | 1 | |
| | Focal | *Verticillium dahliae* | PRJNA171348 | 78 | (de Jonge et al., 2013; Zhang et al., 2019) |
| | | | PRJNA169154 | 11 | |
| | | | PRJNA693498 | 10 | |
| | | | PRJNA639910 | 7 | |
| | Outgroup | Verticillium nonalfalfae (TAB2 strain) | PRJNA283258 | 1 | |
| | Focal | *Zymoseptoria tritici* | PRJNA327615 | 129 | (Hartmann et al., 2017; Oggenfuss et al., 2021) |
| | | | PRJNA596434 | 290 | |
| | | | PRJNA178194 | 9 | |
| | | | PRJNA480739 | 57 | |
| | Outgroup | *Zymoseptoria ardabiliae* (STIR041_1_1_1 strain) | Wallace (cluster) | 1 | |
| Non-pathogenic | Focal | *Penicillium biforme* | PRJNA655754 | 28 | (Ropars et al., 2020) |
| | Outgroup | *Penicillium arizonense* (CBS:141311 strain) | PRJNA397890 | 1 | |
| | Focal | Neurospora discreta | PRJNA295117 | 52 | |

| | | | | |
|---|---|---|---|---|
| Outgroup | Neurospora metzenbergii (P4149 strain) | PRJNA486257 | 1 | (Gladieux et al., 2020) |

**Table S2.** Reference genome used for mapping reads of 20 focal species.

| Focal species | Reference genome | NCBI / JGI acession | Median GC | Reference |
|---|---|---|---|---|
| *Aspergilus flavus* | NRRL3357 | GCA_014117465.1 | 48 | (Nierman et al., 2015) |
| *Botrytis cinerea* | B05.10 | GCA_000143535.4 | 42 | (Van Kan et al., 2017) |
| *Cercospora beticola* | 09-40 | GCF_002742065.1 | 51.3 | (de Jonge et al., 2018) |
| *Cryphonectria parasitica* | EP155 | GCF_011745365.1 | 50.8 | (Demené et al., 2019) |
| *Fusarium graminearum* | PH-1 | GCA_000240135.3 | 48.3 | (Lu et al., 2022) |
| *Magnaporthe oryzae* (rice host) | 70-15 | GCA_000002495.2 | 51 | (Dean et al., 2005) |
| *Magnaporthe oryzae* (triticum host) | 70-15 | GCA_000002495.2 | 51 | |
| *Neurospora discreta* | FGSC 8579 | http://fungidb.org | | |
| *Ophiostoma novo-ulmi* subsp. *americana* | H327 | https://mycocosm.jgi.doe.gov/Ophnu1/Ophnu1.home.html | | (Forgetta et al., 2013) |
| *Ophiostoma novo-ulmi* subsp. *novo-ulmi* | H327 | https://mycocosm.jgi.doe.gov/Ophnu1/Ophnu1.home.html | | |
| *Ophiostoma ulmi* | H327 | https://mycocosm.jgi.doe.gov/Ophnu1/Ophnu1.home.html | | |
| *Parastagonospora nodorum* | Sn15 | GCA_016801405.1 | 50.3 | (Bertazzoni et al., 2021) |
| *Penicillium biforme* | ASM207223v1 | GCA_002072235.1 | 47.9 | (Nielsen et al., 2017) |
| *Pyrenophora teres* f. sp. *teres* | 0-1 | GCA_000166005.1 | 50.9 | (Ellwood et al., 2010) |
| *Rhynchosporium commune* | UK7 | GCA_900074885.1 | 42.3 | (Torriani et al., 2014) |
| *Sclerotinia sclerotiorum* | 1980 UF-70 | GCA_000146945.2 | 41.8 | (Amselem et al., 2011) |
| *Sphaerulina musiva* | SO2202 | GCA_000320565.2 | 51.1 | (Ohm et al., 2012) |
| *Venturia inaequalis* | Vi1 | https://mycocosm.jgi.doe.gov/Venin1/Venin1.info.html | | (Deng et al., 2017) |
| *Verticillium dahliae* | VdLs.17 | GCA_000150675.2 | 55.6 | (Klosterman et al., 2011) |
| *Zymoseptoria tritici* | IPO323 | GCF_000219625.1 | 52.2 | (Goodwin et al., 2011) |

**Table S3.** Number of bi-allelic SNPs in the gene dataset without missing information for the target species dataset. Species were mapped against the stablished reference genome.

| Species | Number of SNPs (without missing information) |
|---|---|
| *Aspergilus flavus* | 626,827 |
| *Botrytis cinerea* | 140,615 |
| *Cercospora beticola* | 406,625 |
| *Cryphonectria parasitica* | 64,568 |
| *Fusarium graminearum* | 377,483 |
| *Magnaporthe oryzae* rice | 40,269 |
| *Magnaporthe oryzae Triticum* | 307,642 |
| *Neurospora discreta* | 1,490,695 |
| *Ophiostoma novo-ulmi subsp. americana* | 261,957 |
| *Ophiostoma novo-ulmi subsp. novo-ulmi* | 393,379 |
| *Ophiostoma ulmi* | 1,131,849 |
| *Parastagonospora nodorum* | 980,923 |
| *Penicillium biforme* | 1,093,012 |
| *Pyrenophora teres f. sp. teres* | 30,321 |
| *Rhynchosporium commune* | 495,281 |
| *Sclerotinia sclerotiorum* | 1,664,053 |
| *Sphaerulina musiva* | 2,566,304 |
| *Venturia inaequalis* | 687,494 |
| *Verticillium dahliae* | 75,649 |
| *Zymoseptoria tritici* | 547,711 |

**Table S4.** The number of proteins per target species**.** Secreted and effectors are based on the pipeline described in the methods section. Effectors are a subset of secreted proteins.

| Species | Total number of proteins for the species as: | | |
|---|---|---|---|
| | All | Secreted | Effectors |
| *Aspergilus flavus* | 12009 | 984 | 317 |
| *Botrytis cinerea* | 13703 | 958 | 350 |
| *Cercospora beticola* | 12463 | 1081 | 376 |
| *Cryphonectria parasitica* | 11588 | 734 | 253 |
| *Fusarium graminearum* | 13313 | 1022 | 389 |
| *Magnaporthe oryzae rice* | 12989 | 1444 | 615 |
| *Magnaporthe oryzae Triticum* | 12989 | 1444 | 615 |
| *Neurospora discreta* | 9948 | 583 | 207 |
| *Ophiostoma novo-ulmi subsp. americana* | 8640 | 446 | 113 |
| *Ophiostoma novo-ulmi subsp. novo-ulmi* | 8640 | 446 | 113 |
| *Ophiostoma ulmi* | 8640 | 446 | 113 |
| *Parastagonospora nodorum* | 13605 | 1162 | 390 |
| *Penicillium biforme* | 11870 | 733 | 225 |
| *Pyrenophora teres f. sp. teres* | 11578 | 825 | 285 |
| *Rhynchosporium commune* | 12211 | 992 | 337 |
| *Sclerotinia sclerotiorum* | 14490 | 695 | 278 |
| *Sphaerulina musiva* | 10144 | 647 | 226 |
| *Venturia inaequalis* | 13233 | 1361 | 627 |
| *Verticillium dahliae* | 10535 | 908 | 234 |
| *Zymoseptoria tritici* | 11839 | 935 | 365 |

**Table S5.** Estimations of transition / transversion rate ratio (kappa) based on core genes, total number of core genes, number of genes classified as non-secreted, secreted and effectors.

| Species | kappa | Total number of core genes | Number of protein-coding genes grouped as: | | |
|---|---|---|---|---|---|
| | | | Non-secreted | Secreted | Effectors |
| *Aspergilus flavus* | 3.92 | 5909 | 5480 | 277 | 152 |
| *Botrytis cinerea* | 5.27 | 3307 | 3102 | 136 | 69 |
| *Cercospora beticola* | 3.21 | 6523 | 6107 | 280 | 136 |
| *Cryphonectria parasitica* | 3.55 | 5431 | 5111 | 194 | 126 |
| *Fusarium graminearum* | 5.35 | 7727 | 7173 | 342 | 212 |
| *Magnaporthe oryzae rice* | 2.64 | 7122 | 6490 | 386 | 246 |
| *Magnaporthe oryzae Triticum* | 3.68 | 7415 | 6700 | 434 | 281 |
| *Neurospora discreta* | 3.86 | 4694 | 4452 | 151 | 91 |
| *Ophiostoma novo-ulmi subsp. americana* | 2.19 | 4682 | 4477 | 139 | 66 |
| *Ophiostoma novo-ulmi subsp. novo-ulmi* | 2.23 | 4693 | 4485 | 141 | 67 |
| *Ophiostoma ulmi* | 2.40 | 4813 | 4593 | 150 | 70 |
| *Parastagonospora nodorum* | 1.63 | 4630 | 4352 | 176 | 102 |
| *Penicillium biforme* | 2.14 | 4351 | 4162 | 124 | 65 |
| *Pyrenophora teres f. sp. teres* | 5.07 | 6899 | 6418 | 317 | 164 |
| *Rhynchosporium commune* | 2.55 | 5767 | 5471 | 200 | 96 |
| *Sclerotinia sclerotiorum* | 3.73 | 5756 | 5420 | 207 | 129 |
| *Sphaerulina musiva* | 3.47 | 5489 | 5228 | 163 | 98 |
| *Venturia inaequalis* | 3.14 | 5495 | 5174 | 206 | 115 |
| *Verticillium dahliae* | 3.92 | 3800 | 3500 | 226 | 74 |
| *Zymoseptoria tritici* | 3.05 | 4935 | 4697 | 163 | 75 |

**Table S6.** Estimations of rho, number of genes within each rho category and total rho categories per species.

| Species | Species_rho | Rho_category | N_of_genes_per_category | Total_of_categories_[illegible] |
|---|---|---|---|---|
| *A. flavus* | 0.002170 | 1.00E-05 | 620 | 10 |
| *A. flavus* | 0.002170 | 5.31E-05 | 477 | 10 |
| *A. flavus* | 0.002170 | 0.00013058 | 548 | 10 |
| *A. flavus* | 0.002170 | 0.00021745 | 547 | 10 |
| *A. flavus* | 0.002170 | 0.00035502 | 550 | 10 |
| *A. flavus* | 0.002170 | 0.00055791 | 552 | 10 |
| *A. flavus* | 0.002170 | 0.00098383 | 543 | 10 |
| *A. flavus* | 0.002170 | 0.0016014 | 549 | 10 |
| *A. flavus* | 0.002170 | 0.0025235 | 548 | 10 |
| *A. flavus* | 0.002170 | 0.0051079 | 546 | 10 |
| *B. cinera* | 0.002231 | 0 | 2242 | 2 |
| *B. cinera* | 0.002231 | 0.002754 | 860 | 2 |
| *C. beticola* | 0.001603 | 4.32E-05 | 411 | 15 |
| *C. beticola* | 0.001603 | 8.59E-05 | 408 | 15 |
| *C. beticola* | 0.001603 | 0.00012914 | 403 | 15 |
| *C. beticola* | 0.001603 | 0.00017348 | 407 | 15 |
| *C. beticola* | 0.001603 | 0.00021756 | 408 | 15 |
| *C. beticola* | 0.001603 | 0.00029417 | 407 | 15 |
| *C. beticola* | 0.001603 | 0.00039573 | 409 | 15 |
| *C. beticola* | 0.001603 | 0.00052719 | 407 | 15 |
| *C. beticola* | 0.001603 | 0.00067654 | 409 | 15 |
| *C. beticola* | 0.001603 | 0.00086783 | 403 | 15 |
| *C. beticola* | 0.001603 | 0.0011358 | 409 | 15 |
| *C. beticola* | 0.001603 | 0.0015739 | 406 | 15 |
| *C. beticola* | 0.001603 | 0.0023324 | 407 | 15 |
| *C. beticola* | 0.001603 | 0.0037664 | 411 | 15 |
| *C. beticola* | 0.001603 | 0.0090464 | 402 | 15 |
| *C. parasitica* | 0.004620 | 7.17E-05 | 344 | 15 |
| *C. parasitica* | 0.004620 | 0.00010905 | 338 | 15 |
| *C. parasitica* | 0.004620 | 0.00013214 | 341 | 15 |
| *C. parasitica* | 0.004620 | 0.00015493 | 340 | 15 |
| *C. parasitica* | 0.004620 | 0.00017882 | 350 | 15 |
| *C. parasitica* | 0.004620 | 0.00020189 | 333 | 15 |
| *C. parasitica* | 0.004620 | 0.00023686 | 340 | 15 |
| *C. parasitica* | 0.004620 | 0.00027689 | 340 | 15 |
| *C. parasitica* | 0.004620 | 0.00032094 | 341 | 15 |
| *C. parasitica* | 0.004620 | 0.0003722 | 341 | 15 |
| *C. parasitica* | 0.004620 | 0.0004341 | 343 | 15 |
| *C. parasitica* | 0.004620 | 0.00053707 | 342 | 15 |
| *C. parasitica* | 0.004620 | 0.00071082 | 338 | 15 |
| *C. parasitica* | 0.004620 | 0.0011031 | 344 | 15 |
| *C. parasitica* | 0.004620 | 0.0088023 | 336 | 15 |
| *F. graminearum* | 0.001331 | 9.99E-06 | 1856 | 4 |
| *F. graminearum* | 0.001331 | 8.02E-05 | 1732 | 4 |
| *F. graminearum* | 0.001331 | 0.00030674 | 1792 | 4 |
| *F. graminearum* | 0.001331 | 0.0035751 | 1793 | 4 |
| *M. oryzae rice* | 0.000922 | 9.85E-06 | 1429 | 5 |
| *M. oryzae rice* | 0.000922 | 1.44E-05 | 1169 | 5 |
| *M. oryzae rice* | 0.000922 | 2.05E-05 | 1376 | 5 |
| *M. oryzae rice* | 0.000922 | 3.14E-05 | 1314 | 5 |
| *M. oryzae rice* | 0.000922 | 0.00083796 | 1202 | 5 |
| *M. oryzae Triticum* | 0.008401 | 0 | 1292 | 3 |
| *M. oryzae Triticum* | 0.008401 | 9.88E-06 | 4667 | 3 |
| *M. oryzae Triticum* | 0.008401 | 0.026787 | 741 | 3 |
| *N. discreta* | 0.001410 | 9.87E-06 | 1039 | 5 |

| | | | | |
|---|---|---|---|---|
| *N. discreta* | 0.001410 | 1.00E-05 | 774 | 5 |
| *N. discreta* | 0.001410 | 8.35E-05 | 861 | 5 |
| *N. discreta* | 0.001410 | 0.00038812 | 888 | 5 |
| *N. discreta* | 0.001410 | 0.0013265 | 890 | 5 |
| *O. novo-ulmi subsp. americana* | 0.000380 | 9.97E-06 | 3582 | 2 |
| *O. novo-ulmi subsp. americana* | 0.000380 | 0.0004623 | 895 | 2 |
| *O. novo-ulmi subsp. novo-ulmi* | 0.001192 | 9.98E-06 | 523 | 10 |
| *O. novo-ulmi subsp. novo-ulmi* | 0.001192 | 3.71E-05 | 386 | 10 |
| *O. novo-ulmi subsp. novo-ulmi* | 0.001192 | 8.33E-05 | 437 | 10 |
| *O. novo-ulmi subsp. novo-ulmi* | 0.001192 | 0.00017678 | 448 | 10 |
| *O. novo-ulmi subsp. novo-ulmi* | 0.001192 | 0.00034418 | 449 | 10 |
| *O. novo-ulmi subsp. novo-ulmi* | 0.001192 | 0.00056297 | 450 | 10 |
| *O. novo-ulmi subsp. novo-ulmi* | 0.001192 | 0.00089122 | 447 | 10 |
| *O. novo-ulmi subsp. novo-ulmi* | 0.001192 | 0.0013605 | 448 | 10 |
| *O. novo-ulmi subsp. novo-ulmi* | 0.001192 | 0.002222 | 451 | 10 |
| *O. novo-ulmi subsp. novo-ulmi* | 0.001192 | 0.00586 | 446 | 10 |
| *O. ulmi* | 0.000057 | 1.00E-05 | 2615 | 2 |
| *O. ulmi* | 0.000057 | 5.37E-05 | 1978 | 2 |
| *P. biforme* | 0.011405 | 1.00E-05 | 2081 | 2 |
| *P. biforme* | 0.011405 | 0.022637 | 2081 | 2 |
| *P. nodorum* | 0.017821 | 0.00050455 | 292 | 15 |
| *P. nodorum* | 0.017821 | 0.00129223 | 289 | 15 |
| *P. nodorum* | 0.017821 | 0.00231873 | 290 | 15 |
| *P. nodorum* | 0.017821 | 0.00341133 | 291 | 15 |
| *P. nodorum* | 0.017821 | 0.00477346 | 290 | 15 |
| *P. nodorum* | 0.017821 | 0.0060776 | 294 | 15 |
| *P. nodorum* | 0.017821 | 0.00724309 | 288 | 15 |
| *P. nodorum* | 0.017821 | 0.00870338 | 290 | 15 |
| *P. nodorum* | 0.017821 | 0.01048706 | 291 | 15 |
| *P. nodorum* | 0.017821 | 0.01279628 | 287 | 15 |
| *P. nodorum* | 0.017821 | 0.01633377 | 292 | 15 |
| *P. nodorum* | 0.017821 | 0.02180633 | 289 | 15 |
| *P. nodorum* | 0.017821 | 0.02980106 | 293 | 15 |
| *P. nodorum* | 0.017821 | 0.04421197 | 288 | 15 |
| *P. nodorum* | 0.017821 | 0.11097865 | 288 | 15 |
| *P. teres f. teres* | 0.000322 | 1.69E-05 | 429 | 15 |
| *P. teres f. teres* | 0.000322 | 4.13E-05 | 431 | 15 |
| *P. teres f. teres* | 0.000322 | 6.72E-05 | 471 | 15 |
| *P. teres f. teres* | 0.000322 | 8.44E-05 | 393 | 15 |
| *P. teres f. teres* | 0.000322 | 9.65E-05 | 419 | 15 |
| *P. teres f. teres* | 0.000322 | 0.00011185 | 425 | 15 |
| *P. teres f. teres* | 0.000322 | 0.00012816 | 432 | 15 |
| *P. teres f. teres* | 0.000322 | 0.00014657 | 423 | 15 |
| *P. teres f. teres* | 0.000322 | 0.00017037 | 428 | 15 |
| *P. teres f. teres* | 0.000322 | 0.00019926 | 432 | 15 |
| *P. teres f. teres* | 0.000322 | 0.00024126 | 428 | 15 |
| *P. teres f. teres* | 0.000322 | 0.00028074 | 426 | 15 |
| *P. teres f. teres* | 0.000322 | 0.00033082 | 431 | 15 |

| | | | | |
|---|---|---|---|---|
| *P. teres f. teres* | 0.000322 | 0.00042621 | 424 | 15 |
| *P. teres f. teres* | 0.000322 | 0.0012511 | 426 | 15 |
| *R. commune* | 0.006148 | 0.00030559 | 365 | 15 |
| *R. commune* | 0.006148 | 0.00071794 | 368 | 15 |
| *R. commune* | 0.006148 | 0.00101103 | 363 | 15 |
| *R. commune* | 0.006148 | 0.00135262 | 364 | 15 |
| *R. commune* | 0.006148 | 0.00166486 | 364 | 15 |
| *R. commune* | 0.006148 | 0.00198078 | 368 | 15 |
| *R. commune* | 0.006148 | 0.00241026 | 365 | 15 |
| *R. commune* | 0.006148 | 0.00284836 | 361 | 15 |
| *R. commune* | 0.006148 | 0.00333494 | 365 | 15 |
| *R. commune* | 0.006148 | 0.00392059 | 368 | 15 |
| *R. commune* | 0.006148 | 0.0046525 | 362 | 15 |
| *R. commune* | 0.006148 | 0.0055177 | 364 | 15 |
| *R. commune* | 0.006148 | 0.00677076 | 366 | 15 |
| *R. commune* | 0.006148 | 0.00887086 | 364 | 15 |
| *R. commune* | 0.006148 | 0.02468806 | 364 | 15 |
| *S. musiva* | 0.117371 | 0.0026484 | 350 | 15 |
| *S. musiva* | 0.117371 | 0.0116438 | 351 | 15 |
| *S. musiva* | 0.117371 | 0.0241135 | 347 | 15 |
| *S. musiva* | 0.117371 | 0.0356612 | 347 | 15 |
| *S. musiva* | 0.117371 | 0.0455559 | 348 | 15 |
| *S. musiva* | 0.117371 | 0.0535576 | 351 | 15 |
| *S. musiva* | 0.117371 | 0.0630465 | 350 | 15 |
| *S. musiva* | 0.117371 | 0.0731292 | 346 | 15 |
| *S. musiva* | 0.117371 | 0.0864802 | 347 | 15 |
| *S. musiva* | 0.117371 | 0.1036392 | 351 | 15 |
| *S. musiva* | 0.117371 | 0.1214073 | 347 | 15 |
| *S. musiva* | 0.117371 | 0.1449064 | 350 | 15 |
| *S. musiva* | 0.117371 | 0.1840964 | 346 | 15 |
| *S. musiva* | 0.117371 | 0.2485171 | 350 | 15 |
| *S. musiva* | 0.117371 | 0.4386902 | 347 | 15 |
| *S. sclerotiorum* | 0.000098 | 6.03E-07 | 2721 | 2 |
| *S. sclerotiorum* | 0.000098 | 2.26E-05 | 2699 | 2 |
| *V. dahliae* | 0.000995 | 9.81E-06 | 1750 | 2 |
| *V. dahliae* | 0.000995 | 0.0015852 | 1750 | 2 |
| *V. inaequalis* | 0.004688 | 8.57E-06 | 1777 | 3 |
| *V. inaequalis* | 0.004688 | 0.00024445 | 1675 | 3 |
| *V. inaequalis* | 0.004688 | 0.0025087 | 1722 | 3 |
| *Z. tritici* | 0.035691 | 0.0034166 | 316 | 15 |
| *Z. tritici* | 0.035691 | 0.0082738 | 312 | 15 |
| *Z. tritici* | 0.035691 | 0.0123093 | 313 | 15 |
| *Z. tritici* | 0.035691 | 0.0159844 | 312 | 15 |
| *Z. tritici* | 0.035691 | 0.019551 | 316 | 15 |
| *Z. tritici* | 0.035691 | 0.0224869 | 310 | 15 |
| *Z. tritici* | 0.035691 | 0.025554 | 313 | 15 |
| *Z. tritici* | 0.035691 | 0.0294889 | 314 | 15 |
| *Z. tritici* | 0.035691 | 0.0339806 | 316 | 15 |
| *Z. tritici* | 0.035691 | 0.0385984 | 313 | 15 |
| *Z. tritici* | 0.035691 | 0.0439783 | 312 | 15 |
| *Z. tritici* | 0.035691 | 0.0509697 | 314 | 15 |
| *Z. tritici* | 0.035691 | 0.0604389 | 310 | 15 |
| *Z. tritici* | 0.035691 | 0.0733526 | 313 | 15 |
| *Z. tritici* | 0.035691 | 0.1146989 | 313 | 15 |

**Table S7.** Description of models for the distribution of fitness effects and premises used in GRAPES (Galtier, 2016).

| DFE model | Premise |
|---|---|
| Neutral | This model considers two categories for the DFE: (i) neutral mutations and (ii) strongly deleterious mutations, with neglectable contribution to polymorphism or divergence |
| GammaZero | Models the distribution of DFE considering mutations with slight negative effects on fitness, which can contribute to polymorphism and divergence. These mutations are modeled as a Gamma distribution. No inclusion of positive effects. |
| GammaGamma | Builds on the previous model but assumes the existence of mutations with weak positive effects on fitness. These new mutations are modeled as a Gamma distribution. |
| GammaExpo | Builds upon the previous model, but here weakly advantageous mutations are assumed to be exponentially distributed. |
| DisplGamma | This model assumes a displaced negative Gamma distribution of weakly advantageous mutations. |
| ScaledBeta | Builds on the neutral model but includes a class of strongly deleterious mutations, but assumes a Beta-shaped distribution of weak-effect mutations |

**Table S8.** Summary statistics for alignments between target and outgroup species. Pn, Ps, Dn, Ds refer to the total number of nonsynonymous and synonymous SNPs and substitutions, respectively. Lpn, Lps, Ldn, Lds refer to the total number of nonsynonymous and synonymous sites available in the within-species and between-species gene alignments, respectively.

| Target species | Pn | Lpn | Ps | Lps | Dn | Ldn | Ds | Lds | pNpS | dS | dN | dNdS |
|---|---|---|---|---|---|---|---|---|---|---|---|---|
| A. flavus | 22609 | 6490458 | 41839 | 2585184 | 190403.7 | 6490849 | 476795.3 | 2584793 | 0.2152 | 0.1845 | 0.0293 | 0.1590 |
| B. cinera | 7301 | 3574744 | 13497 | 1458632 | 20040.33 | 3574640 | 33241.67 | 1458736 | 0.2207 | 0.0228 | 0.0056 | 0.2460 |
| C. beticola | 21787 | 7224824 | 76074 | 2762902 | 203696.7 | 7223646 | 627819.3 | 2764080 | 0.1095 | 0.2271 | 0.0282 | 0.1241 |
| C. parasitica | 2998 | 5346821 | 3955 | 2097697 | 193314.3 | 5345542 | 422555.7 | 2098976 | 0.2974 | 0.2013 | 0.0362 | 0.1796 |
| F. graminearum | 20788 | 7788257 | 43344 | 3202354 | 95281 | 7788171 | 321143 | 3202440 | 0.1972 | 0.1003 | 0.0122 | 0.1220 |
| M. oryzae rice | 5521 | 7826550 | 3616 | 2912193 | 7072.083 | 7826533 | 8198.917 | 2912210 | 0.5681 | 0.0028 | 0.0009 | 0.3210 |
| M. oryzae Triticum | 9774 | 8017209 | 18503 | 3160620 | 15869.58 | 8017292 | 26060.42 | 3160537 | 0.2082 | 0.0082 | 0.0020 | 0.2401 |
| N. discreta | 42024 | 5460666 | 105849 | 2153469 | 162048.9 | 5460831 | 344786.1 | 2153304 | 0.1566 | 0.1601 | 0.0297 | 0.1853 |
| O. americana | 3200 | 5505003 | 15945 | 2016879 | 354208.6 | 5502939 | 870032.4 | 2018943 | 0.0735 | 0.4309 | 0.0644 | 0.1494 |
| O. novo | 6901 | 5507002 | 36100 | 2023979 | 354021 | 5505270 | 869979 | 2025711 | 0.0703 | 0.4295 | 0.0643 | 0.1497 |
| O. ulmi | 22332 | 5650391 | 111785 | 2107516 | 364514.5 | 5649601 | 896041.5 | 2108306 | 0.0745 | 0.4250 | 0.0645 | 0.1518 |
| P. biforme | 1786 | 4779826 | 7096 | 1710014 | 404431.1 | 4779918 | 874553.9 | 1709922 | 0.0900 | 0.5115 | 0.0846 | 0.1654 |
| P. nodorum | 1409 | 4910551 | 5293 | 1639940 | 466766.3 | 4909764 | 1027515 | 1640727 | 0.0889 | 0.6263 | 0.0951 | 0.1518 |
| P. teres f. teres | 5114 | 7128290 | 5664 | 2918524 | 25471 | 7128248 | 42420 | 2918566 | 0.3697 | 0.0145 | 0.0036 | 0.2458 |
| R. commune | 20648 | 6592775 | 22800 | 2392636 | 442506.5 | 6592727 | 1018476 | 2392684 | 0.3287 | 0.4257 | 0.0671 | 0.1577 |
| S. musiva | 115218 | 5932220 | 449600 | 2304427 | 115562.8 | 5931553 | 238861.3 | 2305094 | 0.0995 | 0.1036 | 0.0195 | 0.1880 |
| S. sclerotiorum | 85683 | 6146514 | 238005 | 2365827 | 180131.3 | 6145713 | 390407.7 | 2366628 | 0.1386 | 0.1650 | 0.0293 | 0.1777 |
| V. dahliae | 2498 | 4148984 | 3949 | 1646737 | 72236.08 | 4149201 | 188069.9 | 1646520 | 0.2511 | 0.1142 | 0.0174 | 0.1524 |
| V. inaequalis | 42086 | 6253391 | 296469 | 2364406 | 353016.6 | 6253539 | 769244.4 | 2364258 | 0.0537 | 0.3254 | 0.0565 | 0.1735 |
| Z. tritici | 42725 | 5671083 | 99867 | 2148702 | 148123.4 | 5670784 | 460715.6 | 2149001 | 0.1621 | 0.2144 | 0.0261 | 0.1218 |

**Table S9.** Estimates of rates of adaptation under various models of distribution of fitness effects for the core portion of protein-coding genes across species. Akaike's information criterion (AIC).

| Data | model | Log likelihood | AIC | alpha | omegaA | omegaNA | Species |
|---|---|---|---|---|---|---|---|
| All core genes | Neutral | -41153.118 | 82498.236 | 0.9 | 0.143 | 0.016 | *A. flavus* |
| | GammaZero | -834.056 | 1862.112 | 0 | 0 | 0.159 | *A. flavus* |
| | GammaGamma | -861.656 | 1923.312 | 0 | 0 | 0.159 | *A. flavus* |
| | GammaExpo | -814.796 | 1827.593 | 0.148 | 0.024 | 0.136 | *A. flavus* |
| | DisplGamma | -879.566 | 1955.131 | 0 | 0 | 0.159 | *A. flavus* |
| | ScaledBeta | -860.48 | 1916.959 | 0.161 | 0.025 | 0.134 | *A. flavus* |
| | Average | NA | NA | 0.147 | 0.023 | 0.136 | *A. flavus* |
| All core genes | Neutral | -38920.052 | 78100.103 | 0.988 | 0.243 | 0.003 | *B. cinera* |
| | GammaZero | -1154.033 | 2570.066 | 0 | 0 | 0.246 | *B. cinera* |
| | GammaGamma | -764.196 | 1796.393 | 0.455 | 0.112 | 0.134 | *B. cinera* |
| | GammaExpo | -764.182 | 1794.363 | 0.462 | 0.114 | 0.132 | *B. cinera* |
| | DisplGamma | -1123.861 | 2511.722 | 0 | 0 | 0.246 | *B. cinera* |
| | ScaledBeta | -796.053 | 1856.106 | 0.359 | 0.088 | 0.158 | *B. cinera* |
| | Average | NA | NA | 0.46 | 0.113 | 0.133 | *B. cinera* |
| All core genes | Neutral | -84349.203 | 169126.405 | 0.935 | 0.116 | 0.008 | *C. beticola* |
| | GammaZero | -2960.461 | 6350.921 | 0 | 0 | 0.124 | *C. beticola* |
| | GammaGamma | -5771.613 | 11979.227 | 0 | 0 | 0.124 | *C. beticola* |
| | GammaExpo | -1826.09 | 4086.18 | 0.486 | 0.06 | 0.064 | *C. beticola* |
| | DisplGamma | -3681.131 | 7794.262 | 0 | 0 | 0.124 | *C. beticola* |
| | ScaledBeta | -2316.039 | 5064.079 | 0.249 | 0.031 | 0.093 | *C. beticola* |
| | Average | NA | NA | 0.486 | 0.06 | 0.064 | *C. beticola* |
| All core genes | Neutral | -3742.992 | 7951.983 | 0.847 | 0.152 | 0.027 | *C. parasitica* |
| | GammaZero | -845.677 | 2159.354 | 0 | 0 | 0.18 | *C. parasitica* |
| | GammaGamma | -865.162 | 2204.324 | 0 | 0 | 0.18 | *C. parasitica* |
| | GammaExpo | -855.722 | 2183.445 | 0.072 | 0.013 | 0.167 | *C. parasitica* |
| | DisplGamma | -847.78 | 2165.561 | 0 | 0 | 0.18 | *C. parasitica* |
| | ScaledBeta | -878.922 | 2227.844 | 0.269 | 0.048 | 0.131 | *C. parasitica* |
| | Average | NA | NA | 0 | 0 | 0.18 | *C. parasitica* |
| All core genes | Neutral | -66467.158 | 133058.316 | 0.952 | 0.116 | 0.006 | *F. graminearum* |
| | GammaZero | -522.443 | 1170.886 | 0 | 0 | 0.122 | *F. graminearum* |
| | GammaGamma | -522.444 | 1176.888 | 0 | 0 | 0.122 | *F. graminearum* |
| | GammaExpo | -524.217 | 1178.435 | 0.012 | 0.001 | 0.121 | *F. graminearum* |
| | DisplGamma | -520.959 | 1169.918 | 0 | 0 | 0.122 | *F. graminearum* |
| | ScaledBeta | -554.063 | 1236.127 | 0.025 | 0.003 | 0.119 | *F. graminearum* |
| | Average | NA | NA | 0 | 0 | 0.122 | *F. graminearum* |
| All core genes | Neutral | -15560.293 | 31340.586 | 0.996 | 0.32 | 0.001 | *M. oryzae rice* |
| | GammaZero | -440.356 | 1102.711 | 0 | 0 | 0.321 | *M. oryzae rice* |
| | GammaGamma | -440.356 | 1108.711 | 0 | 0 | 0.321 | *M. oryzae rice* |
| | GammaExpo | -439.148 | 1104.296 | 0 | 0 | 0.321 | *M. oryzae rice* |
| | DisplGamma | -431.796 | 1087.591 | 0 | 0 | 0.321 | *M. oryzae rice* |
| | ScaledBeta | -473.152 | 1170.304 | 0.208 | 0.067 | 0.254 | *M. oryzae rice* |
| | Average | NA | NA | 0 | 0 | 0.321 | *M. oryzae rice* |
| All core genes | Neutral | -70252.55 | 140545.101 | 0.996 | 0.239 | 0.001 | *M. oryzae Triticum* |
| | GammaZero | -415.534 | 873.068 | 0 | 0 | 0.24 | *M. oryzae Triticum* |
| | GammaGamma | -181.739 | 411.477 | 0.266 | 0.064 | 0.176 | *M. oryzae Triticum* |
| | GammaExpo | -175.836 | 397.672 | 0.294 | 0.07 | 0.17 | *M. oryzae Triticum* |
| | DisplGamma | -424.872 | 893.744 | 0 | 0 | 0.24 | *M. oryzae Triticum* |
| | ScaledBeta | -204.077 | 452.153 | 0.299 | 0.072 | 0.168 | *M. oryzae Triticum* |

| | | | | | | | |
|---|---|---|---|---|---|---|---|
| | Average | NA | NA | 0.294 | 0.07 | 0.17 | *M. oryzae Triticum* |
| All core genes | Neutral | -127935.044 | 255978.087 | 0.936 | 0.174 | 0.012 | *N. discreta* |
| | GammaZero | -2917.134 | 5944.268 | 0 | 0 | 0.185 | *N. discreta* |
| | GammaGamma | -848.116 | 1812.233 | 0.402 | 0.074 | 0.111 | *N. discreta* |
| | GammaExpo | -748.921 | 1611.843 | 0.37 | 0.069 | 0.117 | *N. discreta* |
| | DisplGamma | -2971.691 | 6055.382 | 0 | 0 | 0.185 | *N. discreta* |
| | ScaledBeta | -689.795 | 1491.589 | 0.121 | 0.022 | 0.163 | *N. discreta* |
| | Average | NA | NA | 0.121 | 0.022 | 0.163 | *N. discreta* |
| All core genes | Neutral | -11855.775 | 23769.55 | 0.913 | 0.136 | 0.013 | *O. novo-ulmi subsp. americana* |
| | GammaZero | -1330.456 | 2720.913 | 0 | 0 | 0.149 | *O. novo-ulmi subsp. americana* |
| | GammaGamma | -229.56 | 525.119 | 0.476 | 0.071 | 0.078 | *O. novo-ulmi subsp. americana* |
| | GammaExpo | -245.643 | 555.286 | 0.603 | 0.09 | 0.059 | *O. novo-ulmi subsp. americana* |
| | DisplGamma | -1638.897 | 3339.794 | 0 | 0 | 0.149 | *O. novo-ulmi subsp. americana* |
| | ScaledBeta | -230.739 | 523.478 | 0.503 | 0.075 | 0.074 | *O. novo-ulmi subsp. americana* |
| | Average | NA | NA | 0.494 | 0.074 | 0.075 | *O. novo-ulmi subsp. americana* |
| All core genes | Neutral | -27159.832 | 54379.664 | 0.918 | 0.137 | 0.012 | *O. novo-ulmi subsp. novo-ulmi* |
| | GammaZero | -2996.697 | 6055.395 | 0 | 0 | 0.15 | *O. novo-ulmi subsp. novo-ulmi* |
| | GammaGamma | -202.028 | 472.057 | 0.721 | 0.108 | 0.042 | *O. novo-ulmi subsp. novo-ulmi* |
| | GammaExpo | -205.854 | 477.708 | 0.728 | 0.109 | 0.041 | *O. novo-ulmi subsp. novo-ulmi* |
| | DisplGamma | -3629.17 | 7322.34 | 0 | 0 | 0.15 | *O. novo-ulmi subsp. novo-ulmi* |
| | ScaledBeta | -248.976 | 561.952 | 0.565 | 0.085 | 0.065 | *O. novo-ulmi subsp. novo-ulmi* |
| | Average | NA | NA | 0.721 | 0.108 | 0.042 | *O. novo-ulmi subsp. novo-ulmi* |
| All core genes | Neutral | -82848.6 | 165775.2 | 0.915 | 0.139 | 0.013 | *O. ulmi* |
| | GammaZero | -7550.684 | 15181.367 | 0 | 0 | 0.152 | *O. ulmi* |
| | GammaGamma | -335.754 | 757.507 | 0.549 | 0.083 | 0.068 | *O. ulmi* |
| | GammaExpo | -335.749 | 755.498 | 0.493 | 0.075 | 0.077 | *O. ulmi* |
| | DisplGamma | -7550.684 | 15183.367 | 0 | 0 | 0.152 | *O. ulmi* |
| | ScaledBeta | -370.048 | 822.095 | 0.478 | 0.073 | 0.079 | *O. ulmi* |
| | Average | NA | NA | 0.508 | 0.077 | 0.075 | *O. ulmi* |
| All core genes | Neutral | -4304.107 | 8668.213 | 0.871 | 0.144 | 0.021 | *P. biforme* |
| | GammaZero | -638.856 | 1339.712 | 0 | 0 | 0.165 | *P. biforme* |

| | | | | | | | |
|---|---|---|---|---|---|---|---|
| | GammaGamma | -212.071 | 492.142 | 0.706 | 0.117 | 0.049 | *P. biforme* |
| | GammaExpo | -213.729 | 493.458 | 0.745 | 0.123 | 0.042 | *P. biforme* |
| | DisplGamma | -730.414 | 1524.828 | 0 | 0 | 0.165 | *P. biforme* |
| | ScaledBeta | -285.658 | 635.317 | 0.415 | 0.069 | 0.097 | *P. biforme* |
| | Average | NA | NA | 0.72 | 0.119 | 0.046 | *P. biforme* |
| All core genes | Neutral | -3712.838 | 8145.675 | 0.833 | 0.126 | 0.025 | *P. nodorum* |
| | GammaZero | -1663.787 | 4049.574 | 0 | 0 | 0.152 | *P. nodorum* |
| | GammaGamma | -1157.999 | 3043.999 | 0.834 | 0.127 | 0.025 | *P. nodorum* |
| | GammaExpo | -1184.315 | 3094.63 | 0.726 | 0.11 | 0.042 | *P. nodorum* |
| | DisplGamma | -1775.679 | 4275.358 | 0 | 0 | 0.152 | *P. nodorum* |
| | ScaledBeta | -1252.35 | 3228.7 | 0.914 | 0.139 | 0.013 | *P. nodorum* |
| | Average | NA | NA | 0.834 | 0.127 | 0.025 | *P. nodorum* |
| All core genes | Neutral | -17529.809 | 35365.617 | 0.987 | 0.243 | 0.003 | *P. teres f. teres* |
| | GammaZero | -802.092 | 1912.185 | 0 | 0 | 0.246 | *P. teres f. teres* |
| | GammaGamma | -802.326 | 1918.652 | 0 | 0 | 0.246 | *P. teres f. teres* |
| | GammaExpo | -802.092 | 1916.185 | 0 | 0 | 0.246 | *P. teres f. teres* |
| | DisplGamma | -800.983 | 1911.966 | 0 | 0 | 0.246 | *P. teres f. teres* |
| | ScaledBeta | -838.791 | 1987.582 | 0.271 | 0.067 | 0.179 | *P. teres f. teres* |
| | Average | NA | NA | 0 | 0 | 0.246 | *P. teres f. teres* |
| All core genes | Neutral | -6214.243 | 12672.487 | 0.615 | 0.097 | 0.061 | *R. commune* |
| | GammaZero | -846.055 | 1938.109 | 0 | 0 | 0.158 | *R. commune* |
| | GammaGamma | -855.695 | 1963.39 | 0 | 0 | 0.158 | *R. commune* |
| | GammaExpo | -846.056 | 1942.111 | 0 | 0 | 0.158 | *R. commune* |
| | DisplGamma | -798.271 | 1844.542 | 0 | 0 | 0.158 | *R. commune* |
| | ScaledBeta | -920.463 | 2088.926 | 0.217 | 0.034 | 0.123 | *R. commune* |
| | Average | NA | NA | 0 | 0 | 0.158 | *R. commune* |
| All core genes | Neutral | -786993.584 | 1574235.168 | 0.973 | 0.183 | 0.005 | *S. musiva* |
| | GammaZero | -42263.08 | 84776.16 | 0 | 0 | 0.188 | *S. musiva* |
| | GammaGamma | -1063.552 | 2383.104 | 0.659 | 0.124 | 0.064 | *S. musiva* |
| | GammaExpo | -1115.123 | 2484.246 | 0.714 | 0.134 | 0.054 | *S. musiva* |
| | DisplGamma | -42263.08 | 84778.16 | 0 | 0 | 0.188 | *S. musiva* |
| | ScaledBeta | -1770.879 | 3793.758 | 0.791 | 0.149 | 0.039 | *S. musiva* |
| | Average | NA | NA | 0.659 | 0.124 | 0.064 | *S. musiva* |
| All core genes | Neutral | -287746.085 | 575544.17 | 0.941 | 0.167 | 0.011 | *S. sclerotiorum* |
| | GammaZero | -5106.447 | 10266.895 | 0 | 0 | 0.178 | *S. sclerotiorum* |
| | GammaGamma | -541.994 | 1143.989 | 0 | 0 | 0.178 | *S. sclerotiorum* |
| | GammaExpo | -479.61 | 1017.219 | 0.002 | 0 | 0.177 | *S. sclerotiorum* |
| | DisplGamma | -5106.447 | 10268.895 | 0 | 0 | 0.178 | *S. sclerotiorum* |
| | ScaledBeta | -720.263 | 1496.527 | 0.338 | 0.06 | 0.117 | *S. sclerotiorum* |
| | Average | NA | NA | 0.002 | 0 | 0.177 | *S. sclerotiorum* |
| All core genes | Neutral | -5378.129 | 10970.258 | 0.928 | 0.141 | 0.011 | *V. dahliae* |
| | GammaZero | -382.514 | 981.028 | 0 | 0 | 0.152 | *V. dahliae* |
| | GammaGamma | -386.387 | 994.773 | 0 | 0 | 0.152 | *V. dahliae* |
| | GammaExpo | -382.553 | 985.106 | 0 | 0 | 0.152 | *V. dahliae* |
| | DisplGamma | -381.76 | 981.521 | 0 | 0 | 0.152 | *V. dahliae* |
| | ScaledBeta | -401.21 | 1020.42 | 0.029 | 0.004 | 0.148 | *V. dahliae* |
| | Average | NA | NA | 0 | 0 | 0.152 | *V. dahliae* |
| All core genes | Neutral | -303340.299 | 606850.598 | 0.954 | 0.166 | 0.008 | *V. inaequalis* |
| | GammaZero | -51833.991 | 103839.982 | 0 | 0 | 0.174 | *V. inaequalis* |
| | GammaGamma | -1323.631 | 2825.263 | 0.761 | 0.132 | 0.042 | *V. inaequalis* |
| | GammaExpo | -1541.682 | 3259.364 | 0.698 | 0.121 | 0.052 | *V. inaequalis* |
| | DisplGamma | -51833.991 | 103841.982 | 0 | 0 | 0.174 | *V. inaequalis* |
| | ScaledBeta | -2603.723 | 5381.446 | 0.945 | 0.164 | 0.01 | *V. inaequalis* |
| | Average | NA | NA | 0.761 | 0.132 | 0.042 | *V. inaequalis* |
| All core genes | Neutral | -103852.291 | 208678.581 | 0.908 | 0.111 | 0.011 | *Z. tritici* |
| | GammaZero | -5785.255 | 12546.51 | 0 | 0 | 0.122 | *Z. tritici* |
| | GammaGamma | -3272.357 | 7526.715 | 0.604 | 0.074 | 0.048 | *Z. tritici* |
| | GammaExpo | -3280.071 | 7540.143 | 0.46 | 0.056 | 0.066 | *Z. tritici* |
| | DisplGamma | -5785.389 | 12548.778 | 0 | 0 | 0.122 | *Z. tritici* |

| | | | | | | |
|---|---|---|---|---|---|---|
| ScaledBeta | -4432.399 | 9842.797 | 0.711 | 0.087 | 0.035 | *Z. tritici* |
| Average | NA | NA | 0.604 | 0.074 | 0.048 | *Z. tritici* |

**Table S10.** Selected models based on Akaike's information criterion (AIC) per species. Summarized from Table S4.

| Data | model | Log likelihood | AIC | alpha | omegaA | omegaNA | Species |
|---|---|---|---|---|---|---|---|
| All core genes | GammaExpo | -814.796 | 1827.593 | 0.148 | 0.024 | 0.136 | *A. flavus* |
| | GammaExpo | -764.182 | 1794.363 | 0.462 | 0.114 | 0.132 | *B. cinera* |
| | GammaExpo | -1826.09 | 4086.18 | 0.486 | 0.06 | 0.064 | *C. beticola* |
| | GammaZero | -845.677 | 2159.354 | 0 | 0 | 0.18 | *C. parasitica* |
| | DisplGamma | -520.959 | 1169.918 | 0 | 0 | 0.122 | *F. graminearum* |
| | DisplGamma | -431.796 | 1087.591 | 0 | 0 | 0.321 | *M. oryzae rice* |
| | GammaExpo | -175.836 | 397.672 | 0.294 | 0.07 | 0.17 | *M. oryzae Triticum* |
| | ScaledBeta | -689.795 | 1491.589 | 0.121 | 0.022 | 0.163 | *N. discreta* |
| | ScaledBeta | -230.739 | 523.478 | 0.503 | 0.075 | 0.074 | *O. novo-ulmi subsp. americana* |
| | GammaGamma | -202.028 | 472.057 | 0.721 | 0.108 | 0.042 | *O. novo-ulmi subsp. novo-ulmi* |
| | GammaExpo | -335.749 | 755.498 | 0.493 | 0.075 | 0.077 | *O. ulmi* |
| | GammaGamma | -212.071 | 492.142 | 0.706 | 0.117 | 0.049 | *P. biforme* |
| | GammaGamma | -1157.999 | 3043.999 | 0.834 | 0.127 | 0.025 | *P. nodorum* |
| | DisplGamma | -800.983 | 1911.966 | 0 | 0 | 0.246 | *P. teres f. teres* |
| | DisplGamma | -798.271 | 1844.542 | 0 | 0 | 0.158 | *R. commune* |
| | GammaGamma | -1063.552 | 2383.104 | 0.659 | 0.124 | 0.064 | *S. musiva* |
| | GammaExpo | -479.61 | 1017.219 | 0.002 | 0 | 0.177 | *S. sclerotiorum* |
| | GammaZero | -382.514 | 981.028 | 0 | 0 | 0.152 | *V. dahliae* |
| | GammaGamma | -1323.631 | 2825.263 | 0.761 | 0.132 | 0.042 | *V. inaequalis* |
| | GammaGamma | -3272.357 | 7526.715 | 0.604 | 0.074 | 0.048 | *Z. tritici* |

**Table S11.** Estimations of the rates of adaptation across non-secreted, secreted and effector gene categories per species. Akaike's information criterion (AIC).

| Data | model | Log likelihood | AIC | alpha | omegaA | omegaNA | Species |
|---|---|---|---|---|---|---|---|
| Non-secreted | Neutral | -37732.741 | 75657.482 | 0.901 | 0.142 | 0.016 | *A. flavus* |
| Non-secreted | GammaZero | -813.138 | 1820.276 | 0 | 0 | 0.157 | *A. flavus* |
| Non-secreted | GammaGamma | -798.525 | 1797.05 | 0.12 | 0.019 | 0.139 | *A. flavus* |
| Non-secreted | GammaExpo | -799.673 | 1797.347 | 0.124 | 0.02 | 0.138 | *A. flavus* |
| Non-secreted | DisplGamma | -848.975 | 1893.95 | 0 | 0 | 0.157 | *A. flavus* |
| Non-secreted | ScaledBeta | -819.868 | 1835.736 | 0.061 | 0.01 | 0.148 | *A. flavus* |
| Non-secreted | Average | NA | NA | 0.122 | 0.019 | 0.138 | *A. flavus* |
| Secreted | Neutral | -3119.838 | 6431.676 | 0.892 | 0.15 | 0.018 | *A. flavus* |
| Secreted | GammaZero | -478.194 | 1150.388 | 0 | 0 | 0.168 | *A. flavus* |
| Secreted | GammaGamma | -466.933 | 1133.866 | 0.246 | 0.041 | 0.127 | *A. flavus* |
| Secreted | GammaExpo | -464.758 | 1127.515 | 0.29 | 0.049 | 0.119 | *A. flavus* |
| Secreted | DisplGamma | -488.082 | 1172.165 | 0 | 0 | 0.168 | *A. flavus* |
| Secreted | ScaledBeta | -469.741 | 1135.483 | 0.3 | 0.05 | 0.118 | *A. flavus* |
| Secreted | Average | NA | NA | 0.288 | 0.048 | 0.12 | *A. flavus* |
| Effectors | Neutral | -894.109 | 1980.217 | 0.865 | 0.201 | 0.031 | *A. flavus* |
| Effectors | GammaZero | -336.429 | 866.858 | 0 | 0 | 0.232 | *A. flavus* |
| Effectors | GammaGamma | -328.834 | 857.667 | 0 | 0 | 0.232 | *A. flavus* |
| Effectors | GammaExpo | -329.063 | 856.126 | 0.223 | 0.052 | 0.18 | *A. flavus* |
| Effectors | DisplGamma | -337.445 | 870.89 | 0 | 0 | 0.232 | *A. flavus* |
| Effectors | ScaledBeta | -330.501 | 857.002 | 0.269 | 0.062 | 0.17 | *A. flavus* |
| Effectors | Average | NA | NA | 0.188 | 0.043 | 0.188 | *A. flavus* |
| Non-secreted | Neutral | -37148.175 | 74556.349 | 0.988 | 0.242 | 0.003 | *B. cinera* |
| Non-secreted | GammaZero | -1139.891 | 2541.782 | 0 | 0 | 0.245 | *B. cinera* |
| Non-secreted | GammaGamma | -758.519 | 1785.039 | 0.455 | 0.112 | 0.134 | *B. cinera* |
| Non-secreted | GammaExpo | -758.843 | 1783.685 | 0.47 | 0.115 | 0.13 | *B. cinera* |
| Non-secreted | DisplGamma | -1110.879 | 2485.758 | 0 | 0 | 0.245 | *B. cinera* |
| Non-secreted | ScaledBeta | -785.521 | 1835.042 | 0.377 | 0.093 | 0.153 | *B. cinera* |
| Non-secreted | Average | NA | NA | 0.465 | 0.114 | 0.131 | *B. cinera* |
| Secreted | Neutral | -1567.094 | 3394.188 | 0.984 | 0.242 | 0.004 | *B. cinera* |
| Secreted | GammaZero | -293.254 | 848.508 | 0 | 0 | 0.245 | *B. cinera* |
| Secreted | GammaGamma | -286.683 | 841.366 | 0.388 | 0.095 | 0.15 | *B. cinera* |
| Secreted | GammaExpo | -286.96 | 839.92 | 0.392 | 0.096 | 0.149 | *B. cinera* |
| Secreted | DisplGamma | -293.747 | 851.494 | 0 | 0 | 0.245 | *B. cinera* |
| Secreted | ScaledBeta | -295.58 | 855.16 | 0 | 0 | 0.245 | *B. cinera* |
| Secreted | Average | NA | NA | 0.386 | 0.095 | 0.151 | *B. cinera* |
| Effectors | Neutral | -673.719 | 1607.437 | 0.982 | 0.289 | 0.005 | *B. cinera* |
| Effectors | GammaZero | -187.029 | 636.057 | 0 | 0 | 0.294 | *B. cinera* |
| Effectors | GammaGamma | -181.644 | 631.288 | 0.561 | 0.165 | 0.129 | *B. cinera* |
| Effectors | GammaExpo | -181.726 | 629.451 | 0.477 | 0.141 | 0.154 | *B. cinera* |
| Effectors | DisplGamma | -186.874 | 637.747 | 0 | 0 | 0.294 | *B. cinera* |
| Effectors | ScaledBeta | -181.635 | 627.27 | 0.304 | 0.089 | 0.205 | *B. cinera* |
| Effectors | Average | NA | NA | 0.363 | 0.107 | 0.188 | *B. cinera* |
| Non-secreted | Neutral | -80089.912 | 160607.823 | 0.936 | 0.116 | 0.008 | *C. beticola* |
| Non-secreted | GammaZero | -2850.488 | 6130.975 | 0 | 0 | 0.124 | *C. beticola* |
| Non-secreted | GammaGamma | -1811.527 | 4059.055 | 0.49 | 0.06 | 0.063 | *C. beticola* |
| Non-secreted | GammaExpo | -1795.868 | 4025.735 | 0.482 | 0.06 | 0.064 | *C. beticola* |
| Non-secreted | DisplGamma | -3550.958 | 7533.916 | 0 | 0 | 0.124 | *C. beticola* |
| Non-secreted | ScaledBeta | -2179.34 | 4790.68 | 0.327 | 0.04 | 0.083 | *C. beticola* |
| Non-secreted | Average | NA | NA | 0.482 | 0.06 | 0.064 | *C. beticola* |
| Secreted | Neutral | -4271.803 | 8971.606 | 0.924 | 0.117 | 0.01 | *C. beticola* |
| Secreted | GammaZero | -931.661 | 2293.321 | 0 | 0 | 0.126 | *C. beticola* |
| Secreted | GammaGamma | -869.82 | 2175.64 | 0.484 | 0.061 | 0.065 | *C. beticola* |
| Secreted | GammaExpo | -869.459 | 2172.919 | 0.511 | 0.064 | 0.062 | *C. beticola* |
| Secreted | DisplGamma | -948.465 | 2328.929 | 0 | 0 | 0.126 | *C. beticola* |
| Secreted | ScaledBeta | -884.313 | 2200.627 | 0.313 | 0.04 | 0.087 | *C. beticola* |

| | | | | | | | |
|---|---|---|---|---|---|---|---|
| Secreted | Average | NA | NA | 0.506 | 0.064 | 0.062 | *C. beticola* |
| Effectors | Neutral | -1296.084 | 3020.168 | 0.92 | 0.148 | 0.013 | *C. beticola* |
| Effectors | GammaZero | -490.923 | 1411.846 | 0 | 0 | 0.161 | *C. beticola* |
| Effectors | GammaGamma | -471.321 | 1378.642 | 0.542 | 0.087 | 0.074 | *C. beticola* |
| Effectors | GammaExpo | -470.608 | 1375.217 | 0.503 | 0.081 | 0.08 | *C. beticola* |
| Effectors | DisplGamma | -494.719 | 1421.438 | 0 | 0 | 0.161 | *C. beticola* |
| Effectors | ScaledBeta | -475.708 | 1383.415 | 0.351 | 0.056 | 0.104 | *C. beticola* |
| Effectors | Average | NA | NA | 0.507 | 0.082 | 0.079 | *C. beticola* |
| Non-secreted | Neutral | -3566.294 | 7598.587 | 0.847 | 0.152 | 0.027 | *C. parasitica* |
| Non-secreted | GammaZero | -834.344 | 2136.689 | 0 | 0 | 0.179 | *C. parasitica* |
| Non-secreted | GammaGamma | -855.783 | 2185.566 | 0.015 | 0.003 | 0.177 | *C. parasitica* |
| Non-secreted | GammaExpo | -846.796 | 2165.592 | 0.098 | 0.018 | 0.162 | *C. parasitica* |
| Non-secreted | DisplGamma | -835.803 | 2141.606 | 0 | 0 | 0.179 | *C. parasitica* |
| Non-secreted | ScaledBeta | -891.97 | 2253.941 | 0.245 | 0.044 | 0.135 | *C. parasitica* |
| Non-secreted | Average | NA | NA | 0 | 0 | 0.179 | *C. parasitica* |
| Secreted | Neutral | -305.867 | 1077.734 | 0.849 | 0.152 | 0.027 | *C. parasitica* |
| Secreted | GammaZero | -178.494 | 824.989 | 0 | 0 | 0.179 | *C. parasitica* |
| Secreted | GammaGamma | -180.102 | 834.205 | 0.408 | 0.073 | 0.106 | *C. parasitica* |
| Secreted | GammaExpo | -179.312 | 830.623 | 0.221 | 0.04 | 0.139 | *C. parasitica* |
| Secreted | DisplGamma | -178.984 | 827.968 | 0 | 0 | 0.179 | *C. parasitica* |
| Secreted | ScaledBeta | -180.912 | 831.825 | 0.347 | 0.062 | 0.117 | *C. parasitica* |
| Secreted | Average | NA | NA | 0.022 | 0.004 | 0.175 | *C. parasitica* |
| Effectors | Neutral | -116.987 | 699.973 | 0.86 | 0.191 | 0.031 | *C. parasitica* |
| Effectors | GammaZero | -76.637 | 621.274 | 0 | 0 | 0.222 | *C. parasitica* |
| Effectors | GammaGamma | -76.181 | 626.363 | 0.718 | 0.16 | 0.063 | *C. parasitica* |
| Effectors | GammaExpo | -76.304 | 624.608 | 0.601 | 0.134 | 0.089 | *C. parasitica* |
| Effectors | DisplGamma | -76.755 | 623.51 | 0 | 0 | 0.222 | *C. parasitica* |
| Effectors | ScaledBeta | -76.89 | 623.78 | 0.268 | 0.06 | 0.163 | *C. parasitica* |
| Effectors | Average | NA | NA | 0.131 | 0.029 | 0.193 | *C. parasitica* |
| Non-secreted | Neutral | -59033.247 | 118190.495 | 0.952 | 0.116 | 0.006 | *F. graminearum* |
| Non-secreted | GammaZero | -517.958 | 1161.916 | 0 | 0 | 0.122 | *F. graminearum* |
| Non-secreted | GammaGamma | -517.961 | 1167.922 | 0 | 0 | 0.122 | *F. graminearum* |
| Non-secreted | GammaExpo | -517.96 | 1165.92 | 0 | 0 | 0.122 | *F. graminearum* |
| Non-secreted | DisplGamma | -508.617 | 1145.234 | 0 | 0 | 0.122 | *F. graminearum* |
| Non-secreted | ScaledBeta | -543.56 | 1215.12 | 0.012 | 0.001 | 0.121 | *F. graminearum* |
| Non-secreted | Average | NA | NA | 0 | 0 | 0.122 | *F. graminearum* |
| Secreted | Neutral | -5457.743 | 11039.487 | 0.947 | 0.11 | 0.006 | *F. graminearum* |
| Secreted | GammaZero | -349.028 | 824.056 | 0 | 0 | 0.116 | *F. graminearum* |
| Secreted | GammaGamma | -344.938 | 821.876 | 0.148 | 0.017 | 0.099 | *F. graminearum* |
| Secreted | GammaExpo | -344.868 | 819.735 | 0.174 | 0.02 | 0.096 | *F. graminearum* |
| Secreted | DisplGamma | -351.894 | 831.788 | 0 | 0 | 0.116 | *F. graminearum* |
| Secreted | ScaledBeta | -348.478 | 824.956 | 0.128 | 0.015 | 0.101 | *F. graminearum* |
| Secreted | Average | NA | NA | 0.152 | 0.018 | 0.098 | *F. graminearum* |
| Effectors | Neutral | -2195.677 | 4515.354 | 0.957 | 0.141 | 0.006 | *F. graminearum* |
| Effectors | GammaZero | -288.627 | 703.255 | 0 | 0 | 0.147 | *F. graminearum* |
| Effectors | GammaGamma | -280.949 | 693.899 | 0.332 | 0.049 | 0.098 | *F. graminearum* |
| Effectors | GammaExpo | -282.218 | 694.435 | 0.368 | 0.054 | 0.093 | *F. graminearum* |
| Effectors | DisplGamma | -297.809 | 723.618 | 0 | 0 | 0.147 | *F. graminearum* |
| Effectors | ScaledBeta | -282.272 | 692.543 | 0.21 | 0.031 | 0.116 | *F. graminearum* |
| Effectors | Average | NA | NA | 0.274 | 0.04 | 0.107 | *F. graminearum* |
| Non-secreted | Neutral | -14530.107 | 29280.214 | 0.996 | 0.322 | 0.001 | *M. oryzae rice* |
| Non-secreted | GammaZero | -434.168 | 1090.336 | 0 | 0 | 0.324 | *M. oryzae rice* |
| Non-secreted | GammaGamma | -434.168 | 1096.336 | 0 | 0 | 0.324 | *M. oryzae rice* |
| Non-secreted | GammaExpo | -434.168 | 1094.336 | 0 | 0 | 0.324 | *M. oryzae rice* |
| Non-secreted | DisplGamma | -426.102 | 1076.204 | 0 | 0 | 0.324 | *M. oryzae rice* |
| Non-secreted | ScaledBeta | -480.743 | 1185.486 | 0.281 | 0.091 | 0.233 | *M. oryzae rice* |
| Non-secreted | Average | NA | NA | 0 | 0 | 0.324 | *M. oryzae rice* |
| Secreted | Neutral | -857.587 | 1935.174 | 0.993 | 0.272 | 0.002 | *M. oryzae rice* |
| Secreted | GammaZero | -150.911 | 523.823 | 0 | 0 | 0.274 | *M. oryzae rice* |
| Secreted | GammaGamma | -150.911 | 529.822 | 0 | 0 | 0.274 | *M. oryzae rice* |

| | | | | | | | |
|---|---|---|---|---|---|---|---|
| Secreted | GammaExpo | -150.911 | 527.823 | 0 | 0 | 0.274 | *M. oryzae rice* |
| Secreted | DisplGamma | -150.476 | 524.952 | 0 | 0 | 0.274 | *M. oryzae rice* |
| Secreted | ScaledBeta | -156.335 | 536.67 | 0.332 | 0.091 | 0.183 | *M. oryzae rice* |
| Secreted | Average | NA | NA | 0 | 0 | 0.274 | *M. oryzae rice* |
| Effectors | Neutral | -407.712 | 1035.424 | 0.995 | 0.372 | 0.002 | *M. oryzae rice* |
| Effectors | GammaZero | -83.433 | 388.867 | 0 | 0 | 0.374 | *M. oryzae rice* |
| Effectors | GammaGamma | -83.198 | 394.396 | 0 | 0 | 0.374 | *M. oryzae rice* |
| Effectors | GammaExpo | -83.267 | 392.534 | 0.184 | 0.069 | 0.305 | *M. oryzae rice* |
| Effectors | DisplGamma | -83.433 | 390.867 | 0 | 0 | 0.374 | *M. oryzae rice* |
| Effectors | ScaledBeta | -83.924 | 391.849 | 0.408 | 0.153 | 0.222 | *M. oryzae rice* |
| Effectors | Average | NA | NA | 0.067 | 0.025 | 0.349 | *M. oryzae rice* |
| Non-secreted | Neutral | -63159.796 | 126359.593 | 0.996 | 0.24 | 0.001 | *M. oryzae Triticum* |
| Non-secreted | GammaZero | -391.615 | 825.23 | 0 | 0 | 0.241 | *M. oryzae Triticum* |
| Non-secreted | GammaGamma | -173.626 | 395.252 | 0.26 | 0.062 | 0.178 | *M. oryzae Triticum* |
| Non-secreted | GammaExpo | -173.567 | 393.135 | 0.283 | 0.068 | 0.173 | *M. oryzae Triticum* |
| Non-secreted | DisplGamma | -400.767 | 845.535 | 0 | 0 | 0.241 | *M. oryzae Triticum* |
| Non-secreted | ScaledBeta | -197.105 | 438.211 | 0.312 | 0.075 | 0.166 | *M. oryzae Triticum* |
| Non-secreted | Average | NA | NA | 0.277 | 0.067 | 0.174 | *M. oryzae Triticum* |
| Secreted | Neutral | -5563.245 | 11166.49 | 0.994 | 0.208 | 0.001 | *M. oryzae Triticum* |
| Secreted | GammaZero | -128.892 | 299.783 | 0 | 0 | 0.209 | *M. oryzae Triticum* |
| Secreted | GammaGamma | -116.388 | 280.775 | 0 | 0 | 0.209 | *M. oryzae Triticum* |
| Secreted | GammaExpo | -116.88 | 279.759 | 0.222 | 0.046 | 0.163 | *M. oryzae Triticum* |
| Secreted | DisplGamma | -131.384 | 306.768 | 0 | 0 | 0.209 | *M. oryzae Triticum* |
| Secreted | ScaledBeta | -119.162 | 282.324 | 0.207 | 0.043 | 0.166 | *M. oryzae Triticum* |
| Secreted | Average | NA | NA | 0.148 | 0.031 | 0.178 | *M. oryzae Triticum* |
| Effectors | Neutral | -1502.785 | 3045.571 | 0.99 | 0.314 | 0.003 | *M. oryzae Triticum* |
| Effectors | GammaZero | -105.314 | 252.628 | 0 | 0 | 0.317 | *M. oryzae Triticum* |
| Effectors | GammaGamma | -99.43 | 246.859 | 0 | 0 | 0.317 | *M. oryzae Triticum* |
| Effectors | GammaExpo | -99.284 | 244.569 | 0.035 | 0.011 | 0.306 | *M. oryzae Triticum* |
| Effectors | DisplGamma | -104.588 | 253.176 | 0 | 0 | 0.317 | *M. oryzae Triticum* |
| Effectors | ScaledBeta | -99.201 | 242.402 | 0.013 | 0.004 | 0.313 | *M. oryzae Triticum* |
| Effectors | Average | NA | NA | 0.017 | 0.005 | 0.312 | *M. oryzae Triticum* |
| Non-secreted | Neutral | -123403.843 | 246915.686 | 0.937 | 0.171 | 0.011 | *N. discreta* |
| Non-secreted | GammaZero | -2722.498 | 5554.995 | 0 | 0 | 0.182 | *N. discreta* |
| Non-secreted | GammaGamma | -720.817 | 1557.634 | 0.329 | 0.06 | 0.122 | *N. discreta* |
| Non-secreted | GammaExpo | -735.311 | 1584.623 | 0.362 | 0.066 | 0.116 | *N. discreta* |
| Non-secreted | DisplGamma | -2722.498 | 5556.995 | 0 | 0 | 0.182 | *N. discreta* |
| Non-secreted | ScaledBeta | -774.808 | 1661.617 | 0.328 | 0.06 | 0.123 | *N. discreta* |
| Non-secreted | Average | NA | NA | 0.329 | 0.06 | 0.122 | *N. discreta* |

| | | | | | | | |
|---|---|---|---|---|---|---|---|
| Secreted | Neutral | -3772.602 | 7653.203 | 0.919 | 0.225 | 0.02 | *N. discreta* |
| Secreted | GammaZero | -372.25 | 854.5 | 0 | 0 | 0.245 | *N. discreta* |
| Secreted | GammaGamma | -257.573 | 631.145 | 0.424 | 0.104 | 0.141 | *N. discreta* |
| Secreted | GammaExpo | -257.624 | 629.248 | 0.429 | 0.105 | 0.14 | *N. discreta* |
| Secreted | DisplGamma | -348.235 | 808.469 | 0 | 0 | 0.245 | *N. discreta* |
| Secreted | ScaledBeta | -256.343 | 624.687 | 0.384 | 0.094 | 0.151 | *N. discreta* |
| Secreted | Average | NA | NA | 0.389 | 0.095 | 0.149 | *N. discreta* |
| Effectors | Neutral | -1183.529 | 2475.059 | 0.927 | 0.249 | 0.02 | *N. discreta* |
| Effectors | GammaZero | -220.453 | 550.906 | 0 | 0 | 0.269 | *N. discreta* |
| Effectors | GammaGamma | -158.905 | 433.811 | 0.568 | 0.153 | 0.116 | *N. discreta* |
| Effectors | GammaExpo | -158.315 | 430.629 | 0.554 | 0.149 | 0.12 | *N. discreta* |
| Effectors | DisplGamma | -210.625 | 533.25 | 0 | 0 | 0.269 | *N. discreta* |
| Effectors | ScaledBeta | -161.106 | 434.212 | 0.476 | 0.128 | 0.141 | *N. discreta* |
| Effectors | Average | NA | NA | 0.547 | 0.147 | 0.122 | *N. discreta* |
| Non-secreted | Neutral | -11467.161 | 22992.322 | 0.914 | 0.135 | 0.013 | *O. novo-ulmi subsp. americana* |
| Non-secreted | GammaZero | -1279.29 | 2618.581 | 0 | 0 | 0.148 | *O. novo-ulmi subsp. americana* |
| Non-secreted | GammaGamma | -227.628 | 521.256 | 0.492 | 0.073 | 0.075 | *O. novo-ulmi subsp. americana* |
| Non-secreted | GammaExpo | -237.748 | 539.496 | 0.579 | 0.086 | 0.062 | *O. novo-ulmi subsp. americana* |
| Non-secreted | DisplGamma | -1641.715 | 3345.43 | 0 | 0 | 0.148 | *O. novo-ulmi subsp. americana* |
| Non-secreted | ScaledBeta | -236.715 | 535.431 | 0.481 | 0.071 | 0.077 | *O. novo-ulmi subsp. americana* |
| Non-secreted | Average | NA | NA | 0.492 | 0.073 | 0.075 | *O. novo-ulmi subsp. americana* |
| Secreted | Neutral | -423.663 | 905.325 | 0.913 | 0.172 | 0.016 | *O. novo-ulmi subsp. americana* |
| Secreted | GammaZero | -145.35 | 350.7 | 0 | 0 | 0.189 | *O. novo-ulmi subsp. americana* |
| Secreted | GammaGamma | -99.391 | 264.782 | 0.77 | 0.145 | 0.043 | *O. novo-ulmi subsp. americana* |
| Secreted | GammaExpo | -99.314 | 262.629 | 0.743 | 0.14 | 0.049 | *O. novo-ulmi subsp. americana* |
| Secreted | DisplGamma | -151.644 | 365.287 | 0 | 0 | 0.189 | *O. novo-ulmi subsp. americana* |
| Secreted | ScaledBeta | -100.05 | 262.101 | 0.566 | 0.107 | 0.082 | *O. novo-ulmi subsp. americana* |
| Secreted | Average | NA | NA | 0.659 | 0.124 | 0.064 | *O. novo-ulmi subsp. americana* |
| Effectors | Neutral | -139.296 | 336.592 | 0.869 | 0.18 | 0.027 | *O. novo-ulmi subsp. americana* |

| | | | | | | | |
|---|---|---|---|---|---|---|---|
| Effectors | GammaZero | -78.661 | 217.322 | 0 | 0 | 0.207 | *O. novo-ulmi subsp. americana* |
| Effectors | GammaGamma | -70.386 | 206.772 | 0.484 | 0.1 | 0.107 | *O. novo-ulmi subsp. americana* |
| Effectors | GammaExpo | -70.748 | 205.496 | 0.668 | 0.138 | 0.069 | *O. novo-ulmi subsp. americana* |
| Effectors | DisplGamma | -79.906 | 221.813 | 0 | 0 | 0.207 | *O. novo-ulmi subsp. americana* |
| Effectors | ScaledBeta | -70.725 | 203.45 | 0.47 | 0.097 | 0.11 | *O. novo-ulmi subsp. americana* |
| Effectors | Average | NA | NA | 0.517 | 0.107 | 0.1 | *O. novo-ulmi subsp. americana* |
| Non-secreted | Neutral | -26170.889 | 52401.777 | 0.918 | 0.136 | 0.012 | *O. novo-ulmi subsp. novo-ulmi* |
| Non-secreted | GammaZero | -2853.955 | 5769.91 | 0 | 0 | 0.148 | *O. novo-ulmi subsp. novo-ulmi* |
| Non-secreted | GammaGamma | -198.755 | 465.51 | 0.672 | 0.1 | 0.049 | *O. novo-ulmi subsp. novo-ulmi* |
| Non-secreted | GammaExpo | -205.347 | 476.695 | 0.739 | 0.11 | 0.039 | *O. novo-ulmi subsp. novo-ulmi* |
| Non-secreted | DisplGamma | -2853.955 | 5771.91 | 0 | 0 | 0.148 | *O. novo-ulmi subsp. novo-ulmi* |
| Non-secreted | ScaledBeta | -246.296 | 556.592 | 0.567 | 0.084 | 0.064 | *O. novo-ulmi subsp. novo-ulmi* |
| Non-secreted | Average | NA | NA | 0.672 | 0.1 | 0.049 | *O. novo-ulmi subsp. novo-ulmi* |
| Secreted | Neutral | -925.957 | 1911.914 | 0.917 | 0.175 | 0.016 | *O. novo-ulmi subsp. novo-ulmi* |
| Secreted | GammaZero | -238.132 | 538.264 | 0 | 0 | 0.191 | *O. novo-ulmi subsp. novo-ulmi* |
| Secreted | GammaGamma | -106.646 | 281.293 | 0.646 | 0.123 | 0.067 | *O. novo-ulmi subsp. novo-ulmi* |
| Secreted | GammaExpo | -107.508 | 281.016 | 0.76 | 0.145 | 0.046 | *O. novo-ulmi subsp. novo-ulmi* |
| Secreted | DisplGamma | -256.708 | 577.415 | 0 | 0 | 0.191 | *O. novo-ulmi subsp. novo-ulmi* |
| Secreted | ScaledBeta | -107.365 | 278.729 | 0.642 | 0.122 | 0.068 | *O. novo-ulmi subsp. novo-ulmi* |
| Secreted | Average | NA | NA | 0.667 | 0.127 | 0.064 | *O. novo-ulmi subsp. novo-ulmi* |

| | | | | | | | |
|---|---|---|---|---|---|---|---|
| Effectors | Neutral | -266.233 | 592.465 | 0.885 | 0.184 | 0.024 | *O. novo-ulmi subsp. novo-ulmi* |
| Effectors | GammaZero | -107.768 | 277.537 | 0 | 0 | 0.208 | *O. novo-ulmi subsp. novo-ulmi* |
| Effectors | GammaGamma | -81.86 | 231.72 | 0 | 0 | 0.208 | *O. novo-ulmi subsp. novo-ulmi* |
| Effectors | GammaExpo | -81.955 | 229.91 | 0.209 | 0.043 | 0.164 | *O. novo-ulmi subsp. novo-ulmi* |
| Effectors | DisplGamma | -109.159 | 282.317 | 0 | 0 | 0.208 | *O. novo-ulmi subsp. novo-ulmi* |
| Effectors | ScaledBeta | -83.145 | 230.289 | 0.562 | 0.117 | 0.091 | *O. novo-ulmi subsp. novo-ulmi* |
| Effectors | Average | NA | NA | 0.302 | 0.063 | 0.145 | *O. novo-ulmi subsp. novo-ulmi* |
| Non-secreted | Neutral | -80266.048 | 160610.095 | 0.916 | 0.138 | 0.013 | *O. ulmi* |
| Non-secreted | GammaZero | -7278.049 | 14636.098 | 0 | 0 | 0.15 | *O. ulmi* |
| Non-secreted | GammaGamma | -335.72 | 757.44 | 0.444 | 0.067 | 0.084 | *O. ulmi* |
| Non-secreted | GammaExpo | -338.858 | 761.715 | 0.544 | 0.082 | 0.068 | *O. ulmi* |
| Non-secreted | DisplGamma | -8751.928 | 17585.855 | 0 | 0 | 0.15 | *O. ulmi* |
| Non-secreted | ScaledBeta | -357.827 | 797.654 | 0.478 | 0.072 | 0.078 | *O. ulmi* |
| Non-secreted | Average | NA | NA | 0.454 | 0.068 | 0.082 | *O. ulmi* |
| Secreted | Neutral | -2266.798 | 4611.597 | 0.895 | 0.174 | 0.021 | *O. ulmi* |
| Secreted | GammaZero | -372.502 | 825.003 | 0 | 0 | 0.195 | *O. ulmi* |
| Secreted | GammaGamma | -145.685 | 377.37 | 0.148 | 0.029 | 0.166 | *O. ulmi* |
| Secreted | GammaExpo | -145.908 | 375.816 | 0.301 | 0.059 | 0.136 | *O. ulmi* |
| Secreted | DisplGamma | -397.134 | 876.269 | 0 | 0 | 0.195 | *O. ulmi* |
| Secreted | ScaledBeta | -158.787 | 399.574 | 0.5 | 0.098 | 0.098 | *O. ulmi* |
| Secreted | Average | NA | NA | 0.253 | 0.049 | 0.146 | *O. ulmi* |
| Effectors | Neutral | -598.97 | 1275.941 | 0.885 | 0.191 | 0.025 | *O. ulmi* |
| Effectors | GammaZero | -156.619 | 393.239 | 0 | 0 | 0.215 | *O. ulmi* |
| Effectors | GammaGamma | -96.974 | 279.948 | 0.476 | 0.103 | 0.113 | *O. ulmi* |
| Effectors | GammaExpo | -97.687 | 279.373 | 0.561 | 0.121 | 0.094 | *O. ulmi* |
| Effectors | DisplGamma | -161.917 | 405.835 | 0 | 0 | 0.215 | *O. ulmi* |
| Effectors | ScaledBeta | -98.154 | 278.307 | 0.491 | 0.106 | 0.11 | *O. ulmi* |
| Effectors | Average | NA | NA | 0.508 | 0.109 | 0.106 | *O. ulmi* |
| Non-secreted | Neutral | -4146.859 | 8353.719 | 0.872 | 0.143 | 0.021 | *P. biforme* |
| Non-secreted | GammaZero | -609.632 | 1281.263 | 0 | 0 | 0.164 | *P. biforme* |
| Non-secreted | GammaGamma | -210.767 | 489.533 | 0.71 | 0.116 | 0.047 | *P. biforme* |
| Non-secreted | GammaExpo | -211.979 | 489.958 | 0.738 | 0.121 | 0.043 | *P. biforme* |
| Non-secreted | DisplGamma | -691.506 | 1447.013 | 0 | 0 | 0.164 | *P. biforme* |
| Non-secreted | ScaledBeta | -248.085 | 560.171 | 0.84 | 0.138 | 0.026 | *P. biforme* |
| Non-secreted | Average | NA | NA | 0.723 | 0.118 | 0.045 | *P. biforme* |
| Secreted | Neutral | -242.893 | 545.787 | 0.868 | 0.174 | 0.026 | *P. biforme* |
| Secreted | GammaZero | -131.586 | 325.172 | 0 | 0 | 0.2 | *P. biforme* |
| Secreted | GammaGamma | -107.016 | 282.032 | 0.727 | 0.145 | 0.055 | *P. biforme* |
| Secreted | GammaExpo | -107.099 | 280.197 | 0.693 | 0.139 | 0.062 | *P. biforme* |
| Secreted | DisplGamma | -136.027 | 336.055 | 0 | 0 | 0.2 | *P. biforme* |
| Secreted | ScaledBeta | -108.644 | 281.289 | 0.567 | 0.113 | 0.087 | *P. biforme* |
| Secreted | Average | NA | NA | 0.663 | 0.133 | 0.068 | *P. biforme* |
| Effectors | Neutral | -74.555 | 209.11 | 0.817 | 0.215 | 0.048 | *P. biforme* |
| Effectors | GammaZero | -59.991 | 181.983 | 0 | 0 | 0.263 | *P. biforme* |
| Effectors | GammaGamma | -55.046 | 178.092 | 0.728 | 0.192 | 0.072 | *P. biforme* |
| Effectors | GammaExpo | -54.752 | 175.503 | 0.686 | 0.181 | 0.083 | *P. biforme* |

| | | | | | | | |
|---|---|---|---|---|---|---|---|
| Effectors | DisplGamma | -60.057 | 184.114 | 0 | 0 | 0.263 | *P. biforme* |
| Effectors | ScaledBeta | -56.653 | 177.306 | 0.93 | 0.245 | 0.018 | *P. biforme* |
| Effectors | Average | NA | NA | 0.729 | 0.192 | 0.071 | *P. biforme* |
| Non-secreted | Neutral | -3609.3 | 7938.6 | 0.834 | 0.124 | 0.025 | *P. nodorum* |
| Non-secreted | GammaZero | -1612.844 | 3947.688 | 0 | 0 | 0.149 | *P. nodorum* |
| Non-secreted | GammaGamma | -1141.867 | 3011.735 | 0.832 | 0.124 | 0.025 | *P. nodorum* |
| Non-secreted | GammaExpo | -1157.616 | 3041.232 | 0.754 | 0.112 | 0.037 | *P. nodorum* |
| Non-secreted | DisplGamma | -1784.872 | 4293.743 | 0 | 0 | 0.149 | *P. nodorum* |
| Non-secreted | ScaledBeta | -1172.725 | 3069.449 | 0.891 | 0.133 | 0.016 | *P. nodorum* |
| Non-secreted | Average | NA | NA | 0.832 | 0.124 | 0.025 | *P. nodorum* |
| Secreted | Neutral | -188.719 | 1097.437 | 0.792 | 0.157 | 0.041 | *P. nodorum* |
| Secreted | GammaZero | -154.929 | 1031.859 | 0 | 0 | 0.198 | *P. nodorum* |
| Secreted | GammaGamma | -136.554 | 1001.109 | 0.867 | 0.172 | 0.026 | *P. nodorum* |
| Secreted | GammaExpo | -137.892 | 1001.784 | 0.764 | 0.151 | 0.047 | *P. nodorum* |
| Secreted | DisplGamma | -157.278 | 1038.556 | 0 | 0 | 0.198 | *P. nodorum* |
| Secreted | ScaledBeta | -143.294 | 1010.588 | 0.929 | 0.184 | 0.014 | *P. nodorum* |
| Secreted | Average | NA | NA | 0.825 | 0.163 | 0.035 | *P. nodorum* |
| Effectors | Neutral | -151.024 | 1022.047 | 0.868 | 0.209 | 0.032 | *P. nodorum* |
| Effectors | GammaZero | -126.889 | 975.779 | 0 | 0 | 0.241 | *P. nodorum* |
| Effectors | GammaGamma | -110.45 | 948.9 | 0.809 | 0.195 | 0.046 | *P. nodorum* |
| Effectors | GammaExpo | -110.45 | 946.9 | 0.833 | 0.201 | 0.04 | *P. nodorum* |
| Effectors | DisplGamma | -125.79 | 975.58 | 0 | 0 | 0.241 | *P. nodorum* |
| Effectors | ScaledBeta | -109.344 | 942.688 | 0.934 | 0.225 | 0.016 | *P. nodorum* |
| Effectors | Average | NA | NA | 0.918 | 0.221 | 0.02 | *P. nodorum* |
| Non-secreted | Neutral | -16568.814 | 33443.627 | 0.987 | 0.245 | 0.003 | *P. teres f. teres* |
| Non-secreted | GammaZero | -787.015 | 1882.03 | 0 | 0 | 0.248 | *P. teres f. teres* |
| Non-secreted | GammaGamma | -787.029 | 1888.058 | 0 | 0 | 0.248 | *P. teres f. teres* |
| Non-secreted | GammaExpo | -787.233 | 1886.466 | 0.005 | 0.001 | 0.246 | *P. teres f. teres* |
| Non-secreted | DisplGamma | -786.155 | 1882.31 | 0 | 0 | 0.248 | *P. teres f. teres* |
| Non-secreted | ScaledBeta | -787.545 | 1885.09 | 0.04 | 0.01 | 0.238 | *P. teres f. teres* |
| Non-secreted | Average | NA | NA | 0.004 | 0.001 | 0.247 | *P. teres f. teres* |
| Secreted | Neutral | -1000.067 | 2306.134 | 0.98 | 0.209 | 0.004 | *P. teres f. teres* |
| Secreted | GammaZero | -268.513 | 845.026 | 0 | 0 | 0.213 | *P. teres f. teres* |
| Secreted | GammaGamma | -268.513 | 851.026 | 0 | 0 | 0.213 | *P. teres f. teres* |
| Secreted | GammaExpo | -268.512 | 849.025 | 0 | 0 | 0.213 | *P. teres f. teres* |
| Secreted | DisplGamma | -268.313 | 846.627 | 0 | 0 | 0.213 | *P. teres f. teres* |
| Secreted | ScaledBeta | -268.988 | 847.976 | 0.046 | 0.01 | 0.204 | *P. teres f. teres* |
| Secreted | Average | NA | NA | 0.006 | 0.001 | 0.212 | *P. teres f. teres* |
| Effectors | Neutral | -354.229 | 1014.457 | 0.979 | 0.248 | 0.005 | *P. teres f. teres* |
| Effectors | GammaZero | -139.759 | 587.519 | 0 | 0 | 0.254 | *P. teres f. teres* |
| Effectors | GammaGamma | -139.743 | 593.486 | 0.111 | 0.028 | 0.225 | *P. teres f. teres* |
| Effectors | GammaExpo | -139.759 | 591.518 | 0 | 0 | 0.254 | *P. teres f. teres* |
| Effectors | DisplGamma | -139.712 | 589.424 | 0 | 0 | 0.254 | *P. teres f. teres* |
| Effectors | ScaledBeta | -139.884 | 589.767 | 0.225 | 0.057 | 0.197 | *P. teres f. teres* |
| Effectors | Average | NA | NA | 0.041 | 0.011 | 0.243 | *P. teres f. teres* |
| Non-secreted | Neutral | -5960.536 | 12165.072 | 0.615 | 0.096 | 0.06 | *R. commune* |
| Non-secreted | GammaZero | -845.606 | 1937.212 | 0 | 0 | 0.156 | *R. commune* |
| Non-secreted | GammaGamma | -859.081 | 1970.163 | 0 | 0 | 0.156 | *R. commune* |
| Non-secreted | GammaExpo | -845.609 | 1941.217 | 0 | 0 | 0.156 | *R. commune* |
| Non-secreted | DisplGamma | -790.616 | 1829.231 | 0 | 0 | 0.156 | *R. commune* |
| Non-secreted | ScaledBeta | -907.749 | 2063.499 | 0.208 | 0.032 | 0.123 | *R. commune* |
| Non-secreted | Average | NA | NA | 0 | 0 | 0.156 | *R. commune* |
| Secreted | Neutral | -569.349 | 1382.698 | 0.59 | 0.115 | 0.08 | *R. commune* |
| Secreted | GammaZero | -392.538 | 1031.077 | 0 | 0 | 0.195 | *R. commune* |
| Secreted | GammaGamma | -394.666 | 1041.332 | 0.013 | 0.002 | 0.192 | *R. commune* |
| Secreted | GammaExpo | -392.625 | 1035.25 | 0.047 | 0.009 | 0.186 | *R. commune* |
| Secreted | DisplGamma | -393.355 | 1034.709 | 0 | 0 | 0.195 | *R. commune* |
| Secreted | ScaledBeta | -398.713 | 1045.426 | 0.32 | 0.062 | 0.132 | *R. commune* |
| Secreted | Average | NA | NA | 0.005 | 0.001 | 0.194 | *R. commune* |
| Effectors | Neutral | -285.668 | 815.336 | 0.638 | 0.158 | 0.089 | *R. commune* |

| | | | | | | | |
|---|---|---|---|---|---|---|---|
| Effectors | GammaZero | -217.611 | 681.221 | 0 | 0 | 0.247 | *R. commune* |
| Effectors | GammaGamma | -210.251 | 672.502 | 0.555 | 0.137 | 0.11 | *R. commune* |
| Effectors | GammaExpo | -211.114 | 672.228 | 0.437 | 0.108 | 0.139 | *R. commune* |
| Effectors | DisplGamma | -219.31 | 686.62 | 0 | 0 | 0.247 | *R. commune* |
| Effectors | ScaledBeta | -212.519 | 673.038 | 0.378 | 0.093 | 0.154 | *R. commune* |
| Effectors | Average | NA | NA | 0.46 | 0.114 | 0.133 | *R. commune* |
| Non-secreted | Neutral | -759609.615 | 1519467.23 | 0.974 | 0.18 | 0.005 | *S. musiva* |
| Non-secreted | GammaZero | -39152.393 | 78554.786 | 0 | 0 | 0.184 | *S. musiva* |
| Non-secreted | GammaGamma | -1095.633 | 2447.266 | 0.707 | 0.13 | 0.054 | *S. musiva* |
| Non-secreted | GammaExpo | -1101.992 | 2457.985 | 0.656 | 0.121 | 0.063 | *S. musiva* |
| Non-secreted | DisplGamma | -39152.393 | 78556.786 | 0 | 0 | 0.184 | *S. musiva* |
| Non-secreted | ScaledBeta | -1739.827 | 3731.654 | 0.809 | 0.149 | 0.035 | *S. musiva* |
| Non-secreted | Average | NA | NA | 0.707 | 0.13 | 0.054 | *S. musiva* |
| Secreted | Neutral | -23250.841 | 46749.683 | 0.968 | 0.213 | 0.007 | *S. musiva* |
| Secreted | GammaZero | -2419.07 | 5088.14 | 0 | 0 | 0.22 | *S. musiva* |
| Secreted | GammaGamma | -588.317 | 1432.633 | 0.914 | 0.201 | 0.019 | *S. musiva* |
| Secreted | GammaExpo | -587.488 | 1428.976 | 0.702 | 0.154 | 0.066 | *S. musiva* |
| Secreted | DisplGamma | -2419.071 | 5090.141 | 0 | 0 | 0.22 | *S. musiva* |
| Secreted | ScaledBeta | -610.508 | 1473.016 | 0.657 | 0.144 | 0.076 | *S. musiva* |
| Secreted | Average | NA | NA | 0.731 | 0.161 | 0.059 | *S. musiva* |
| Effectors | Neutral | -5416.986 | 11081.971 | 0.955 | 0.367 | 0.017 | *S. musiva* |
| Effectors | GammaZero | -1307.011 | 2864.021 | 0 | 0 | 0.385 | *S. musiva* |
| Effectors | GammaGamma | -450.259 | 1156.518 | 0.294 | 0.113 | 0.271 | *S. musiva* |
| Effectors | GammaExpo | -448.955 | 1151.911 | 0.595 | 0.229 | 0.156 | *S. musiva* |
| Effectors | DisplGamma | -1268.265 | 2788.53 | 0 | 0 | 0.385 | *S. musiva* |
| Effectors | ScaledBeta | -452.004 | 1156.008 | 0.705 | 0.271 | 0.114 | *S. musiva* |
| Effectors | Average | NA | NA | 0.582 | 0.224 | 0.161 | *S. musiva* |
| Non-secreted | Neutral | -274282.751 | 548617.502 | 0.941 | 0.166 | 0.01 | *S. sclerotiorum* |
| Non-secreted | GammaZero | -4728.242 | 9510.483 | 0 | 0 | 0.176 | *S. sclerotiorum* |
| Non-secreted | GammaGamma | -541.788 | 1143.576 | 0 | 0 | 0.176 | *S. sclerotiorum* |
| Non-secreted | GammaExpo | -461.376 | 980.751 | 0.002 | 0 | 0.176 | *S. sclerotiorum* |
| Non-secreted | DisplGamma | -4728.242 | 9512.483 | 0 | 0 | 0.176 | *S. sclerotiorum* |
| Non-secreted | ScaledBeta | -572.7 | 1201.401 | 0.181 | 0.032 | 0.144 | *S. sclerotiorum* |
| Non-secreted | Average | NA | NA | 0.002 | 0 | 0.176 | *S. sclerotiorum* |
| Secreted | Neutral | -10177.929 | 20407.859 | 0.932 | 0.183 | 0.013 | *S. sclerotiorum* |
| Secreted | GammaZero | -383.944 | 821.888 | 0 | 0 | 0.196 | *S. sclerotiorum* |
| Secreted | GammaGamma | -141.702 | 343.403 | 0 | 0 | 0.196 | *S. sclerotiorum* |
| Secreted | GammaExpo | -143.09 | 344.18 | 0.295 | 0.058 | 0.138 | *S. sclerotiorum* |
| Secreted | DisplGamma | -383.944 | 823.888 | 0 | 0 | 0.196 | *S. sclerotiorum* |
| Secreted | ScaledBeta | -136.987 | 329.973 | 0.016 | 0.003 | 0.193 | *S. sclerotiorum* |
| Secreted | Average | NA | NA | 0.017 | 0.003 | 0.193 | *S. sclerotiorum* |
| Effectors | Neutral | -3525.59 | 7103.179 | 0.924 | 0.214 | 0.017 | *S. sclerotiorum* |
| Effectors | GammaZero | -221.96 | 497.92 | 0 | 0 | 0.231 | *S. sclerotiorum* |
| Effectors | GammaGamma | -111.324 | 282.648 | 0 | 0 | 0.231 | *S. sclerotiorum* |
| Effectors | GammaExpo | -112.831 | 283.662 | 0.549 | 0.127 | 0.104 | *S. sclerotiorum* |
| Effectors | DisplGamma | -211.124 | 478.248 | 0 | 0 | 0.231 | *S. sclerotiorum* |
| Effectors | ScaledBeta | -111.912 | 279.825 | 0.439 | 0.101 | 0.13 | *S. sclerotiorum* |
| Effectors | Average | NA | NA | 0.374 | 0.086 | 0.145 | *S. sclerotiorum* |
| Non-secreted | Neutral | -5097.404 | 10408.808 | 0.929 | 0.141 | 0.011 | *V. dahliae* |
| Non-secreted | GammaZero | -344.662 | 905.324 | 0 | 0 | 0.151 | *V. dahliae* |
| Non-secreted | GammaGamma | -353.33 | 928.66 | 0 | 0 | 0.151 | *V. dahliae* |
| Non-secreted | GammaExpo | -344.837 | 909.674 | 0 | 0 | 0.151 | *V. dahliae* |
| Non-secreted | DisplGamma | -344.272 | 906.545 | 0 | 0 | 0.151 | *V. dahliae* |
| Non-secreted | ScaledBeta | -360.136 | 938.272 | 0.019 | 0.003 | 0.149 | *V. dahliae* |
| Non-secreted | Average | NA | NA | 0 | 0 | 0.151 | *V. dahliae* |
| Secreted | Neutral | -380.869 | 975.737 | 0.901 | 0.145 | 0.016 | *V. dahliae* |
| Secreted | GammaZero | -163.142 | 542.285 | 0 | 0 | 0.161 | *V. dahliae* |
| Secreted | GammaGamma | -163.36 | 548.719 | 0 | 0 | 0.161 | *V. dahliae* |
| Secreted | GammaExpo | -163.133 | 546.266 | 0 | 0 | 0.161 | *V. dahliae* |
| Secreted | DisplGamma | -162.281 | 542.562 | 0 | 0 | 0.161 | *V. dahliae* |

| | | | | | | | |
|---|---|---|---|---|---|---|---|
| Secreted | ScaledBeta | -166.505 | 551.009 | 0.098 | 0.016 | 0.145 | *V. dahliae* |
| Secreted | Average | NA | NA | 0.001 | 0 | 0.161 | *V. dahliae* |
| Effectors | Neutral | -88.031 | 390.062 | 0.893 | 0.175 | 0.021 | *V. dahliae* |
| Effectors | GammaZero | -57.451 | 330.901 | 0 | 0 | 0.196 | *V. dahliae* |
| Effectors | GammaGamma | -57.451 | 336.902 | 0 | 0 | 0.196 | *V. dahliae* |
| Effectors | GammaExpo | -57.451 | 334.902 | 0 | 0 | 0.196 | *V. dahliae* |
| Effectors | DisplGamma | -57.273 | 332.546 | 0 | 0 | 0.196 | *V. dahliae* |
| Effectors | ScaledBeta | -58.27 | 334.54 | 0.112 | 0.022 | 0.174 | *V. dahliae* |
| Effectors | Average | NA | NA | 0.01 | 0.002 | 0.194 | *V. dahliae* |
| Non-secreted | Neutral | -299065.33 | 598300.66 | 0.954 | 0.163 | 0.008 | *V. inaequalis* |
| Non-secreted | GammaZero | -49794.921 | 99761.842 | 0 | 0 | 0.171 | *V. inaequalis* |
| Non-secreted | GammaGamma | -1299.559 | 2777.119 | 0.758 | 0.129 | 0.041 | *V. inaequalis* |
| Non-secreted | GammaExpo | -1506.905 | 3189.81 | 0.693 | 0.118 | 0.052 | *V. inaequalis* |
| Non-secreted | DisplGamma | -49794.921 | 99763.842 | 0 | 0 | 0.171 | *V. inaequalis* |
| Non-secreted | ScaledBeta | -2553.17 | 5280.34 | 0.945 | 0.161 | 0.009 | *V. inaequalis* |
| Non-secreted | Average | NA | NA | 0.758 | 0.129 | 0.041 | *V. inaequalis* |
| Secreted | Neutral | -4078.865 | 8327.731 | 0.932 | 0.207 | 0.015 | *V. inaequalis* |
| Secreted | GammaZero | -1190.941 | 2553.883 | 0 | 0 | 0.222 | *V. inaequalis* |
| Secreted | GammaGamma | -326.047 | 830.093 | 0.738 | 0.164 | 0.058 | *V. inaequalis* |
| Secreted | GammaExpo | -331.446 | 838.892 | 0.659 | 0.146 | 0.076 | *V. inaequalis* |
| Secreted | DisplGamma | -1184.114 | 2542.227 | 0 | 0 | 0.222 | *V. inaequalis* |
| Secreted | ScaledBeta | -345.496 | 864.991 | 0.94 | 0.208 | 0.013 | *V. inaequalis* |
| Secreted | Average | NA | NA | 0.737 | 0.163 | 0.058 | *V. inaequalis* |
| Effectors | Neutral | -1340.791 | 2851.582 | 0.938 | 0.247 | 0.016 | *V. inaequalis* |
| Effectors | GammaZero | -538.069 | 1248.139 | 0 | 0 | 0.264 | *V. inaequalis* |
| Effectors | GammaGamma | -186.838 | 551.676 | 0.898 | 0.237 | 0.027 | *V. inaequalis* |
| Effectors | GammaExpo | -188.611 | 553.221 | 0.912 | 0.241 | 0.023 | *V. inaequalis* |
| Effectors | DisplGamma | -529.867 | 1233.734 | 0 | 0 | 0.264 | *V. inaequalis* |
| Effectors | ScaledBeta | -192.621 | 559.243 | 0.954 | 0.252 | 0.012 | *V. inaequalis* |
| Effectors | Average | NA | NA | 0.904 | 0.238 | 0.025 | *V. inaequalis* |
| Non-secreted | Neutral | -100170.173 | 201314.345 | 0.909 | 0.11 | 0.011 | *Z. tritici* |
| Non-secreted | GammaZero | -5629.966 | 12235.932 | 0 | 0 | 0.121 | *Z. tritici* |
| Non-secreted | GammaGamma | -3211.584 | 7405.168 | 0.502 | 0.06 | 0.06 | *Z. tritici* |
| Non-secreted | GammaExpo | -3254.89 | 7489.78 | 0.471 | 0.057 | 0.064 | *Z. tritici* |
| Non-secreted | DisplGamma | -5797.721 | 12573.441 | 0 | 0 | 0.121 | *Z. tritici* |
| Non-secreted | ScaledBeta | -4360.86 | 9699.72 | 0.711 | 0.086 | 0.035 | *Z. tritici* |
| Non-secreted | Average | NA | NA | 0.502 | 0.06 | 0.06 | *Z. tritici* |
| Secreted | Neutral | -4438.532 | 9851.063 | 0.9 | 0.131 | 0.015 | *Z. tritici* |
| Secreted | GammaZero | -1426.699 | 3829.399 | 0 | 0 | 0.145 | *Z. tritici* |
| Secreted | GammaGamma | -1317.824 | 3617.649 | 0.661 | 0.096 | 0.049 | *Z. tritici* |
| Secreted | GammaExpo | -1318.333 | 3616.667 | 0.651 | 0.094 | 0.051 | *Z. tritici* |
| Secreted | DisplGamma | -1426.879 | 3831.758 | 0 | 0 | 0.145 | *Z. tritici* |
| Secreted | ScaledBeta | -1379.754 | 3737.507 | 0.828 | 0.12 | 0.025 | *Z. tritici* |
| Secreted | Average | NA | NA | 0.655 | 0.095 | 0.05 | *Z. tritici* |
| Effectors | Neutral | -1042.869 | 3059.738 | 0.892 | 0.172 | 0.021 | *Z. tritici* |
| Effectors | GammaZero | -516.386 | 2008.771 | 0 | 0 | 0.193 | *Z. tritici* |
| Effectors | GammaGamma | -494.686 | 1971.372 | 0.65 | 0.125 | 0.068 | *Z. tritici* |
| Effectors | GammaExpo | -494.693 | 1969.387 | 0.635 | 0.122 | 0.07 | *Z. tritici* |
| Effectors | DisplGamma | -521.743 | 2021.486 | 0 | 0 | 0.193 | *Z. tritici* |
| Effectors | ScaledBeta | -497.803 | 1973.607 | 0.649 | 0.125 | 0.068 | *Z. tritici* |
| Effectors | Average | NA | NA | 0.64 | 0.123 | 0.069 | *Z. tritici* |

**Table S12.** Number of accessory gene and categories per species.

| species | effector_protein | nonsecreted | secreted_protein | total_accessory |
|---|---|---|---|---|
| *A. flavus* | 30 | 654 | 43 | 727 |
| *B. cinera* | 167 | 6116 | 287 | 6570 |
| *C. beticola* | 27 | 585 | 34 | 646 |
| *C. parasitica* | 4 | 533 | 3 | 540 |
| *F. graminearum* | 39 | 764 | 44 | 847 |
| *M. oryzae rice* | 128 | 860 | 91 | 1079 |
| *M. oryzae Triticum* | 40 | 510 | 18 | 568 |
| *N. discreta* | 17 | 819 | 22 | 858 |
| *O. novo-ulmi subsp. americana* | 0 | 63 | 4 | 67 |
| *O. novo-ulmi subsp. novo-ulmi* | 0 | 92 | 3 | 95 |
| *O. ulmi* | 0 | 122 | 5 | 127 |
| *P. biforme* | 51 | 1510 | 64 | 1625 |
| *P. nodorum* | 40 | 1034 | 62 | 1136 |
| *P. teres f. teres* | 14 | 477 | 6 | 497 |
| *R. commune* | 58 | 792 | 57 | 907 |
| *S. musiva* | 11 | 307 | 7 | 325 |
| *S. sclerotiorum* | 43 | 2638 | 31 | 2712 |
| *V. dahliae* | 23 | 1436 | 76 | 1535 |
| *V. inaequalis* | 213 | 1240 | 77 | 1530 |
| *Z. tritici* | 145 | 2947 | 129 | 3221 |

**Table S13.** Number of protein-coding genes and number of isolates used for the accessory and core gene datasets. The same isolates were used for both gene datasets, but different genes.

| **Species** | **Number of genes used** | **Number of isolates** |
|---|---|---|
| *Aspergilus flavus* | 47 | 52 |
| *Botrytis cinerea* | 3524 | 47 |
| *Cercospora beticola* | 111 | 90 |
| *Cryphonectria parasitica* | 8 | 220 |
| *Fusarium graminearum* | 89 | 19 |
| *Magnaporthe oryzae* rice | 136 | 27 |
| *Magnaporthe oryzae triticum** | 15 | 9 |
| *Neurospora discreta* | 128 | 22 |
| *Ophiostoma novo-ulmi subsp. Americana** | 2 | 25 |
| *Ophiostoma novo-ulmi subsp. novo-ulmi** | 6 | 26 |
| *Ophiostoma ulmi** | 6 | 35 |
| *Parastagonospora nodorum* | 58 | 226 |
| *Penicillium biforme** | 14 | 16 |
| *Pyrenophora teres f. sp. teres* | 17 | 124 |
| *Rhynchosporium commune* | 76 | 56 |
| *Sclerotinia sclerotiorum* | 148 | 13 |
| *Sphaerulina musiva* | 22 | 93 |
| *Venturia inaequalis* | 164 | 18 |
| *Verticillium dahliae* | 108 | 60 |
| *Zymoseptoria tritici* | 503 | 36 |

*Grapes could not complete using these datasets.

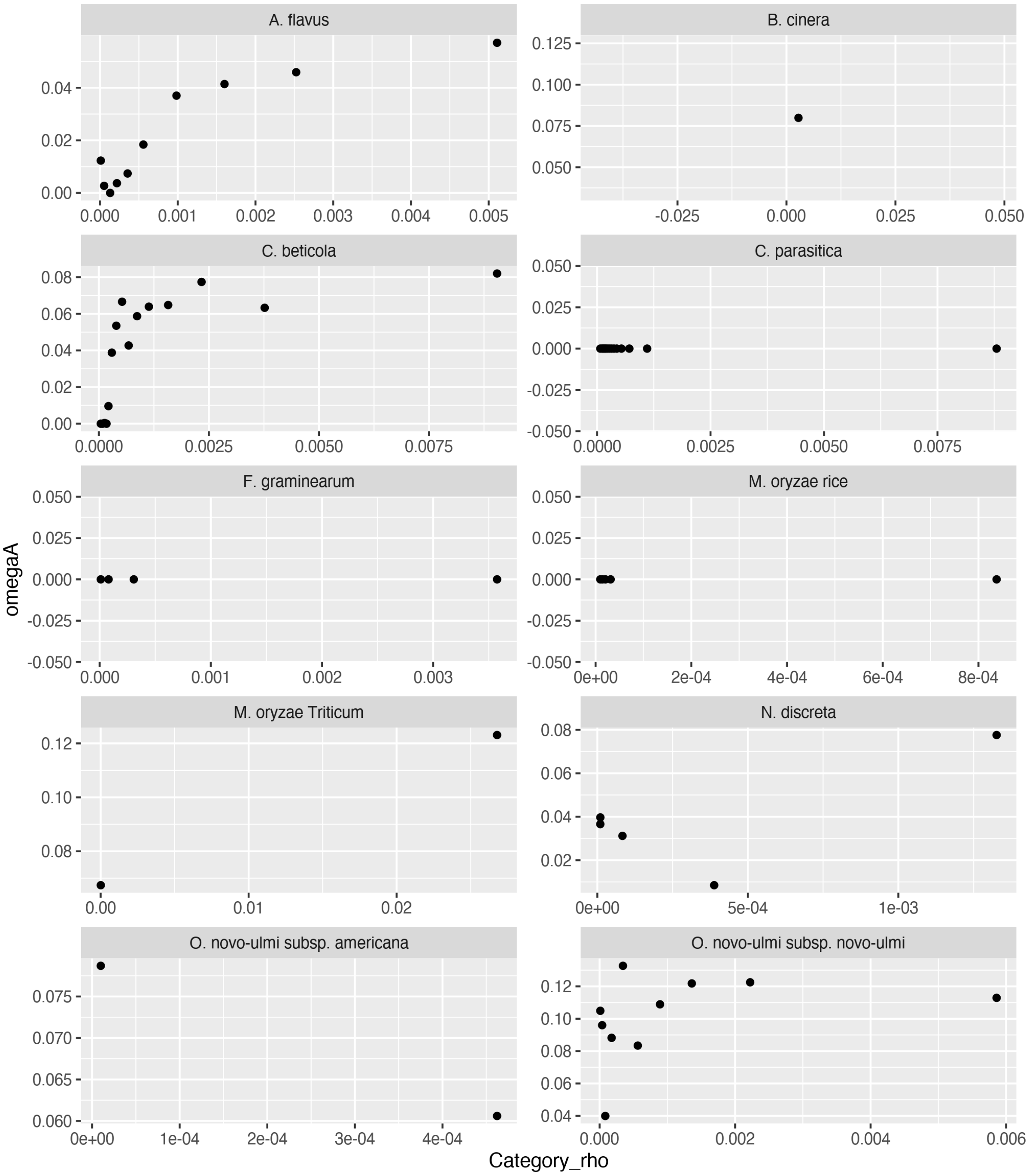


**Supplementary figure 1.** Correlation between omegaA and rho across the genome. Each dot represents a set of genes with similar levels of recombination.

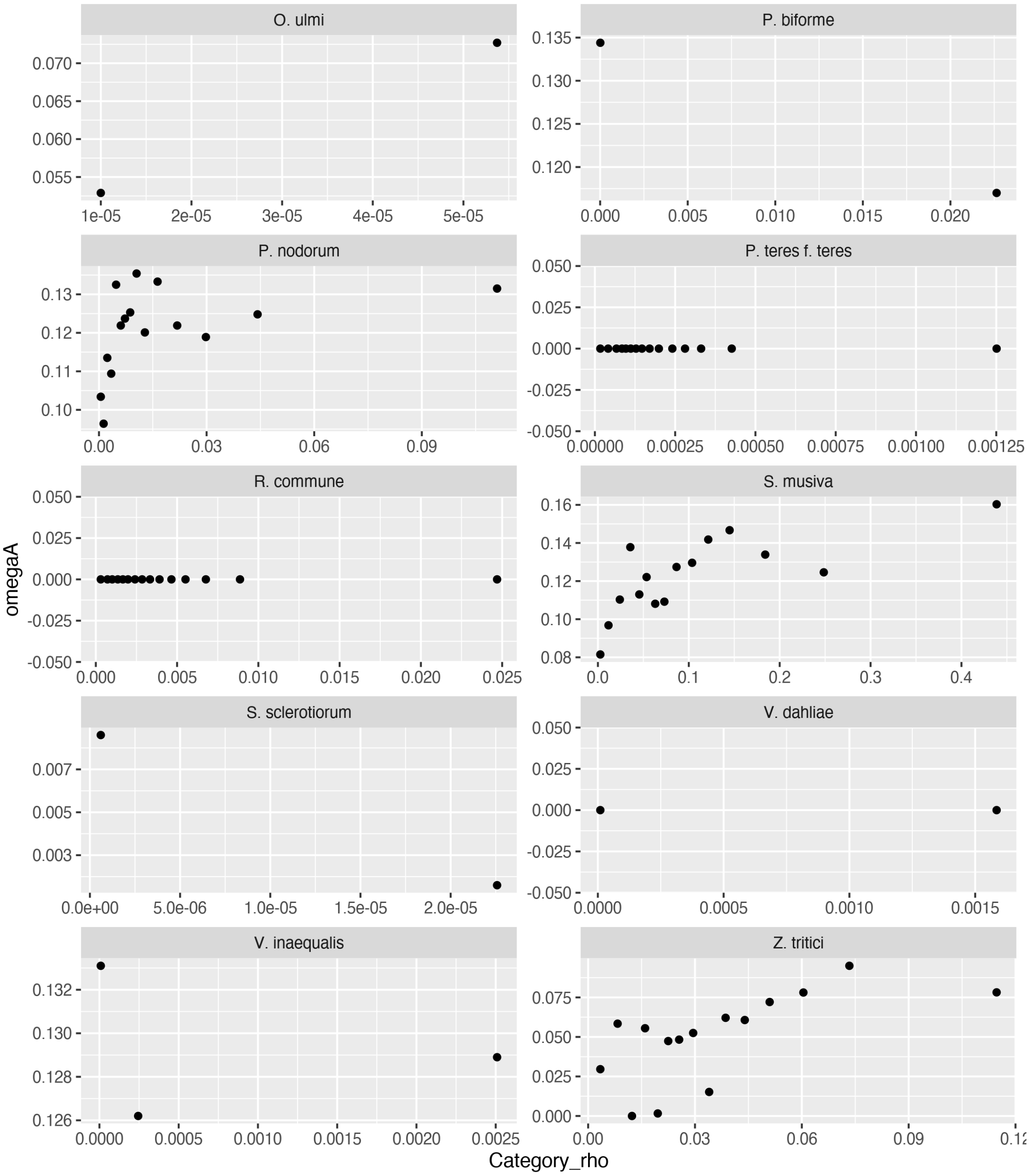


**Supplementary figure 1 (continuation).** Correlation between omegaA and rho across the genome. Each dot represents a set of genes with similar levels of recombination.

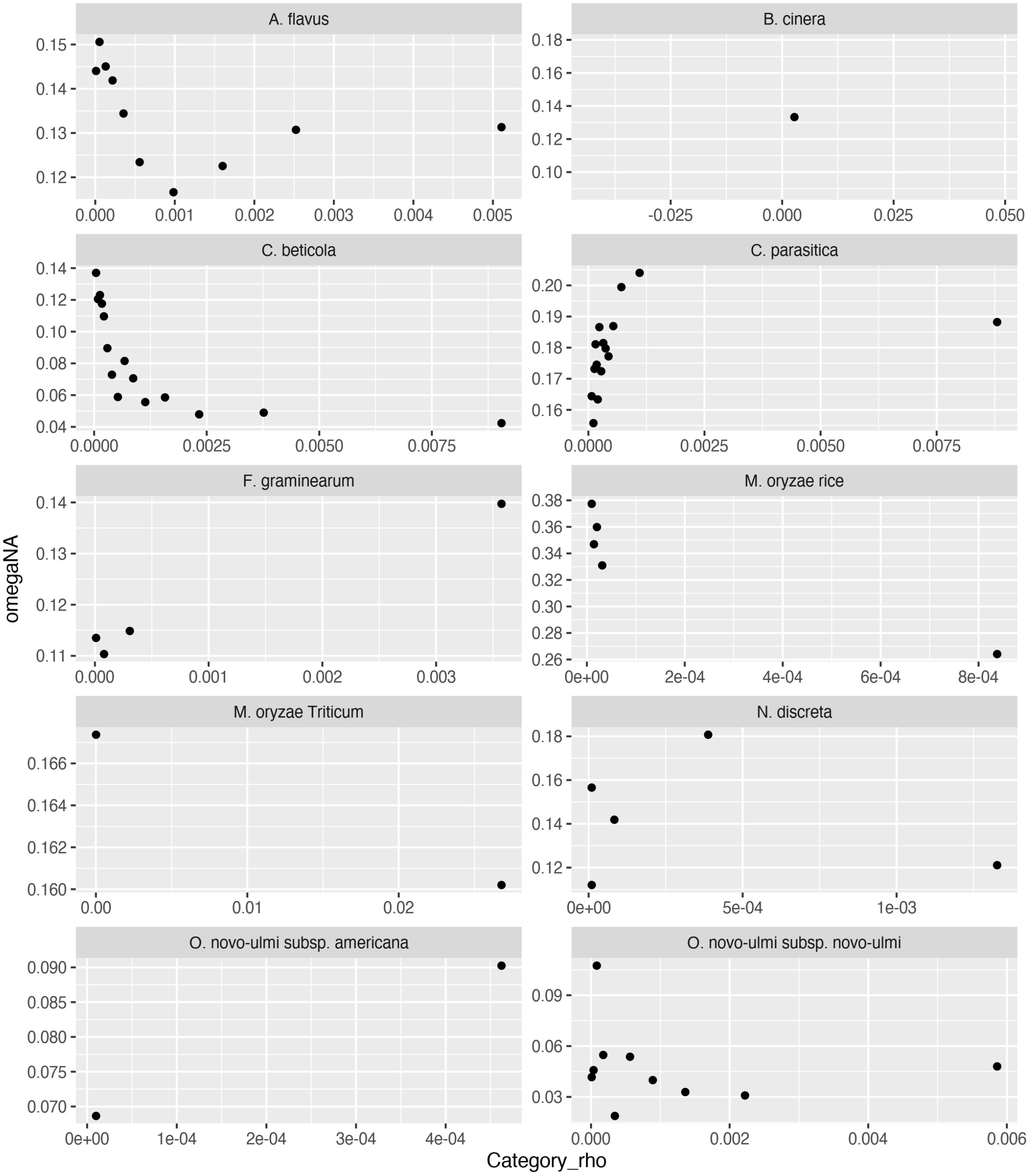


**Supplementary figure 2.** Correlation between omegaNA and rho across the genome. Each dot represents a set of genes with similar levels of recombination.

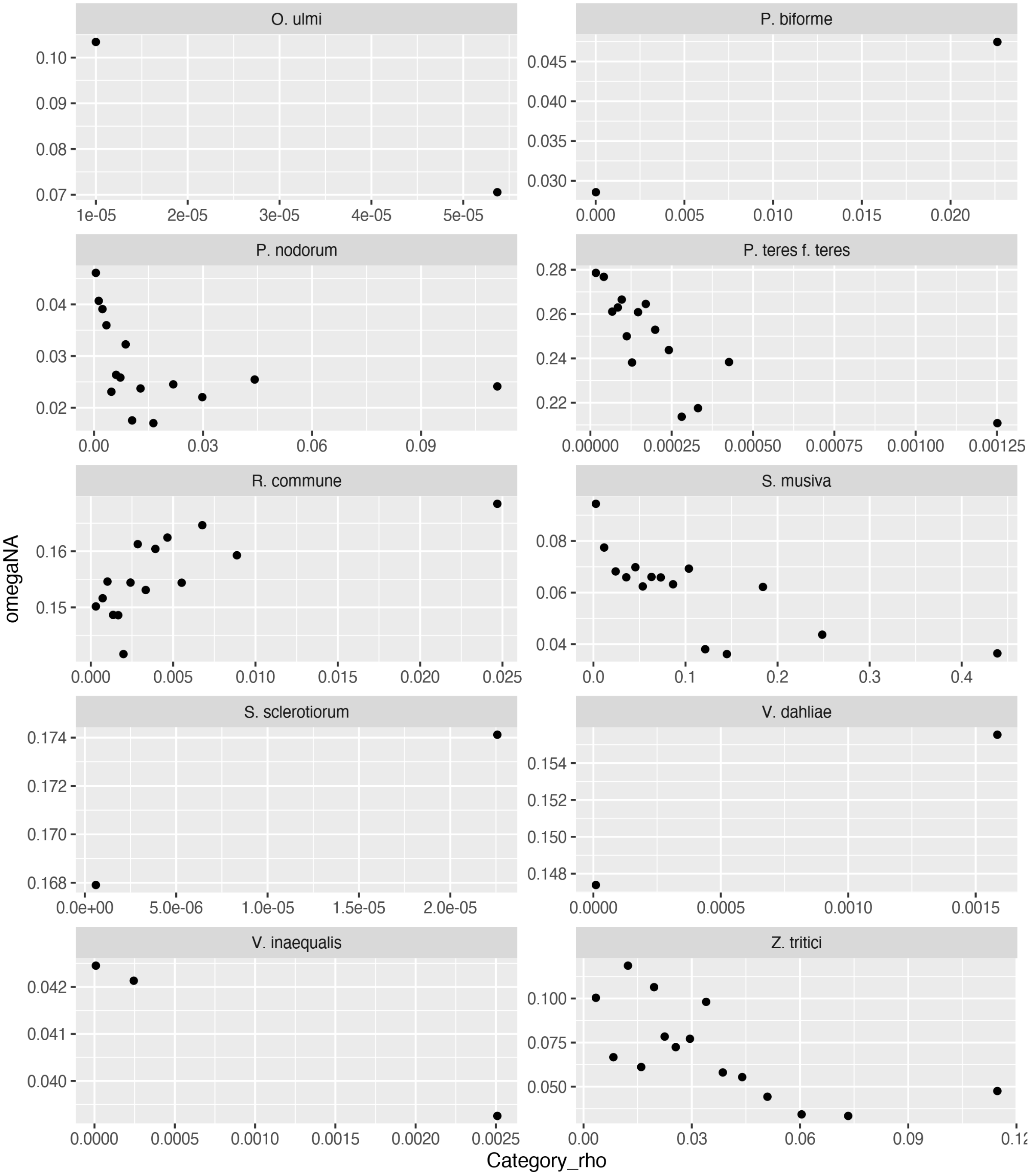


**Supplementary figure 2 (continuation).** Correlation between omegaNA and rho across the genome. Each dot represents a set of genes with similar levels of recombination.

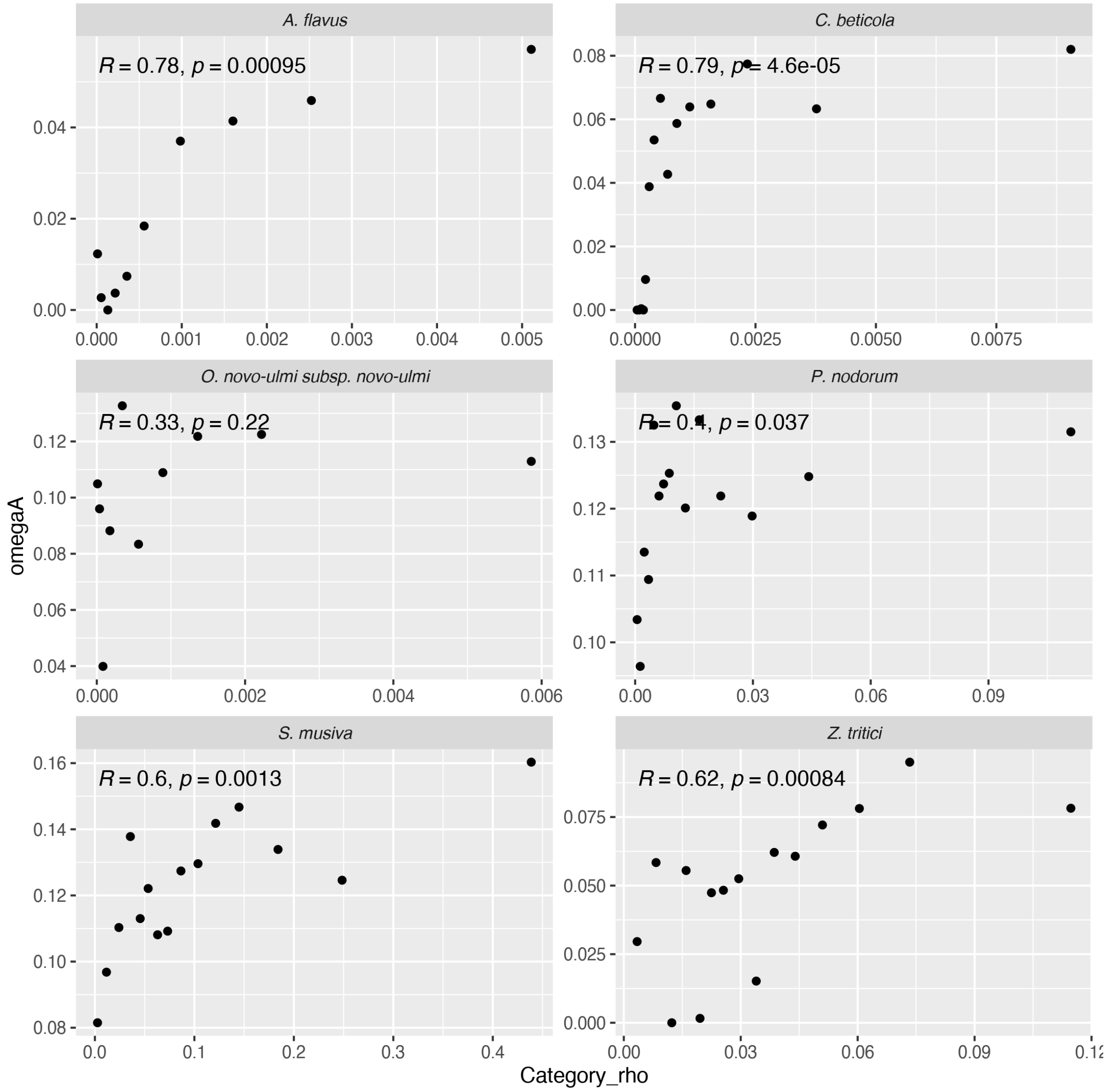


**Supplementary figure 3.** Correlation between omegaA and rho across the genome. Each dot represents a set of genes with similar levels of recombination. Kendall's correlation used.

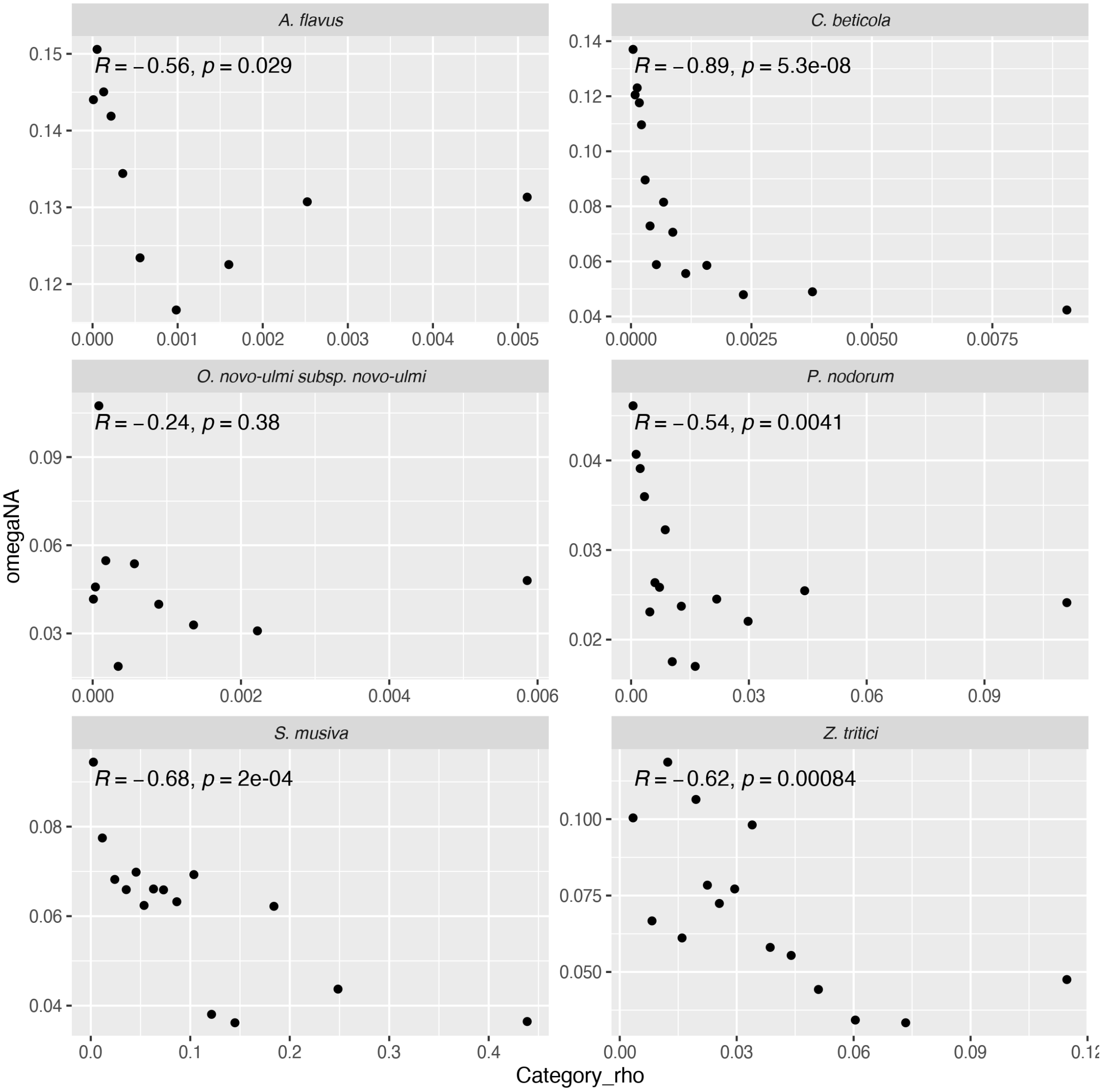


**Supplementary figure 4.** Correlation between omegaNA and rho across the genome. Each dot represents a set of genes with similar levels of recombination. Kendall's correlation used.

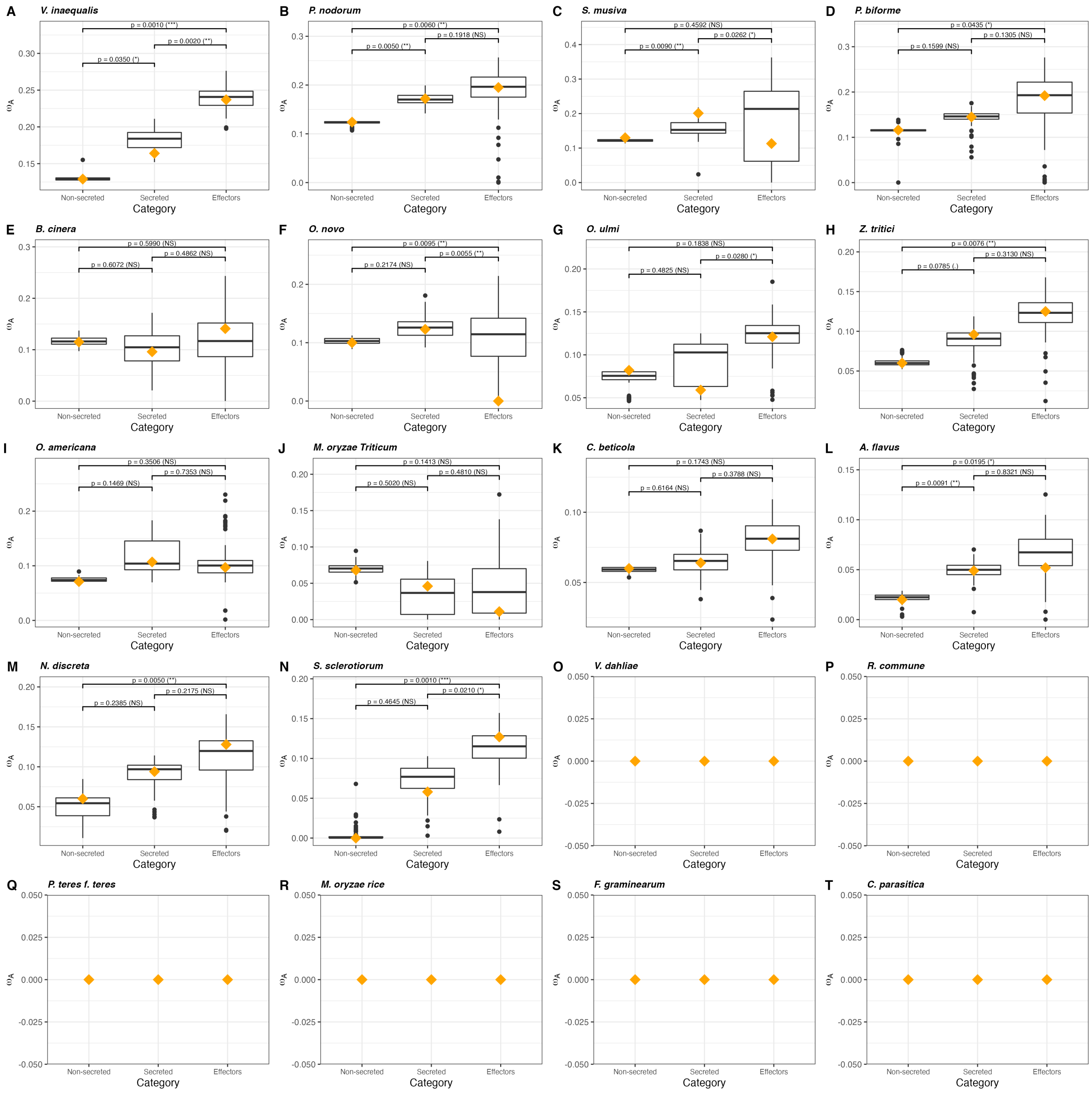


**Supplementary figure 5.** Landscape of adaptive nonsynonymous substitutions rate ($\omega_A$) among non-secreted, secreted and effector gene categories within 20 fungal species. (A) *Venturia inaequalis,* (B) *Parastagonospora nodorum,* (C) *Sphaerulina musiva,* (D) *Penicillium biforme,* (E) *Botrytis cinerea,* (F) *Ophiostoma novo-ulmi subsp. novo-ulmi,* (G) *Ophiostoma ulmi,* (H) *Zymoseptoria tritici,* (I) *Ophiostoma novo-ulmi subsp. americana,* (J) *Magnaporthe oryzae Triticum,* (K) *Cercospora beticola,* (L) *Aspergillus flavus,* (M) *Neurospora discreta,* (N) *Sclerotinia sclerotiorum,* (O) *Verticillium dahliae,* (P) *Rhynchosporium commune,* (Q) *Pyrenophora teres f. sp. teres,* (R) *Magnaporthe oryzae rice,* (S) *Fusarium graminearum,* (T) *Cryphonectria parasitica*. Box-plot distribution was generated using 100 bootstrap replicates performed using the model with the lowest Akaike's information criterion. The orange diamond represents the mean value calculated after averaging over all models of distribution of fitness effects.

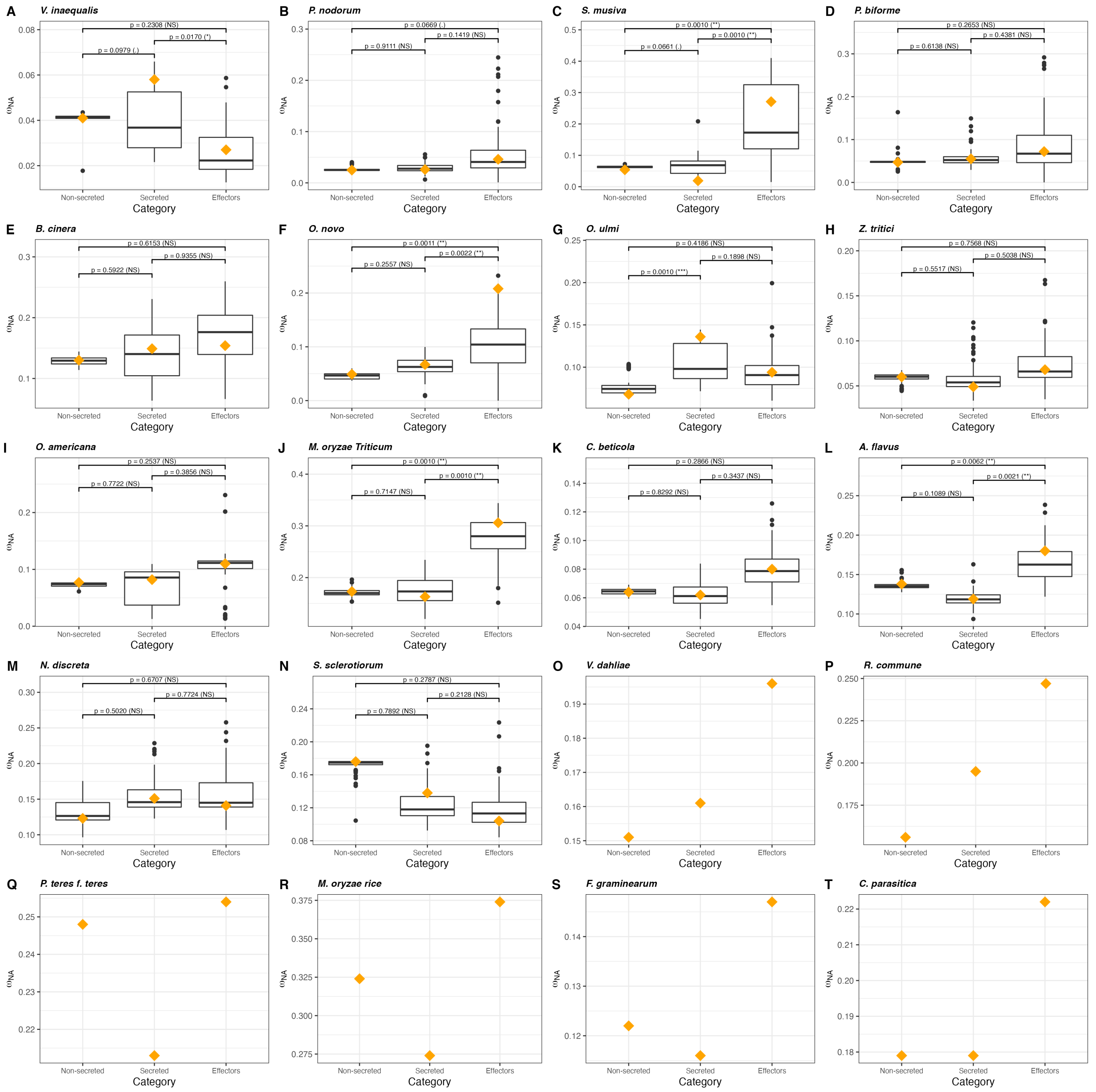

**Supplementary figure 6.** Landscape of nonadaptive nonsynonymous substitutions rate ($\omega_{NA}$) among non-secreted, secreted and effector gene categories within 20 fungal species. (A) *Venturia inaequalis,* (B) *Parastagonospora nodorum,* (C) *Sphaerulina musiva,* (D) *Penicillium biforme,* (E) *Botrytis cinerea,* (F) *Ophiostoma novo-ulmi subsp. novo-ulmi,* (G) *Ophiostoma ulmi,* (H) *Zymoseptoria tritici,* (I) *Ophiostoma novo-ulmi subsp. americana,* (J) *Magnaporthe oryzae Triticum,* (K) *Cercospora beticola,* (L) *Aspergillus flavus,* (M) *Neurospora discreta,* (N) *Sclerotinia sclerotiorum,* (O) *Verticillium dahliae,* (P) *Rhynchosporium commune,* (Q) *Pyrenophora teres f. sp. teres,* (R) *Magnaporthe oryzae rice,* (S) *Fusarium graminearum,* (T) *Cryphonectria parasitica*. Box-plot distribution was generated using 100 bootstrap replicates performed using the model with the lowest Akaike's information criterion. The orange diamond represents the mean value calculated after averaging over all models of distribution of fitness effects.

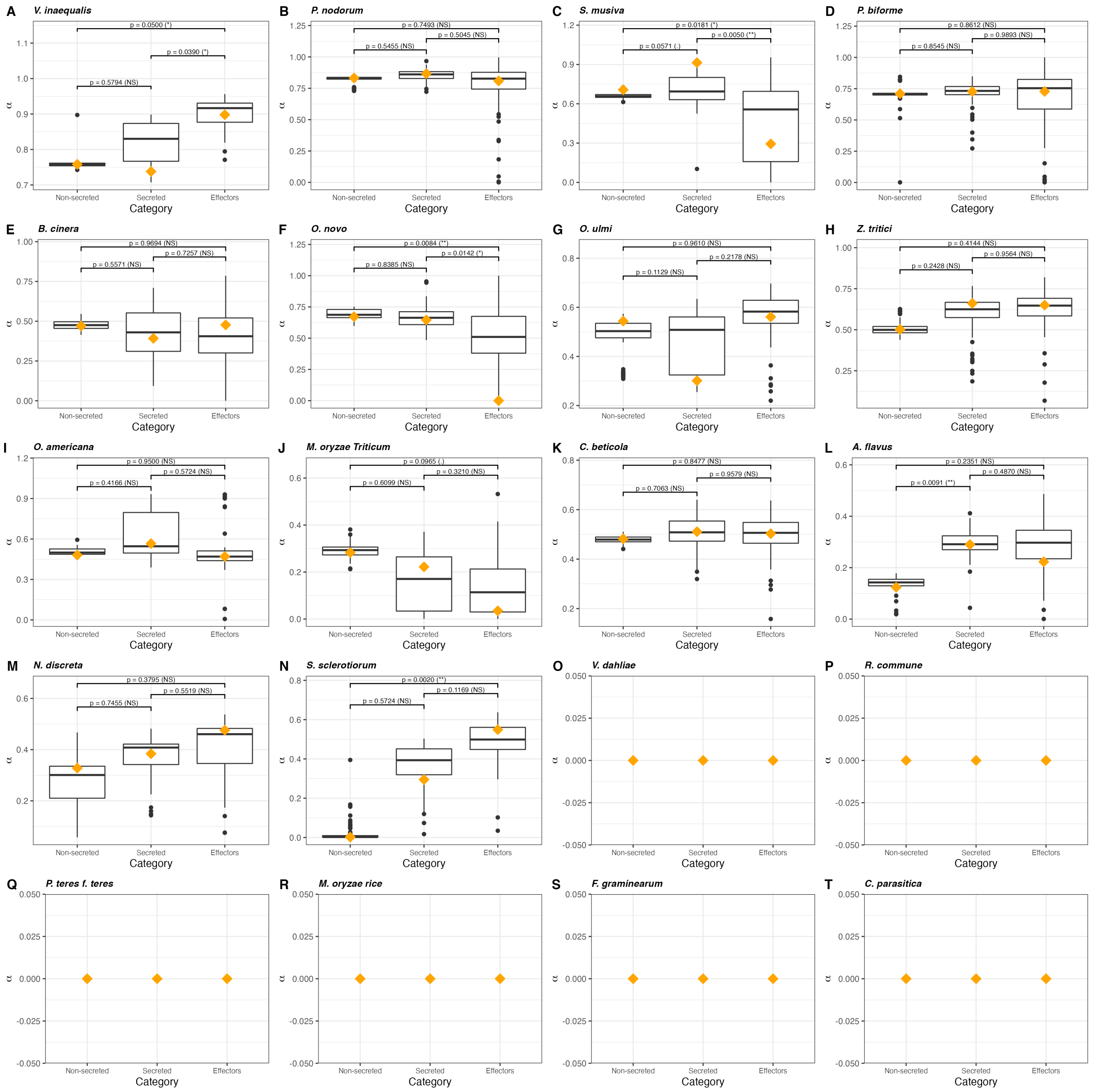


**Supplementary figure 7.** Proportion of amino-acid substitutions that are adaptive (α) among non-secreted, secreted and effector gene categories within 20 fungal species. (A) *Venturia inaequalis,* (B) *Parastagonospora nodorum,* (C) *Sphaerulina musiva,* (D) *Penicillium biforme,* (E) *Botrytis cinerea,* (F) *Ophiostoma novo-ulmi subsp. novo-ulmi,* (G) *Ophiostoma ulmi,* (H) *Zymoseptoria tritici,* (I) *Ophiostoma novo-ulmi subsp. americana,* (J) *Magnaporthe oryzae Triticum,* (K) *Cercospora beticola,* (L) *Aspergillus flavus,* (M) *Neurospora discreta,* (N) *Sclerotinia sclerotiorum,* (O) *Verticillium dahliae,* (P) *Rhynchosporium commune,* (Q) *Pyrenophora teres f. sp. teres,* (R) *Magnaporthe oryzae rice,* (S) *Fusarium graminearum,* (T) *Cryphonectria parasitica*. Box-plot distribution was generated using 100 bootstrap replicates performed using the model with the lowest Akaike's information criterion. The orange diamond represents the mean value calculated after averaging over all models of distribution of fitness effects.

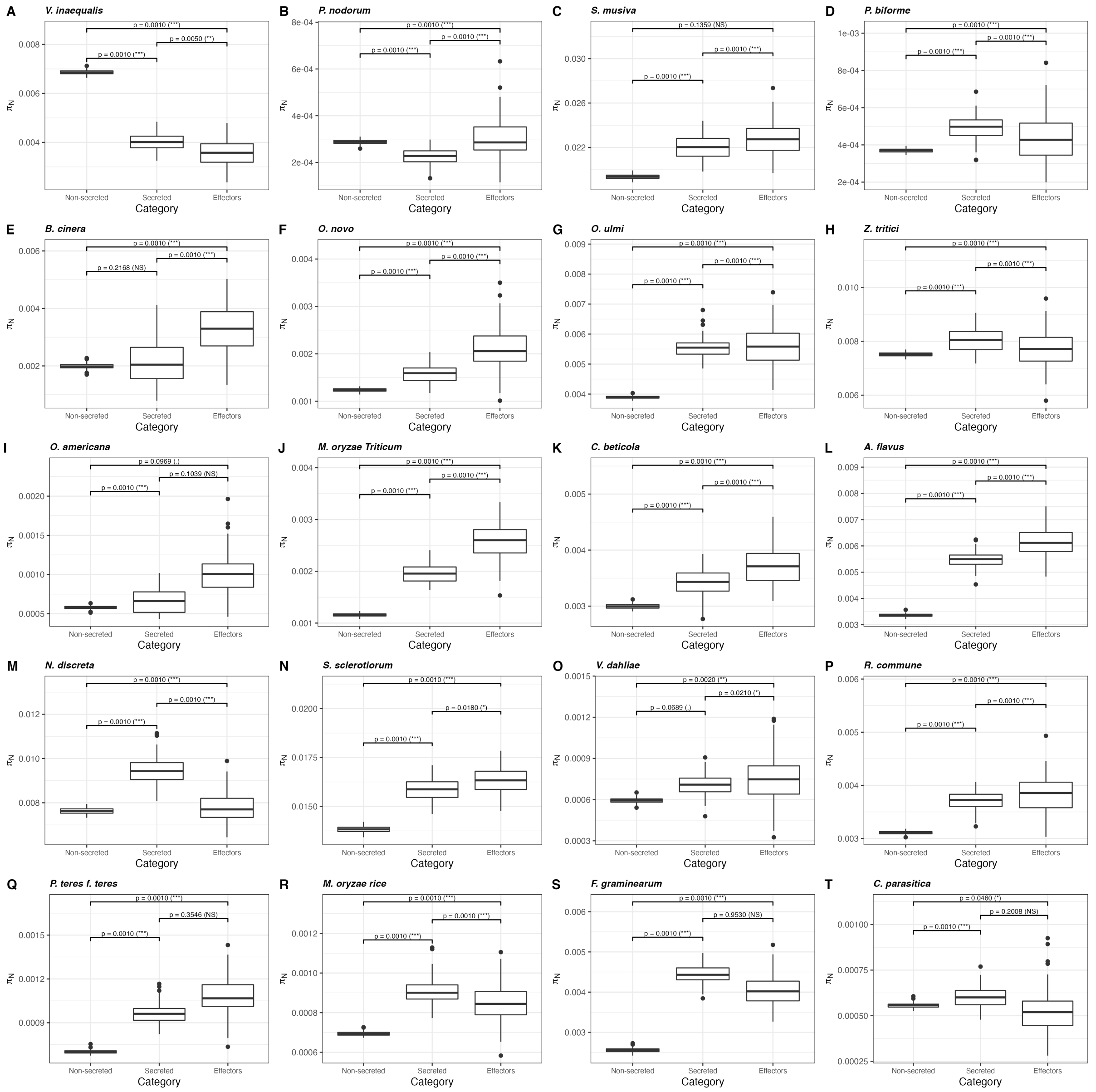


**Supplementary figure 8.** Variation in the nonsynonymous diversity ($\pi_N$) across 20 fungal species. (A) *Venturia inaequalis,* (B) *Parastagonospora nodorum,* (C) *Sphaerulina musiva,* (D) *Penicillium biforme,* (E) *Botrytis cinerea,* (F) *Ophiostoma novo-ulmi subsp. novo-ulmi,* (G) *Ophiostoma ulmi,* (H) *Zymoseptoria tritici,* (I) *Ophiostoma novo-ulmi subsp. americana,* (J) *Magnaporthe oryzae Triticum,* (K) *Cercospora beticola,* (L) *Aspergillus flavus,* (M) *Neurospora discreta,* (N) *Sclerotinia sclerotiorum,* (O) *Verticillium dahliae,* (P) *Rhynchosporium commune,* (Q) *Pyrenophora teres f. sp. teres,* (R) *Magnaporthe oryzae rice,* (S) *Fusarium graminearum,* (T) *Cryphonectria parasitica*. Box-plot distribution was generated using 100 bootstrap replicates performed using the model with the lowest Akaike's information criterion.

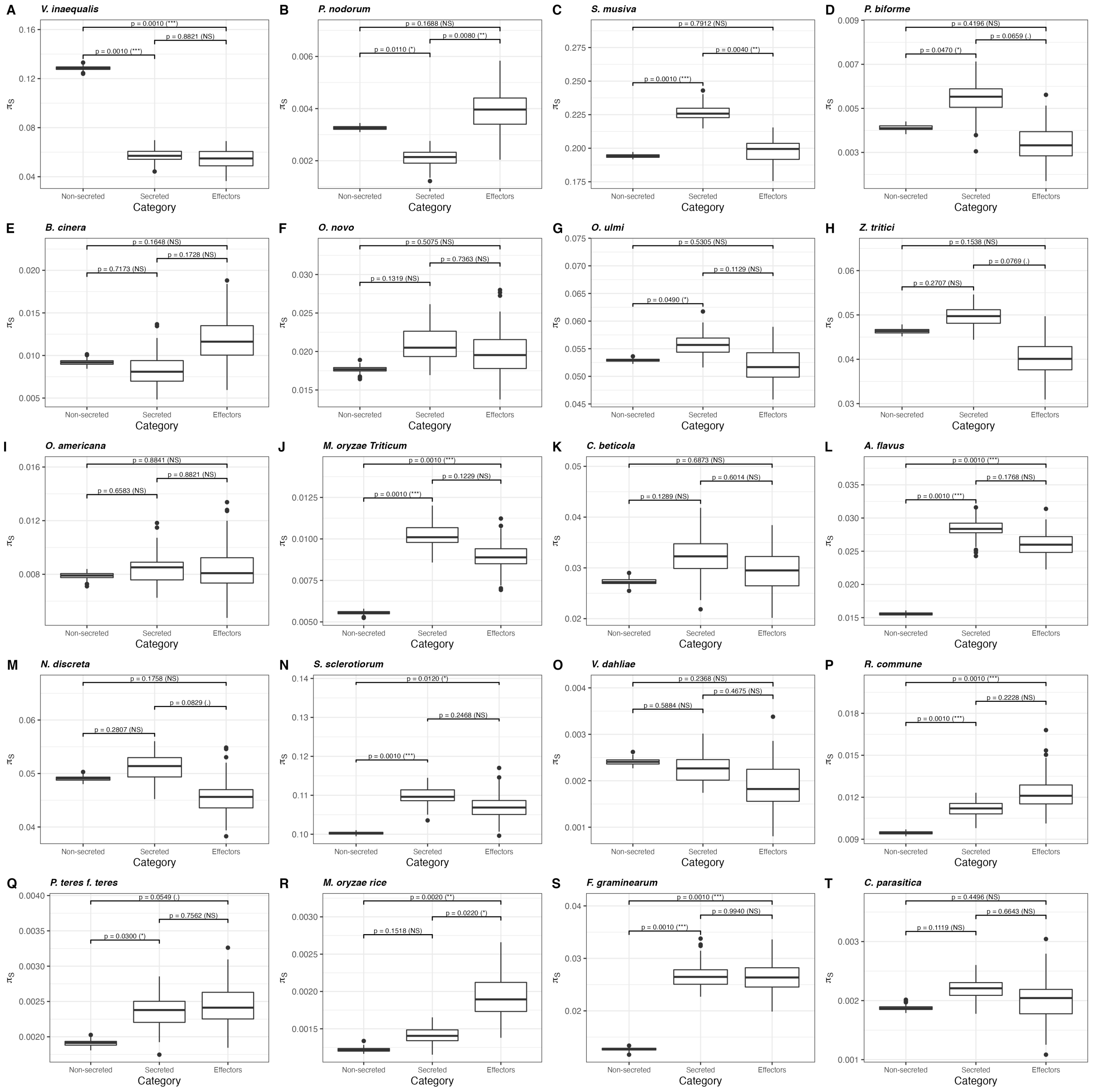


**Supplementary figure 9.** Variation in the synonymous diversity ($\pi_S$) across 20 fungal species. (A) *Venturia inaequalis,* (B) *Parastagonospora nodorum,* (C) *Sphaerulina musiva,* (D) *Penicillium biforme,* (E) *Botrytis cinerea,* (F) *Ophiostoma novo-ulmi subsp. novo-ulmi,* (G) *Ophiostoma ulmi,* (H) *Zymoseptoria tritici,* (I) *Ophiostoma novo-ulmi subsp. americana,* (J) *Magnaporthe oryzae Triticum,* (K) *Cercospora beticola,* (L) *Aspergillus flavus,* (M) *Neurospora discreta,* (N) *Sclerotinia sclerotiorum,* (O) *Verticillium dahliae,* (P) *Rhynchosporium commune,* (Q) *Pyrenophora teres f. sp. teres,* (R) *Magnaporthe oryzae rice,* (S) *Fusarium graminearum,* (T) *Cryphonectria parasitica*. Box-plot distribution was generated using 100 bootstrap replicates performed using the model with the lowest Akaike's information criterion.

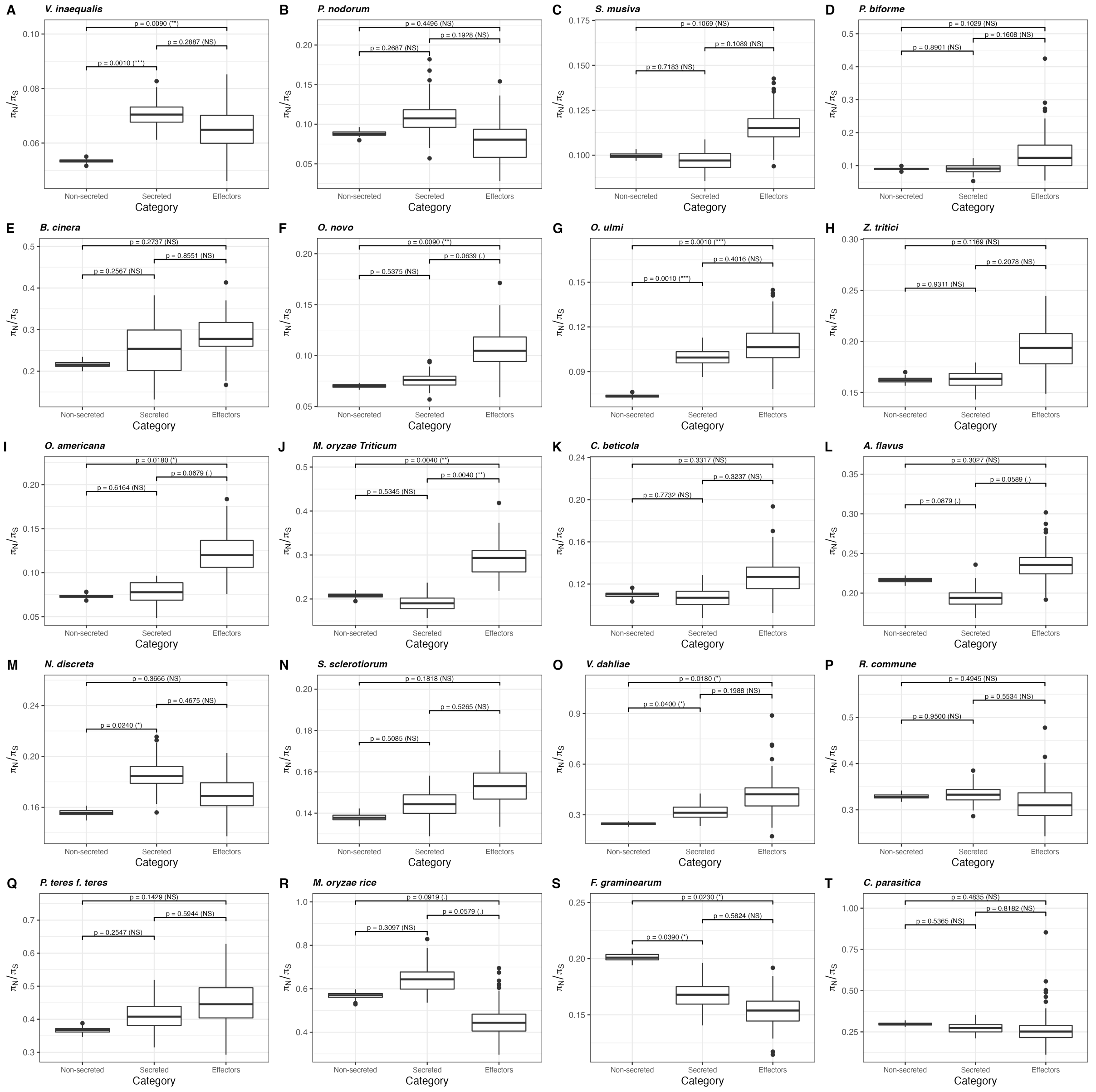


**Supplementary figure 10.** Variation in the ratio of the mean number of nonsynonymous nucleotide differences and mean number of synonymous nucleotide differences ($\pi_N/\pi_S$) across 20 fungal species. (A) *Venturia inaequalis,* (B) *Parastagonospora nodorum,* (C) *Sphaerulina musiva,* (D) *Penicillium biforme,* (E) *Botrytis cinerea,* (F) *Ophiostoma novo-ulmi subsp. novo-ulmi,* (G) *Ophiostoma ulmi,* (H) *Zymoseptoria tritici,* (I) *Ophiostoma novo-ulmi subsp. americana,* (J) *Magnaporthe oryzae Triticum,* (K) *Cercospora beticola,* (L) *Aspergillus flavus,* (M) *Neurospora discreta,* (N) *Sclerotinia sclerotiorum,* (O) *Verticillium dahliae,* (P) *Rhynchosporium commune,* (Q) *Pyrenophora teres f. sp. teres,* (R) *Magnaporthe oryzae rice,* (S) *Fusarium graminearum,* (T) *Cryphonectria parasitica*. Box-plot distribution was generated using 100 bootstrap replicates performed using the model with the lowest Akaike's information criterion.

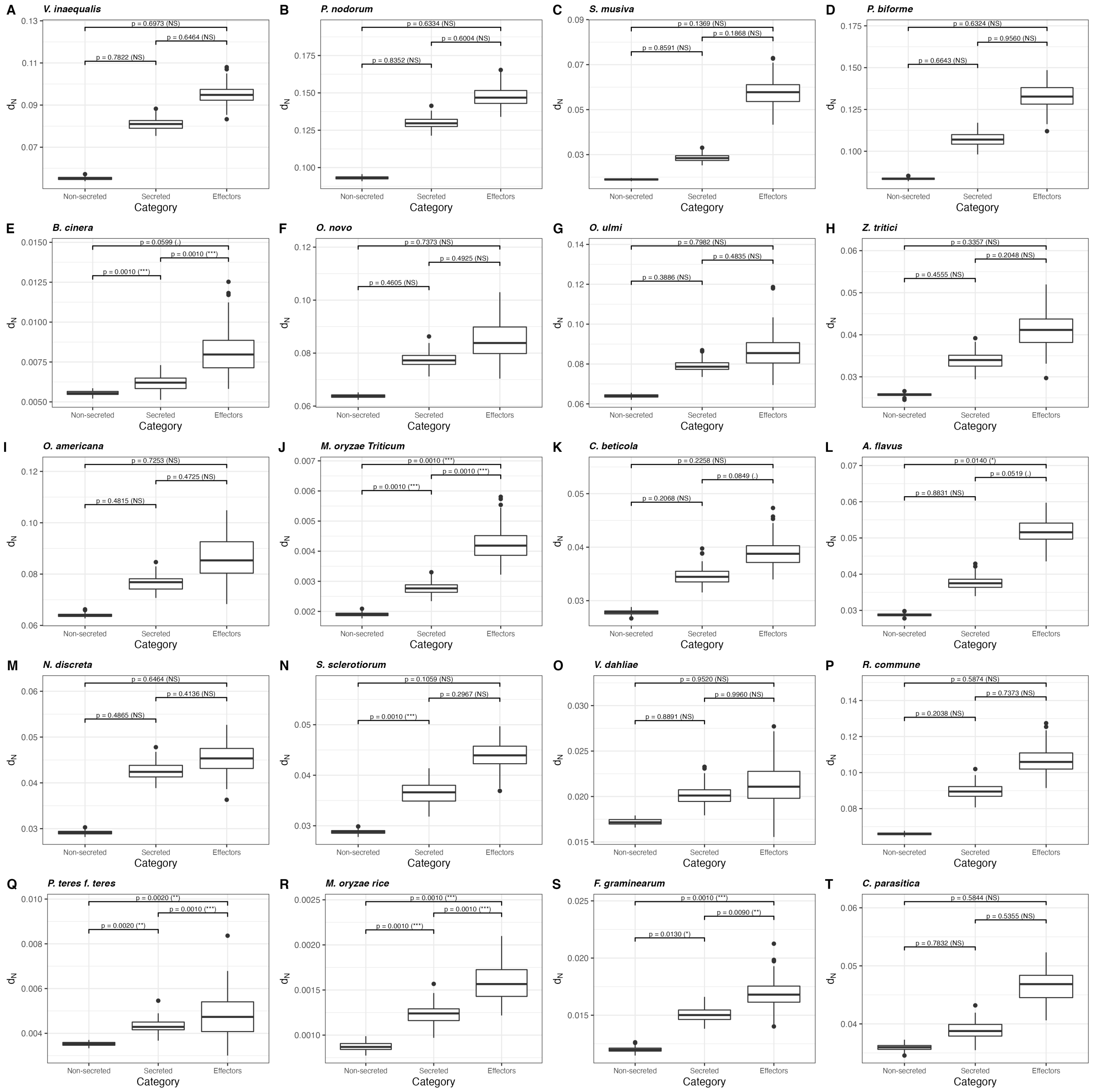


**Supplementary figure 11.** Variation in the number of nonsynonymous substitutions per site ($d_N$) across 20 fungal species. (A) *Venturia inaequalis,* (B) *Parastagonospora nodorum,* (C) *Sphaerulina musiva,* (D) *Penicillium biforme,* (E) *Botrytis cinerea,* (F) *Ophiostoma novo-ulmi subsp. novo-ulmi,* (G) *Ophiostoma ulmi,* (H) *Zymoseptoria tritici,* (I) *Ophiostoma novo-ulmi subsp. americana,* (J) *Magnaporthe oryzae Triticum,* (K) *Cercospora beticola,* (L) *Aspergillus flavus,* (M) *Neurospora discreta,* (N) *Sclerotinia sclerotiorum,* (O) *Verticillium dahliae,* (P) *Rhynchosporium commune,* (Q) *Pyrenophora teres f. sp. teres,* (R) *Magnaporthe oryzae rice,* (S) *Fusarium graminearum,* (T) *Cryphonectria parasitica*. Box-plot distribution was generated using 100 bootstrap replicates performed using the model with the lowest Akaike's information criterion.

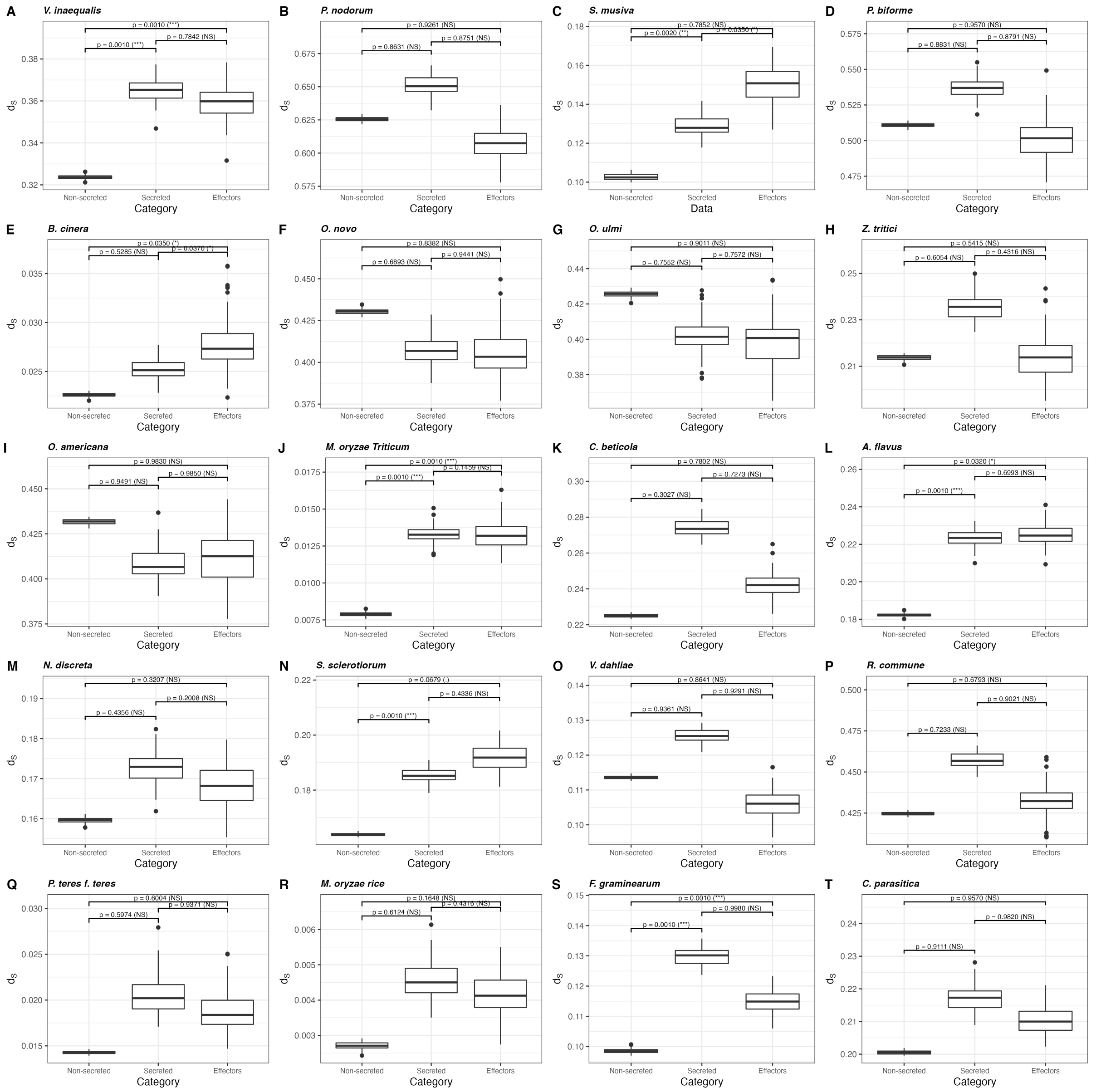


**Supplementary figure 12.** Variation in the number of synonymous substitutions per site ($d_S$) across 20 fungal species. (A) *Venturia inaequalis,* (B) *Parastagonospora nodorum,* (C) *Sphaerulina musiva,* (D) *Penicillium biforme,* (E) *Botrytis cinerea,* (F) *Ophiostoma novo-ulmi subsp. novo-ulmi,* (G) *Ophiostoma ulmi,* (H) *Zymoseptoria tritici,* (I) *Ophiostoma novo-ulmi subsp. americana,* (J) *Magnaporthe oryzae Triticum,* (K) *Cercospora beticola,* (L) *Aspergillus flavus,* (M) *Neurospora discreta,* (N) *Sclerotinia sclerotiorum,* (O) *Verticillium dahliae,* (P) *Rhynchosporium commune,* (Q) *Pyrenophora teres f. sp. teres,* (R) *Magnaporthe oryzae rice,* (S) *Fusarium graminearum,* (T) *Cryphonectria parasitica*. Box-plot distribution was generated using 100 bootstrap replicates performed using the model with the lowest Akaike's information criterion.

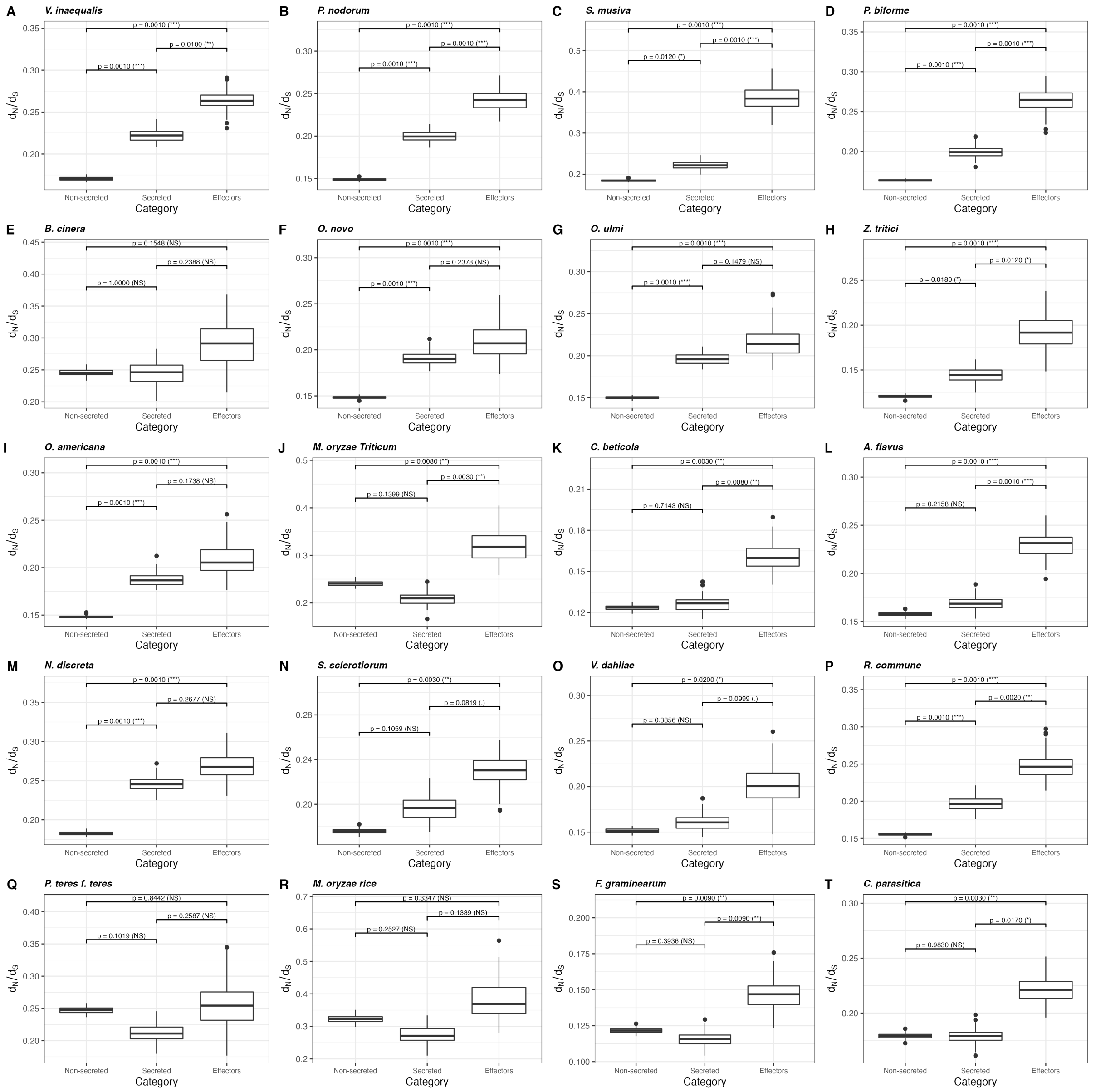


**Supplementary figure 13.** Variation in the rate of nonsynonymous substitutions relative to the rate of synonymous substitutions ($\omega=d_N/d_S$) across 20 fungal species. (A) *Venturia inaequalis,* (B) *Parastagonospora nodorum,* (C) *Sphaerulina musiva,* (D) *Penicillium biforme,* (E) *Botrytis cinerea,* (F) *Ophiostoma novo-ulmi subsp. novo-ulmi,* (G) *Ophiostoma ulmi,* (H) *Zymoseptoria tritici,* (I) *Ophiostoma novo-ulmi subsp. americana,* (J) *Magnaporthe oryzae Triticum,* (K) *Cercospora beticola,* (L) *Aspergillus flavus,* (M) *Neurospora discreta,* (N) *Sclerotinia sclerotiorum,* (O) *Verticillium dahliae,* (P) *Rhynchosporium commune,* (Q) *Pyrenophora teres f. sp. teres,* (R) *Magnaporthe oryzae rice,* (S) *Fusarium graminearum,* (T) *Cryphonectria parasitica*. Box-plot distribution was generated using 100 bootstrap replicates performed using the model with the lowest Akaike's information criterion.

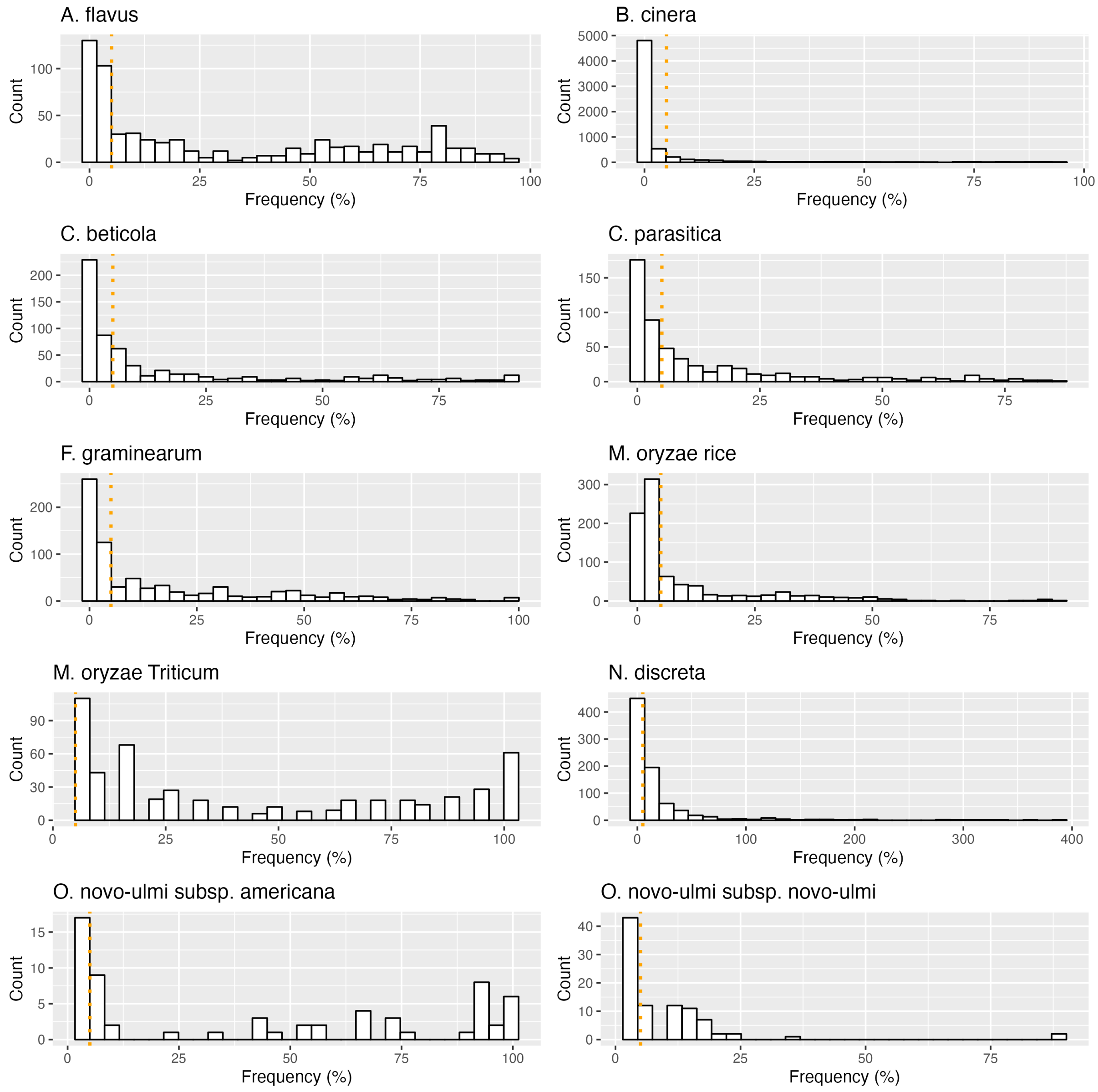


**Supplementary figure 14.** Distribution of deletion frequencies per species. Deletion frequencies were calculated for each gene as the number of isolates missing that specific gene divided by the total number of isolates. Genes coding secreted proteins or effectors are not included. The dotted orange line represents the 5% maximum missingness threshold used to select genes in presence absence polymorphism.

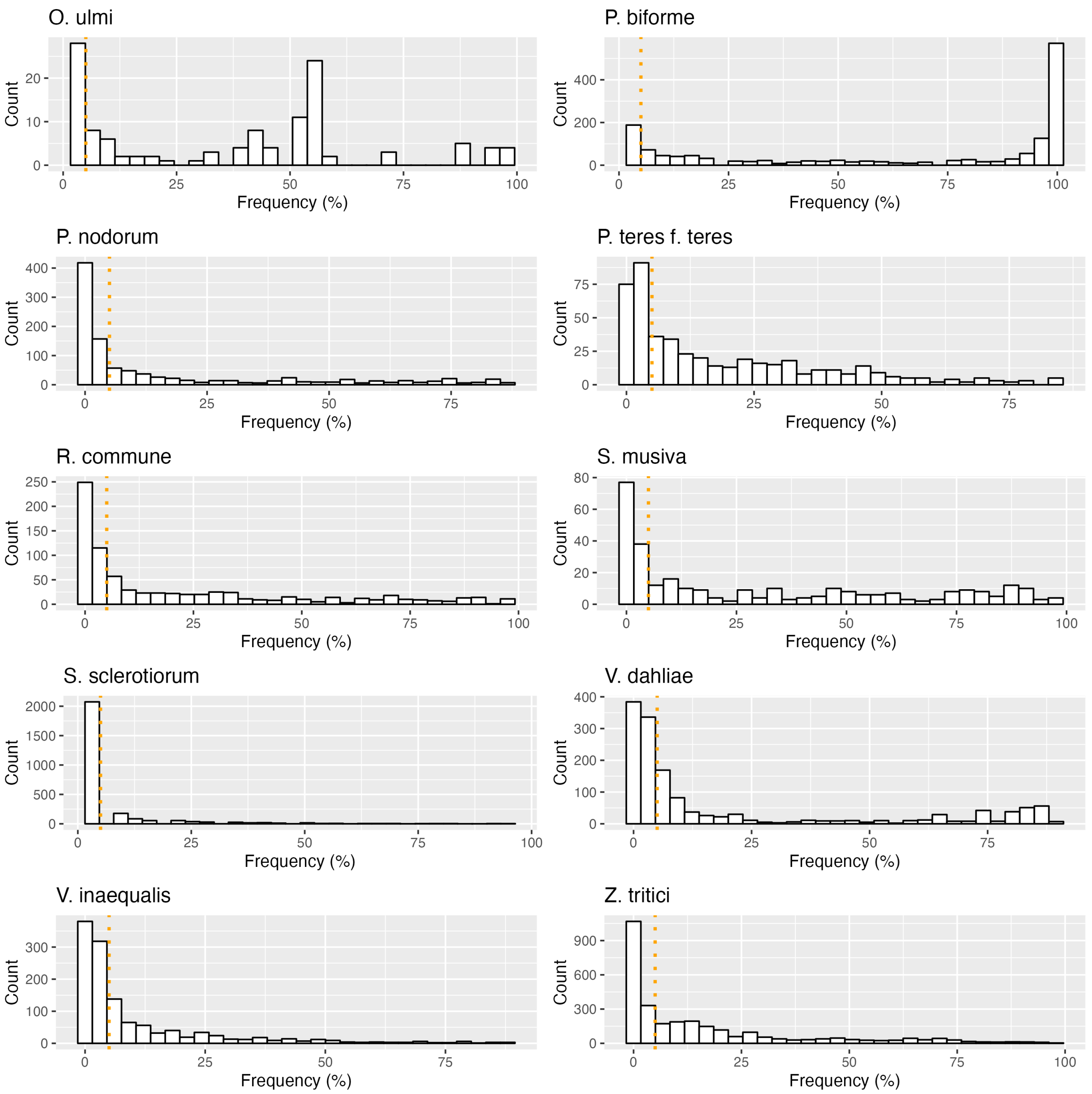


**Supplementary figure 14 (continuation).** Distribution of deletion frequencies per species. Deletion frequencies were calculated for each gene as the number of isolates missing that specific gene divided by the total number of isolates. Genes coding secreted proteins or effectors are not included. The dotted orange line represents the 5% maximum missingness threshold used to select genes in presence absence polymorphism.

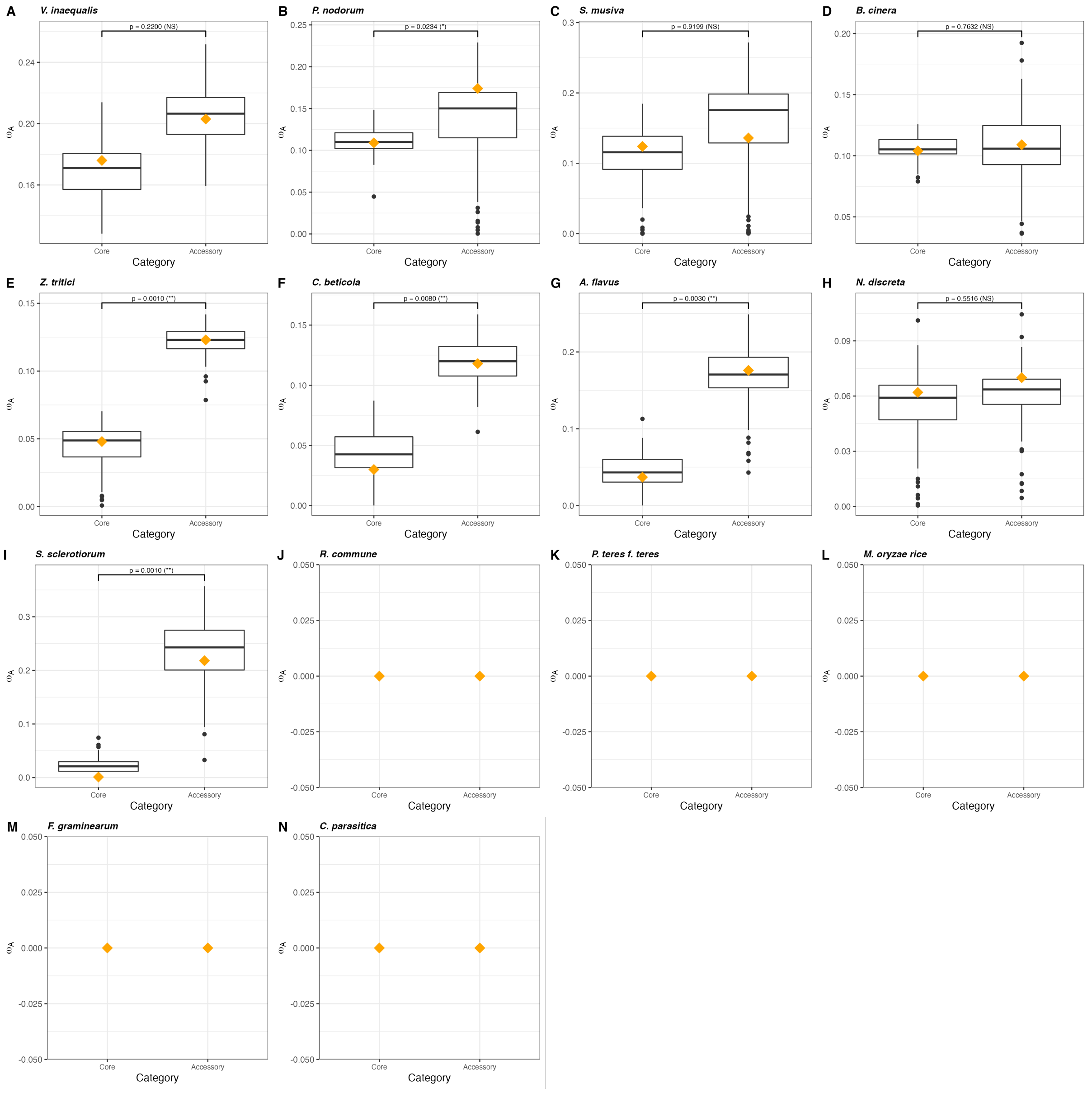


**Supplementary figure 15.** Variation in adaptive nonsynonymous substitutions rate ($\omega_A$) between core and accessory gene datasets within 14 fungal species. (A) *Venturia inaequalis,* (B) *Parastagonospora nodorum,* (C) *Sphaerulina musiva,* (D) *Botrytis cinerea,* (E) *Zymoseptoria tritici,* (F) *Cercospora beticola,* (G) *Aspergillus flavus,* (H) *Neurospora discreta,* (I) *Sclerotinia sclerotiorum,* (J) *Rhynchosporium commune,* (K) *Pyrenophora teres f. sp. teres,* (L) *Magnaporthe oryzae rice,* (M) *Fusarium graminearum,* (N) *Cryphonectria parasitica*. Box-plot distribution was generated using 100 bootstrap replicates performed using the model with the lowest Akaike's information criterion. The orange diamond represents the mean value calculated after averaging over all models of distribution of fitness effects.

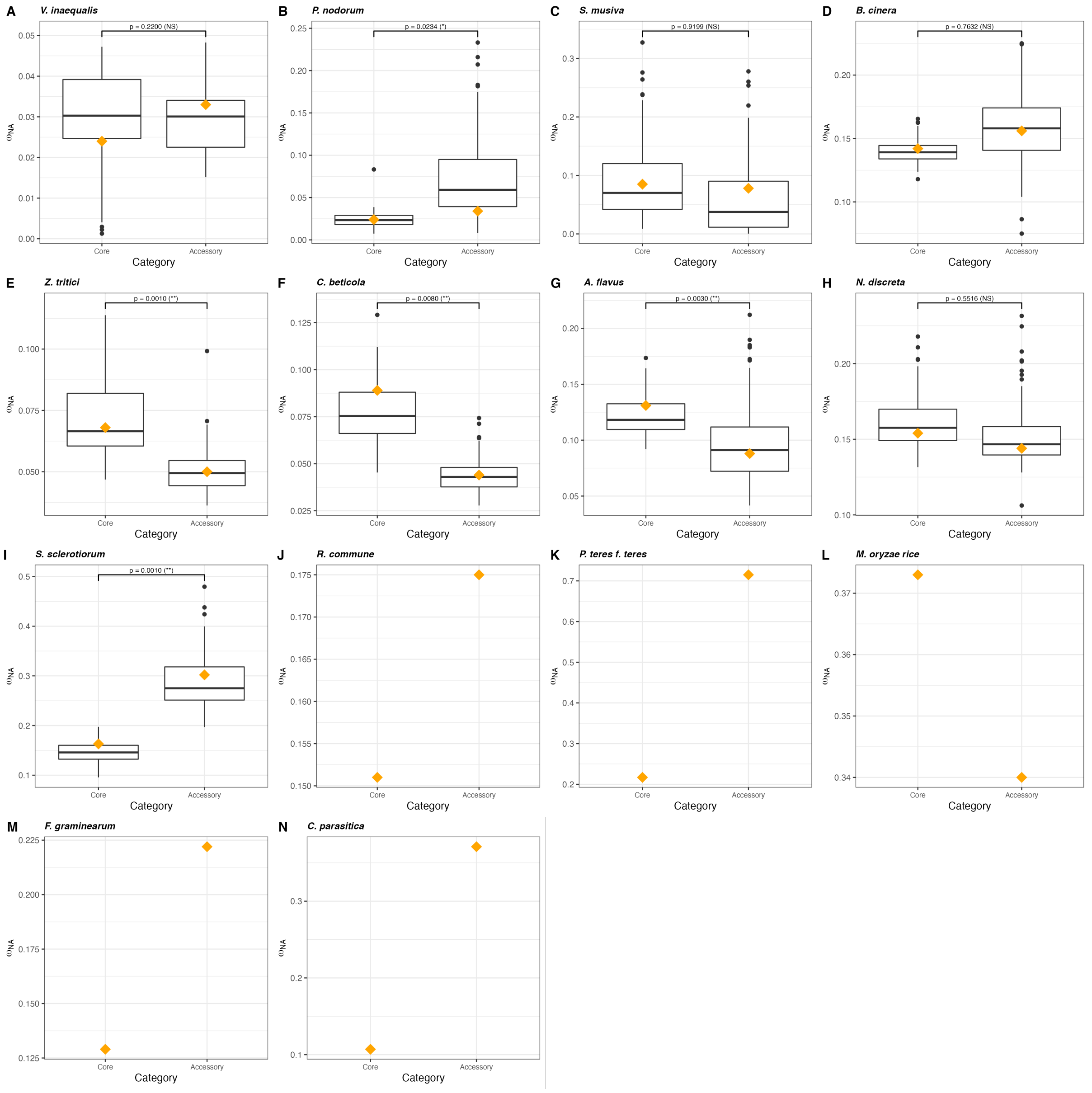


**Supplementary figure 16.** Variation in nonadaptive nonsynosnymous substitutions rate ($\omega_{NA}$) between core and accessory gene datasets within 14 fungal species. (A) *Venturia inaequalis,* (B) *Parastagonospora nodorum,* (C) *Sphaerulina musiva,* (D) *Botrytis cinerea,* (E) *Zymoseptoria tritici,* (F) *Cercospora beticola,* (G) *Aspergillus flavus,* (H) *Neurospora discreta,* (I) *Sclerotinia sclerotiorum,* (J) *Rhynchosporium commune,* (K) *Pyrenophora teres f. sp. teres,* (L) *Magnaporthe oryzae rice,* (M) *Fusarium graminearum,* (N) *Cryphonectria parasitica*. Box-plot distribution was generated using 100 bootstrap replicates performed using the model with the lowest Akaike's information criterion. The orange diamond represents the mean value calculated after averaging over all models of distribution of fitness effects.

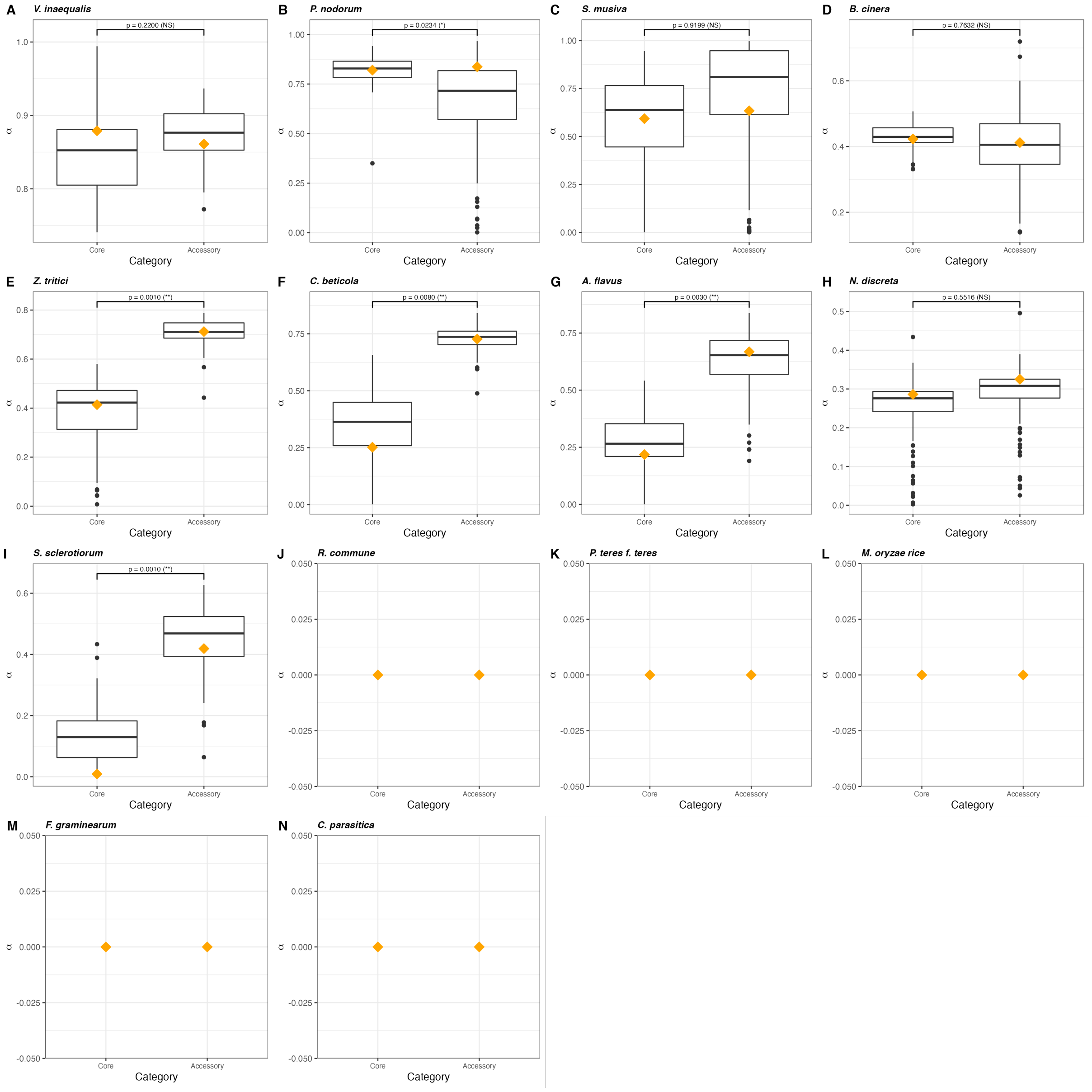


**Supplementary figure 17.** Variation in the proportion of amino-acid substitutions that are adaptive (α) between core genes and accessory genes across 14 fungal species. (A) *Venturia inaequalis,* (B) *Parastagonospora nodorum,* (C) *Sphaerulina musiva,* (D) *Botrytis cinerea,* (E) *Zymoseptoria tritici,* (F) *Cercospora beticola,* (G) *Aspergillus flavus,* (H) *Neurospora discreta,* (I) *Sclerotinia sclerotiorum,* (J) *Rhynchosporium commune,* (K) *Pyrenophora teres f. sp. teres,* (L) *Magnaporthe oryzae rice,* (M) *Fusarium graminearum,* (N) *Cryphonectria parasitica*. Box-plot distribution was generated using 100 bootstrap replicates performed using the model with the lowest Akaike's information criterion. The orange diamond represents the mean value calculated after averaging over all models of distribution of fitness effects.

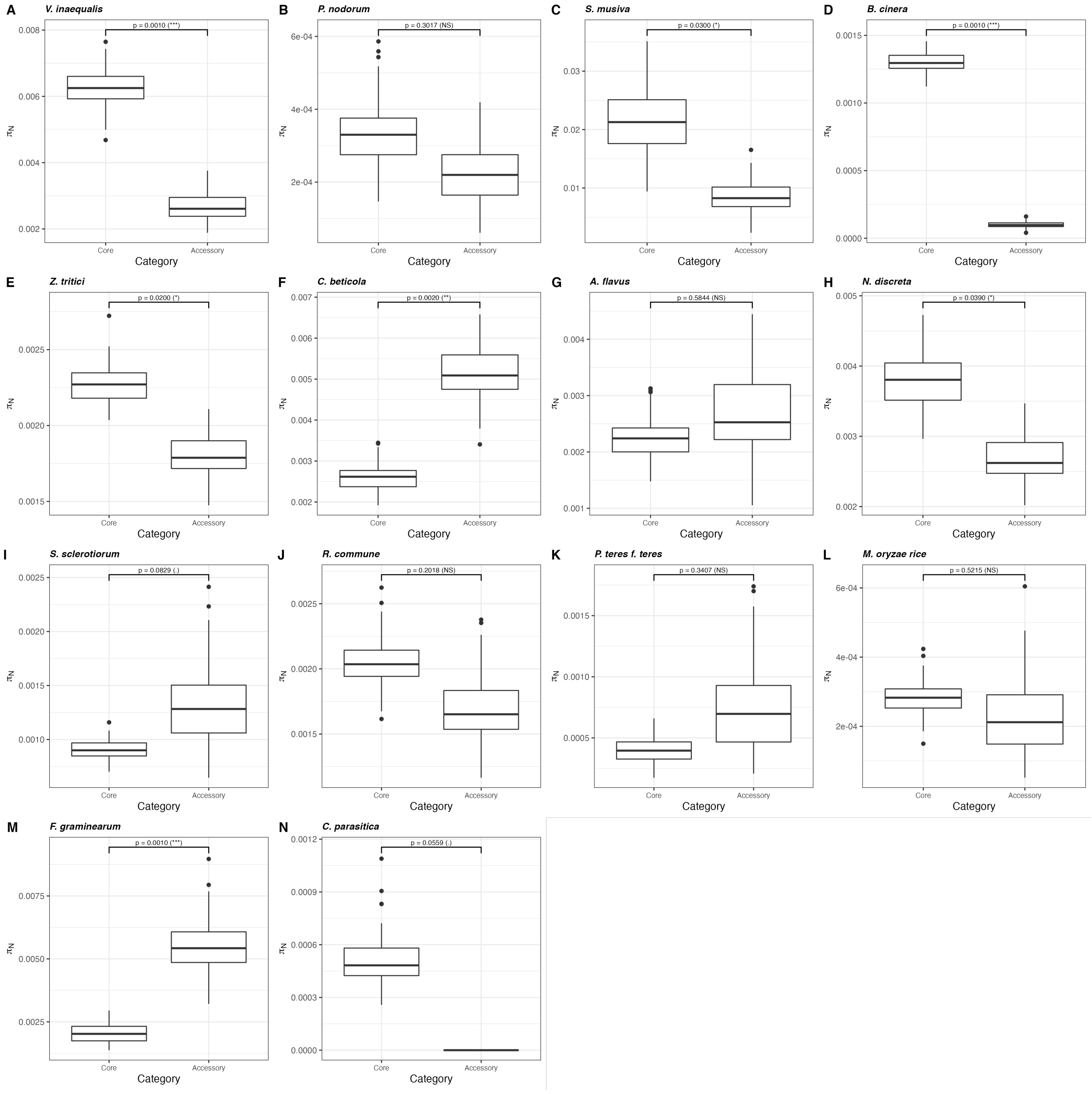


**Supplementary figure 18.** Variation in the nonsynonymous diversity ($\pi_N$) between core genes and accessory genes across 14 fungal species. (A) *Venturia inaequalis,* (B) *Parastagonospora nodorum,* (C) *Sphaerulina musiva,* (D) *Botrytis cinerea,* (E) *Zymoseptoria tritici,* (F) *Cercospora beticola,* (G) *Aspergillus flavus,* (H) *Neurospora discreta,* (I) *Sclerotinia sclerotiorum,* (J) *Rhynchosporium commune,* (K) *Pyrenophora teres f. sp. teres,* (L) *Magnaporthe oryzae rice,* (M) *Fusarium graminearum,* (N) *Cryphonectria parasitica*. Box-plot distribution was generated using 100 bootstrap replicates performed using the model with the lowest Akaike's information criterion.

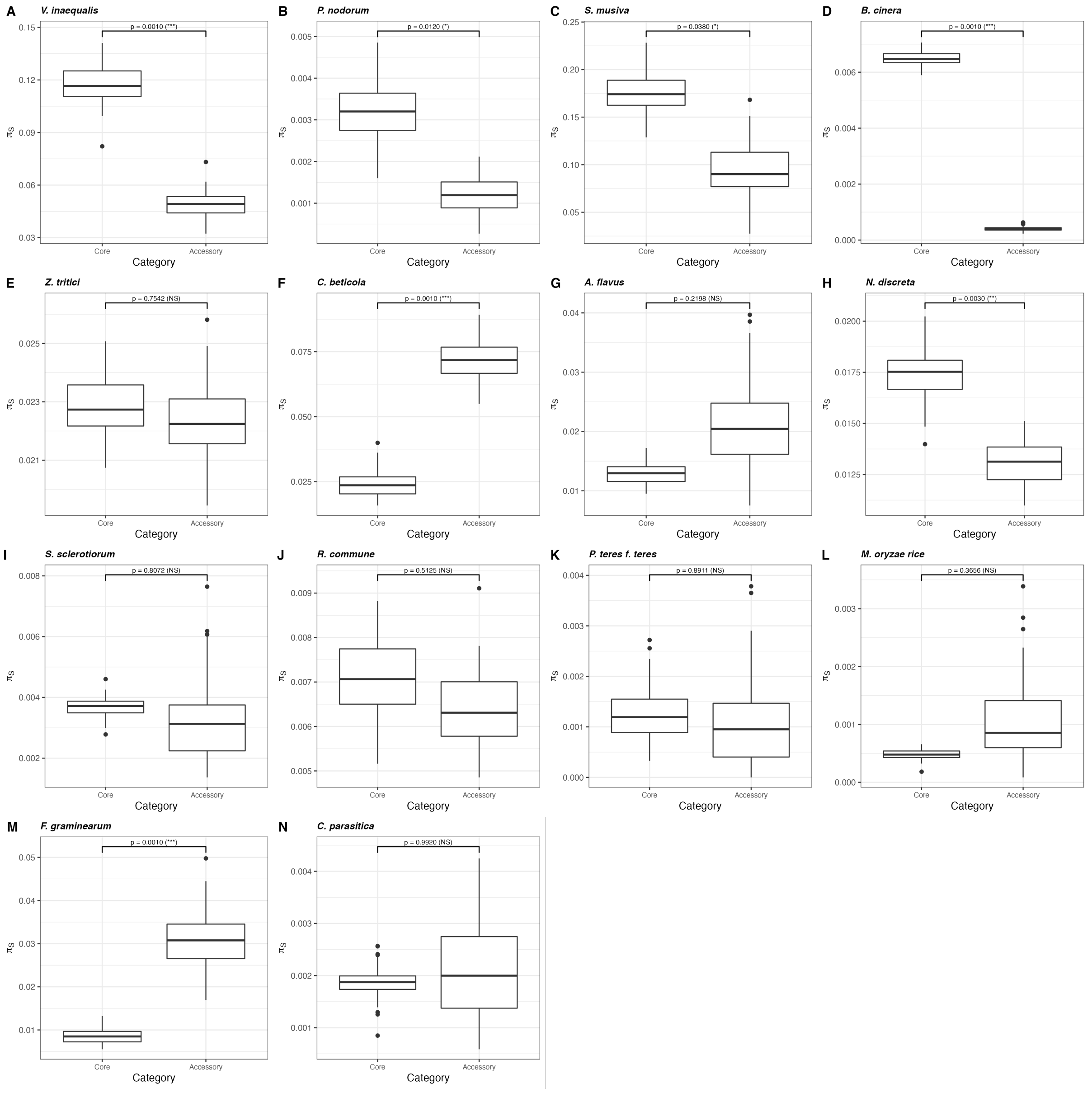


**Supplementary figure 19.** Variation in the synonymous diversity ($\pi_S$) between core genes and accessory genes across 14 fungal species. (A) *Venturia inaequalis,* (B) *Parastagonospora nodorum,* (C) *Sphaerulina musiva,* (D) *Botrytis cinerea,* (E) *Zymoseptoria tritici,* (F) *Cercospora beticola,* (G) *Aspergillus flavus,* (H) *Neurospora discreta,* (I) *Sclerotinia sclerotiorum,* (J) *Rhynchosporium commune,* (K) *Pyrenophora teres f. sp. teres,* (L) *Magnaporthe oryzae rice,* (M) *Fusarium graminearum,* (N) *Cryphonectria parasitica*. Box-plot distribution was generated using 100 bootstrap replicates performed using the model with the lowest Akaike's information criterion.

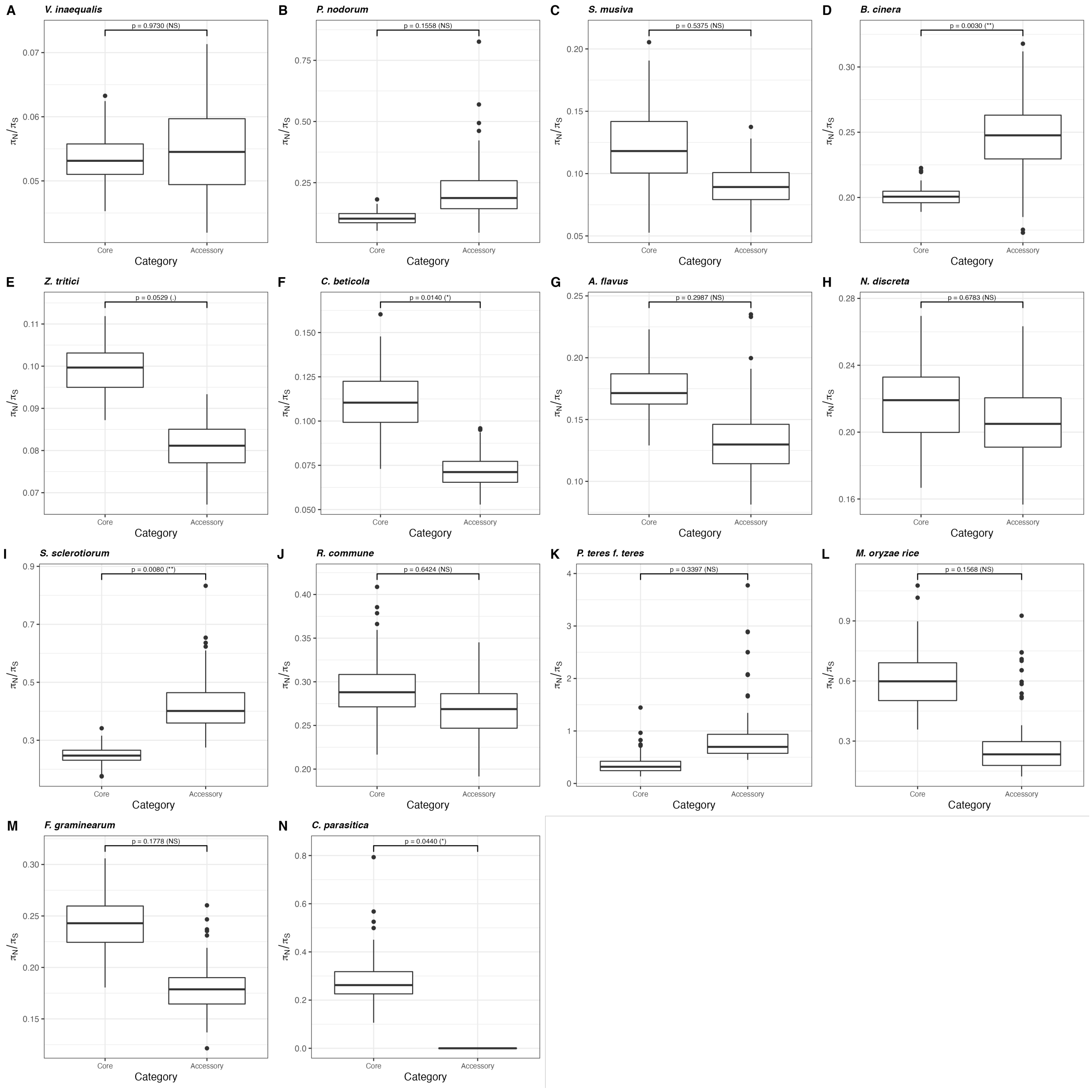


**Supplementary figure 20.** Variation in the ratio of the mean number of nonsynonymous nucleotide differences and mean number of synonymous nucleotide differences ($\pi_N/\pi_S$) between core genes and accessory genes across 14 fungal species. (A) *Venturia inaequalis,* (B) *Parastagonospora nodorum,* (C) *Sphaerulina musiva,* (D) *Botrytis cinerea,* (E) *Zymoseptoria tritici,* (F) *Cercospora beticola,* (G) *Aspergillus flavus,* (H) *Neurospora discreta,* (I) *Sclerotinia sclerotiorum,* (J) *Rhynchosporium commune,* (K) *Pyrenophora teres f. sp. teres,* (L) *Magnaporthe oryzae rice,* (M) *Fusarium graminearum,* (N) *Cryphonectria parasitica*. Box-plot distribution was generated using 100 bootstrap replicates performed using the model with the lowest Akaike's information criterion.

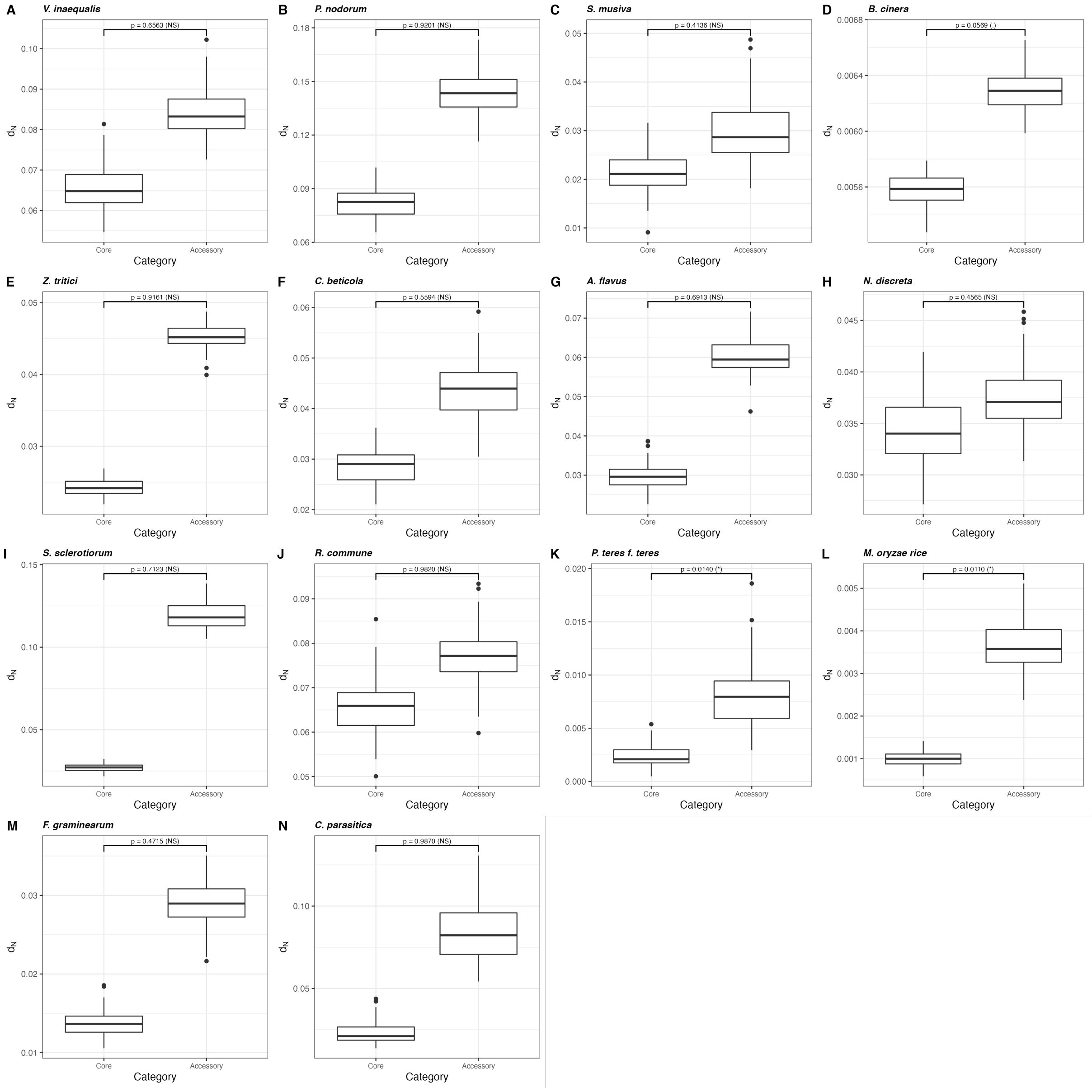


**Supplementary figure 21.** Variation in the number of nonsynonymous substitutions per site ($d_N$) between core genes and accessory genes across 14 fungal species. (A) *Venturia inaequalis,* (B) *Parastagonospora nodorum,* (C) *Sphaerulina musiva,* (D) *Botrytis cinerea,* (E) *Zymoseptoria tritici,* (F) *Cercospora beticola,* (G) *Aspergillus flavus,* (H) *Neurospora discreta,* (I) *Sclerotinia sclerotiorum,* (J) *Rhynchosporium commune,* (K) *Pyrenophora teres f. sp. teres,* (L) *Magnaporthe oryzae rice,* (M) *Fusarium graminearum,* (N) *Cryphonectria parasitica*. Box-plot distribution was generated using 100 bootstrap replicates performed using the model with the lowest Akaike's information criterion.

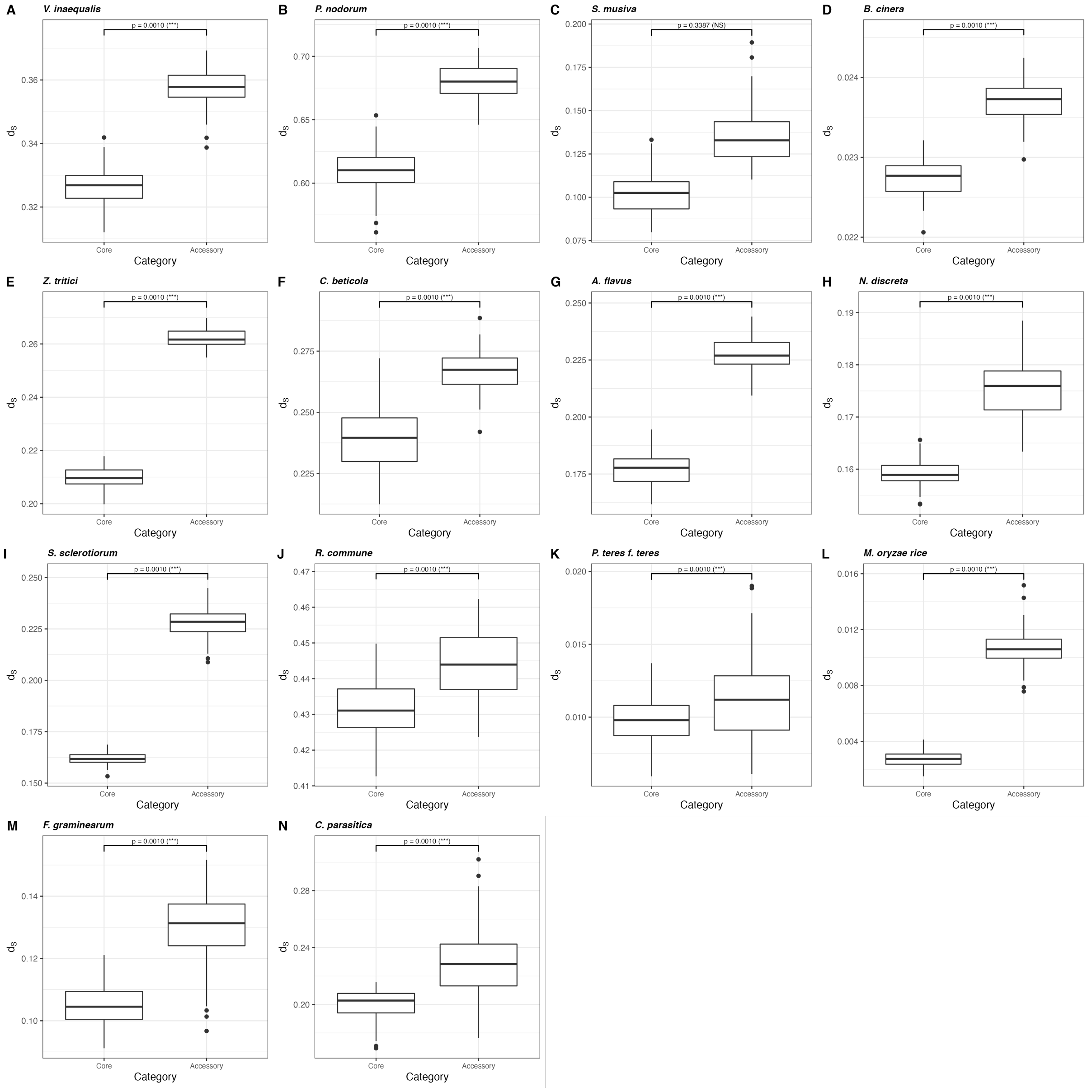


**Supplementary figure 22.** Variation in the number of synonymous substitutions per site ($d_S$) between core genes and accessory genes across 14 fungal species. (A) *Venturia inaequalis,* (B) *Parastagonospora nodorum,* (C) *Sphaerulina musiva,* (D) *Botrytis cinerea,* (E) *Zymoseptoria tritici,* (F) *Cercospora beticola,* (G) *Aspergillus flavus,* (H) *Neurospora discreta,* (I) *Sclerotinia sclerotiorum,* (J) *Rhynchosporium commune,* (K) *Pyrenophora teres f. sp. teres,* (L) *Magnaporthe oryzae rice,* (M) *Fusarium graminearum,* (N) *Cryphonectria parasitica*. Box-plot distribution was generated using 100 bootstrap replicates performed using the model with the lowest Akaike's information criterion.

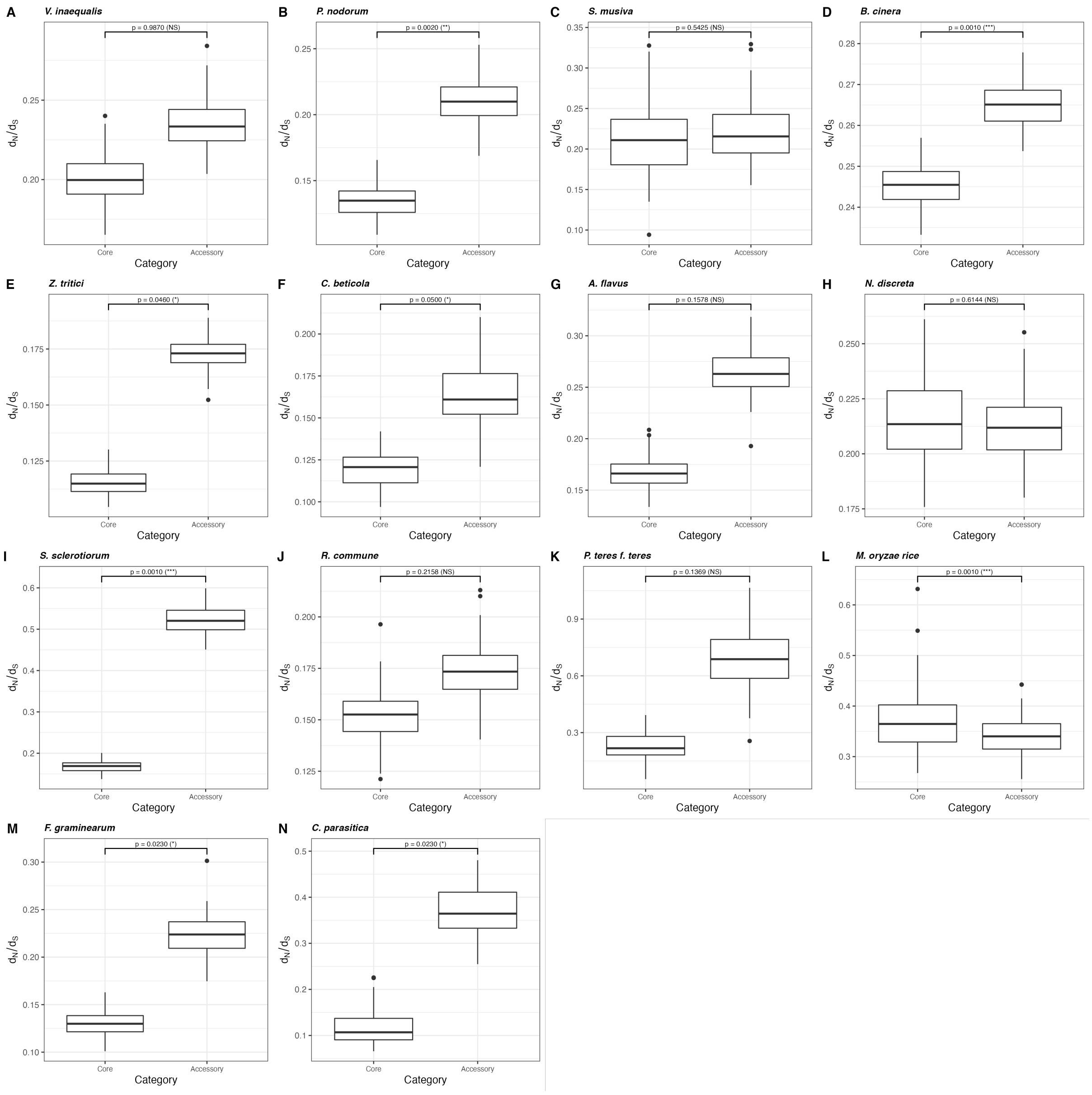


**Supplementary figure 23.** Variation in the rate of nonsynonymous substitutions relative to the rate of synonymous substitutions ($\omega = d_N/d_S$) between core genes and accessory genes across 14 fungal species. (A) *Venturia inaequalis,* (B) *Parastagonospora nodorum,* (C) *Sphaerulina musiva,* (D) *Botrytis cinerea,* (E) *Zymoseptoria tritici,* (F) *Cercospora beticola,* (G) *Aspergillus flavus,* (H) *Neurospora discreta,* (I) *Sclerotinia sclerotiorum,* (J) *Rhynchosporium commune,* (K) *Pyrenophora teres f. sp. teres,* (L) *Magnaporthe oryzae rice,* (M) *Fusarium graminearum,* (N) *Cryphonectria parasitica*. Box-plot distribution was generated using 100 bootstrap replicates performed using the model with the lowest Akaike's information criterion.
