## Supplementary material for "Large-scale analyses reveal the contribution of adaptive evolution in pathogenic and non-pathogenic fungal species": Simulations

### Assessing the impact of population structure on the estimation of the rate of adaptive substitutions

2022-08-19

A population of 10,000 individuals is simulated for 100,000 generations, the first 10,000 generations serving as a burnin, in which no substitution is counted. The simulated "genome" is made of 1,500 genes of 500 bp each, separated by 500 bp of non coding spacers. The recombination rate is uniform along the sequence and set to 1e-7. The mutation rate, also uniform, is set to 2.2e-7. Only mutations within "genes" are recorded. There are three types of mutations, all semi-dominant:

- Mutations $m1$ are neutral (fitness $= 0$), occurring with a probability of 0.25.
- Mutations $m2$ are deleterious, with a fitness taken from a negative gamma distribution of mean -2.5 and shape 0.3. These mutations occur with a probability of $0.75 - p$.
- Mutations $m3$ are advantageous, with a fitness taken from an exponential distribution with mean 0.01, occurring with a probability $p$.

When estimating $\alpha$, $m1$ mutations are the neutral reference (equivalent to synonymous mutations) and mutations $m2$ and $m3$ are the selected mutations (equivalent to non-synonymous mutations). We output the SFS after 100,000 generations for a sample of 20 diploid individuals, and count substitutions in the last 90,000 generations. The values used to estimate parameters are the true ones, we do not assess the impact of possible errors in reconstructing the SFS or divergence values. While only sequencing and calling errors may bias the inferred SFS, estimated divergence parameters may be different from the true ones, since they are computed from an outgroup species. Therefore, the estimation errors of $\alpha$ are only due to the method of calculation per se.

#### Load data

```
dat <- read.table("SimulationResults.csv", sep = ",", header = TRUE)
dat$PropPosSelFactor <- as.ordered(dat$PropPosSel) # For plotting
dat$PropPosSelOrder <- as.numeric(dat$PropPosSelFactor) # For smoothing
library(tidyr)
library(ggplot2)
library(cowplot)
```

#### Alpha as a function of the proportion of adaptive substitutions

Because we would like to assess the inference accuracy for distinct values of $\alpha$, we varied the proportion of positively selected mutations ($p$) in the simulations. We first check how $\alpha$ varies with $p$ (Figure 1):

```
p.alpha <- ggplot(subset(dat, Scenario == "Panmictic"),
            aes(x = PropPosSel, y = Alpha.simulated)) +
  geom_point() + geom_smooth(method = "nls",
                             formula = y ~ a*log(x+exp(b))+c,
                             method.args = list(
                               start = c(a = 1, b = -3, c = 0),
                               algorithm = "port"),
                             se = FALSE) +
  xlab("Proportion of adaptive mutations") +
```

```r
  ylab(expression(alpha)) +
  theme_bw()

p.omega <- ggplot(subset(dat, Scenario == "Panmictic"),
            aes(x = PropPosSel, y = dNdS)) +
  geom_point() + geom_smooth(method = "nls",
                             formula = y ~ a*x^b+c,
                             method.args = list(
                               start = c(a = 1, b = 0.1, c = 0),
                               algorithm = "port"),
                             se = FALSE) +
  xlab("Proportion of adaptive mutations") +
  ylab(expression(omega)) +
  theme_bw()

p.omegaA <- ggplot(subset(dat, Scenario == "Panmictic"),
            aes(x = PropPosSel, y = OmegaA.simulated)) +
  geom_point() + geom_smooth(method = "nls",
                             formula = y ~ a*x^b,
                             method.args = list(
                               start = c(a = 1, b = 0.1),
                               algorithm = "port"),
                             se = FALSE) +
  xlab("Proportion of adaptive mutations") +
  ylab(expression(omega[A])) +
  theme_bw()

p.omegaNA <- ggplot(subset(dat, Scenario == "Panmictic"),
            aes(x = PropPosSel, y = OmegaNA.simulated)) +
  geom_point() + geom_smooth(method = "nls",
                             formula = y ~ a*x^b+c,
                             method.args = list(
                               start = c(a = 1, b = 0.1, c = 0),
                               algorithm = "port"),
                             se = FALSE) +
  xlab("Proportion of adaptive mutations") +
  ylab(expression(omega["NA"])) +
  theme_bw()

plot_grid(p.alpha, p.omega, p.omegaA, p.omegaNA, nrow = 2, labels = "AUTO")
```

There are positive relationships between $p$ and the $\alpha$ (logarithmic function) and $\omega$s (power function). The positive correlation between $p$ and $\omega_{NA}$ may seem at first glance unexpected: in the simulations, the proportion of non-adaptive mutations is 0.75 - p, so their should be a negative relationship. A possible explanation is that more deleterious mutations are fixed in the presence of adaptive ones because of linkage, which is quite strong with our simulation parameters (small intergenic regions + low recombination rate).

#### Panmictic population

We first assess the estimation of parameters in the ideal case of a panmictic population with constant size. We compare three types of estimates:

- Classic: $\alpha = 1 - \frac{p_N/p_S}{d_N/d_S}$, $\omega_A = d_N/d_S - p_N/p_S$, $\omega_{NA} = p_N/p_S$,
- FWW: Fay, Wickoff and Wu's version of these estimates,

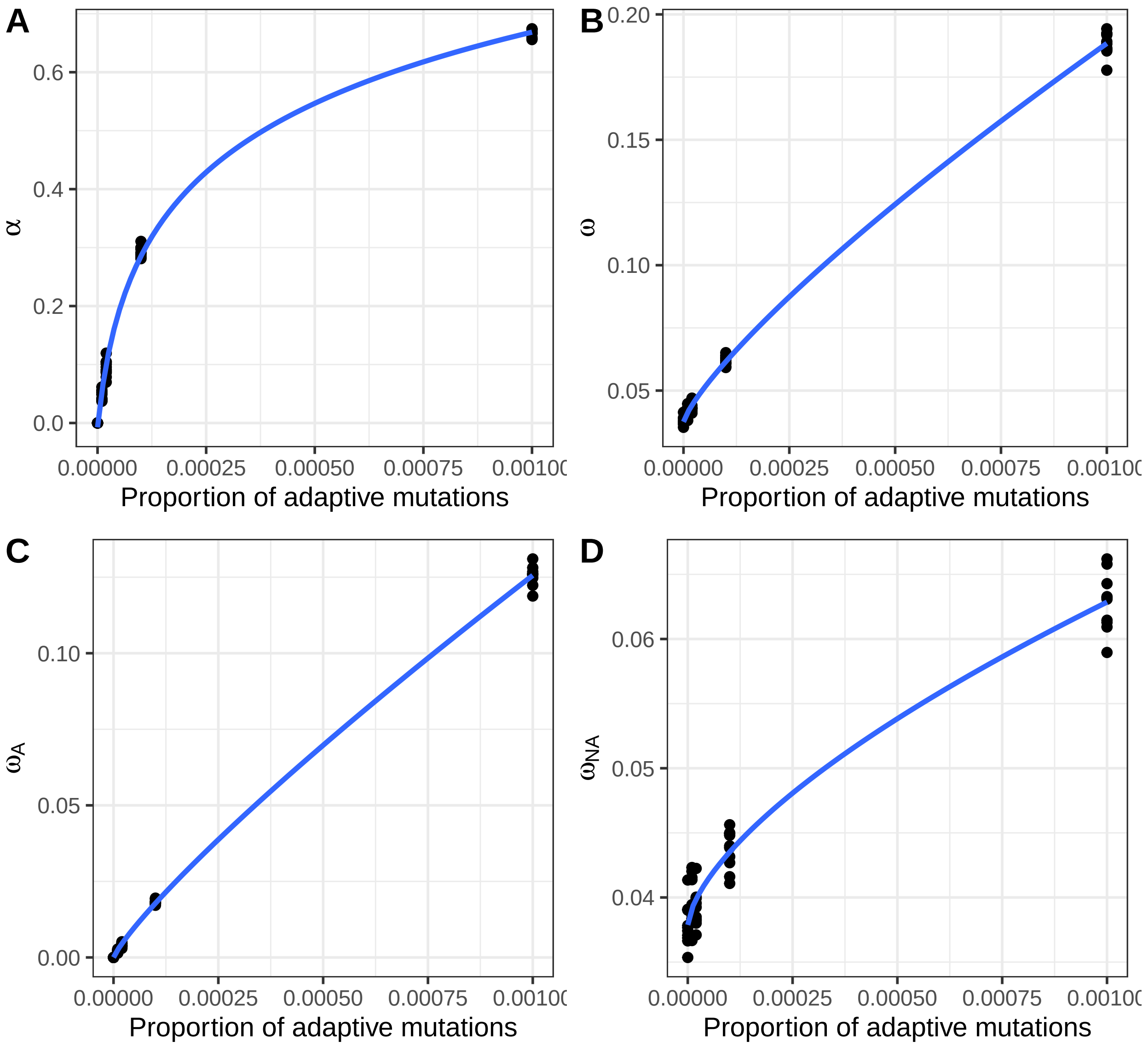


Figure 1: Real parameters as a function of $p$

- GammaExpo: estimates obtained by Grapes after fitting a Gamma+Exponential DFE.

Select and reorganize the data:

```
dat.pan.alpha <- subset(dat, Scenario == "Panmictic") %>% pivot_longer(
  cols = c("Alpha.basic", "Alpha.FWW", "Alpha.inferred"),
  names_to = "AlphaType", values_to = "Alpha")
dat.pan.alpha$AlphaType <- factor(dat.pan.alpha$AlphaType,
  levels = c("Alpha.basic", "Alpha.FWW", "Alpha.inferred"),
  labels = c("Basic", "FWW", "GammaExpo"))

dat.pan.omegaA <- subset(dat, Scenario == "Panmictic") %>% pivot_longer(
  cols = c("OmegaA.basic", "OmegaA.FWW", "OmegaA.inferred"),
  names_to = "AlphaType", values_to = "OmegaA")
dat.pan.omegaA$AlphaType <- factor(dat.pan.omegaA$AlphaType,
  levels = c("OmegaA.basic", "OmegaA.FWW", "OmegaA.inferred"),
  labels = c("Basic", "FWW", "GammaExpo"))

dat.pan.omegaNA <- subset(dat, Scenario == "Panmictic") %>% pivot_longer(
  cols = c("OmegaNA.basic", "OmegaNA.FWW", "OmegaNA.inferred"),
  names_to = "AlphaType", values_to = "OmegaNA")
dat.pan.omegaNA$AlphaType <- factor(dat.pan.omegaNA$AlphaType,
  levels = c("OmegaNA.basic", "OmegaNA.FWW", "OmegaNA.inferred"),
  labels = c("Basic", "FWW", "GammaExpo"))
```

Then plot the results. For each parameter, we plot the error (estimated minus real value) as a function of the proportion of positively selected mutations (Figure 2):

```
p.alpha <- ggplot(dat.pan.alpha,
  aes(x = PropPosSelFactor, y = Alpha - Alpha.simulated)) +
  geom_point() + geom_abline(slope = 0) +
  geom_smooth(aes(x = PropPosSelOrder), method = "loess", se = FALSE) +
  facet_grid(~AlphaType) +
  ylab(expression(alpha - alpha^simulated)) +
  xlab("Proportion of positively selected mutations") +
  theme_bw()

p.omegaA <- ggplot(dat.pan.omegaA,
  aes(x = PropPosSelFactor, y = OmegaA - OmegaA.simulated)) +
  geom_point() + geom_abline(slope = 0) +
  geom_smooth(aes(x = PropPosSelOrder), method = "loess", se = FALSE) +
  facet_grid(~AlphaType) +
  ylab(expression(omega[A] - omega[A]^simulated)) +
  xlab("Proportion of positively selected mutations") +
  theme_bw()

p.omegaNA <- ggplot(dat.pan.omegaNA,
  aes(x = PropPosSelFactor, y = OmegaNA - OmegaNA.simulated)) +
  geom_point() + geom_abline(slope = 0) +
  geom_smooth(aes(x = PropPosSelOrder), method = "loess", se = FALSE) +
  facet_grid(~AlphaType) +
  ylab(expression(omega['NA'] - omega['NA']^simulated)) +
  xlab("Proportion of positively selected mutations") +
  theme_bw()

plot_grid(p.alpha, p.omegaA, p.omegaNA, nrow = 3, labels = "AUTO")
```

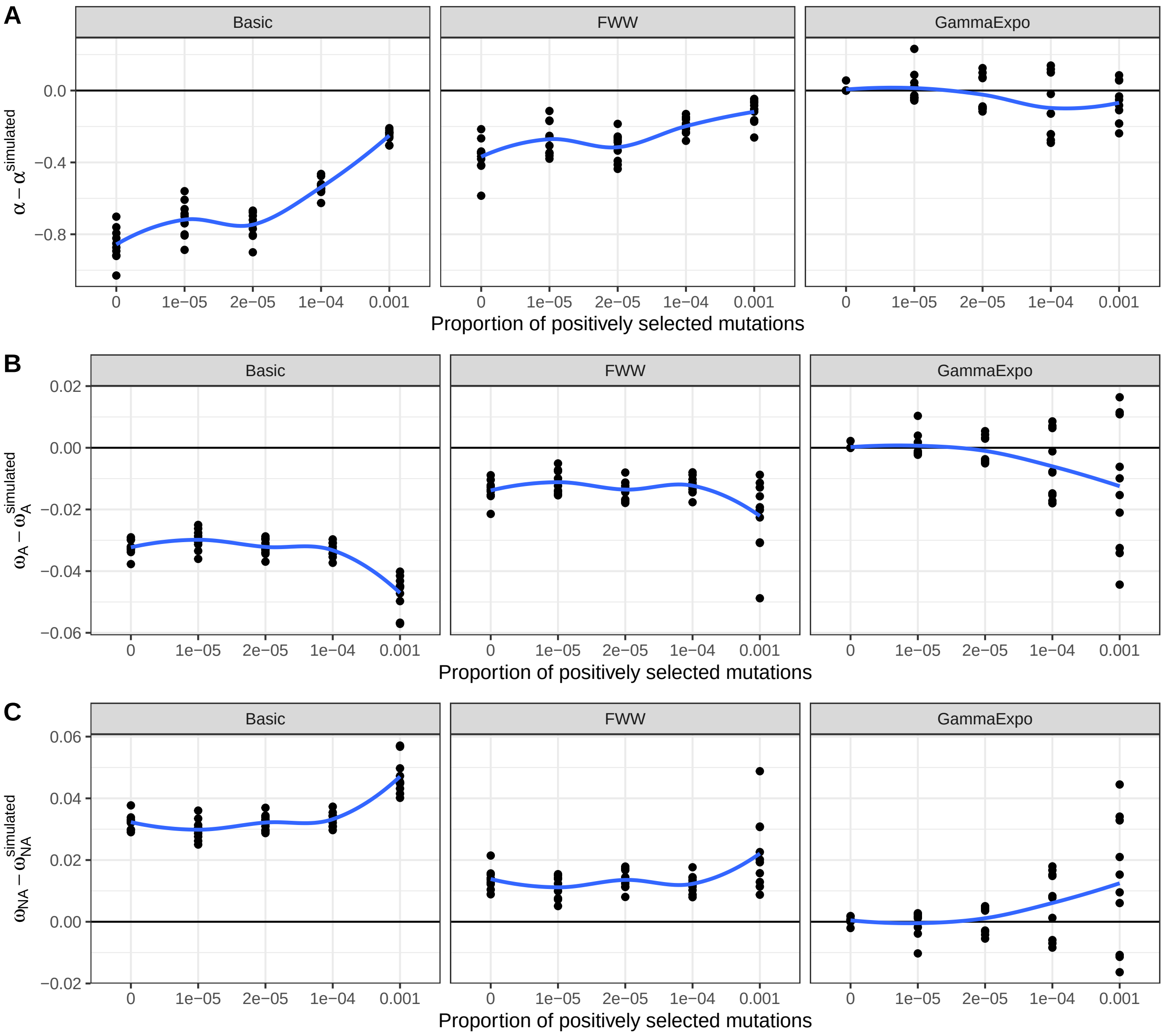


Figure 2: Panmictic population with constant size

We can see that:

- The classic estimates underestimate $\omega_A$ and overestimate $\omega_{NA}$.
- The biases are larger when $p$ is high, but interestingly, the opposite pattern is seen for $\alpha$: it is generally underestimated, but more so when $p$ is low.
- The FWW estimates show the same trend, but reduced. Notably, the biases are more consistent across values of $p$.
- The model-based inference provides mostly unbiased estimates, although some small biases remain for very high proportions of adaptive mutations.
- The estimation variance of model-based estimates increases when the proportion of adaptive mutations is high.

#### Panmictic population with exponential growth / shrinkage

After 10,000 generations (burnin period), the population size changes exponentially from 10,000 to a new size: 1,000, 5,000, 20,000 or 100,000. (In addition to the case where $N$ stays constant at 10,000.) We note as 'N ratio' or 'Size ratio' the ratio of the final $N$ over the initial one.

```
dat$NRatio <- as.ordered(round(log10(dat$NewPopSize/10000), 2))

dat.exp.alpha <- subset(dat, Scenario == "Exponential") %>% pivot_longer(
  cols = c("Alpha.basic", "Alpha.FWW", "Alpha.inferred"),
  names_to = "AlphaType", values_to = "Alpha")
dat.exp.alpha$AlphaType <- factor(dat.exp.alpha$AlphaType,
  levels = c("Alpha.basic", "Alpha.FWW", "Alpha.inferred"),
  labels = c("Basic", "FWW", "GammaExpo"))

dat.exp.omegaA <- subset(dat, Scenario == "Exponential") %>% pivot_longer(
  cols = c("OmegaA.basic", "OmegaA.FWW", "OmegaA.inferred"),
  names_to = "AlphaType", values_to = "OmegaA")
dat.exp.omegaA$AlphaType <- factor(dat.exp.omegaA$AlphaType,
  levels = c("OmegaA.basic", "OmegaA.FWW", "OmegaA.inferred"),
  labels = c("Basic", "FWW", "GammaExpo"))

dat.exp.omegaNA <- subset(dat, Scenario == "Exponential") %>% pivot_longer(
  cols = c("OmegaNA.basic", "OmegaNA.FWW", "OmegaNA.inferred"),
  names_to = "AlphaType", values_to = "OmegaNA")
dat.exp.omegaNA$AlphaType <- factor(dat.exp.omegaNA$AlphaType,
  levels = c("OmegaNA.basic", "OmegaNA.FWW", "OmegaNA.inferred"),
  labels = c("Basic", "FWW", "GammaExpo"))
```

We first investigate how the real values change according to the proportion of adaptive mutations, $p$ (Figure 3):

```
p.alpha <- ggplot(subset(dat.exp.alpha, AlphaType == "Basic"),
  aes(x = PropPosSel, y = Alpha.simulated,
      col = NRatio, group = NewPopSize)) +
  geom_point() + geom_smooth(method = "nls",
                             formula = y ~ a*log(x+exp(b))+c,
                             method.args = list(
                               start = c(a = 1, b = -3, c = 0),
                               algorithm = "port"),
                             se = FALSE) +
  ylab(expression(alpha^simulated)) +
  xlab(expression(p)) +
  guides(color = guide_legend("log10(Size ratio)")) +
  theme_bw()

p.omegaA <- ggplot(subset(dat.exp.omegaA, AlphaType == "Basic"),
  aes(x = PropPosSel, y = OmegaA.simulated,
      col = NRatio, group = NewPopSize)) +
  geom_point() + geom_smooth(method = "nls",
                             formula = y ~ a*x^b,
                             method.args = list(
                               start = c(a = 1, b = 0.1),
                               algorithm = "port"),
                             se = FALSE) +
```

```
  ylab(expression(omega[A]^simulated)) +
  xlab(expression(p)) +
  guides(color = guide_legend("log10(Size ratio)")) +
  theme_bw()

p.omegaNA <- ggplot(subset(dat.exp.omegaNA, AlphaType == "Basic"),
  aes(x = PropPosSel, y = OmegaNA.simulated,
      col = NRatio, group = NewPopSize)) +
  geom_point() + geom_smooth(method = "nls",
                             formula = y ~ a*x^b+c,
                             method.args = list(
                               start = c(a = 1, b = 0.1, c = 0),
                               algorithm = "port"),
                             se = FALSE) +
  ylab(expression(omega['NA']^simulated)) +
  xlab(expression(p)) +
  guides(color = guide_legend("log10(Size ratio)")) +
  theme_bw()

combinePlots <- function(p1, p2, p3) {
  legend <- get_legend(p1)
  p1 <- p1 + theme(legend.position = "none")
  p2 <- p2 + theme(legend.position = "none")
  p3 <- p3 + theme(legend.position = "none")
  p <- plot_grid(p1, p2, p3, nrow = 3, labels = "AUTO")
  plot_grid(p, legend, nrow = 1, rel_widths = c(4,1))
}

combinePlots(p.alpha, p.omegaA, p.omegaNA)
```

The population size ratio as a clear impact on the real values of $\alpha$ and $\omega_A$, while its effect is more complex for $\omega_{NA}$: for low values of adaptive mutations, $\omega_{NA}$ decreases with the $N$ ratio, while the opposite effect is observed for high proportions of adaptive mutations.

We now check the estimates:

```
p.alpha <- ggplot(dat.exp.alpha,
  aes(x = PropPosSelFactor, y = Alpha-Alpha.simulated, col = NRatio)) +
  geom_point() + geom_abline(slope = 0) +
  geom_smooth(aes(x = PropPosSelOrder), method = "loess", se = FALSE) +
  facet_grid(~AlphaType) +
  ylab(expression(alpha - alpha^simulated)) +
  xlab("Proportion of positively selected mutations") +
  guides(color = guide_legend("log10(Size ratio)")) +
  theme_bw()

p.omegaA <- ggplot(dat.exp.omegaA,
  aes(x = PropPosSelFactor, y = OmegaA-OmegaA.simulated, col = NRatio)) +
  geom_point() + geom_abline(slope = 0) +
  geom_smooth(aes(x = PropPosSelOrder), method = "loess", se = FALSE) +
  facet_grid(~AlphaType) +
  ylab(expression(omega[A] - omega[A]^simulated)) +
  xlab("Proportion of positively selected mutations") +
  guides(color = guide_legend("log10(Size ratio)")) +
  theme_bw()
```

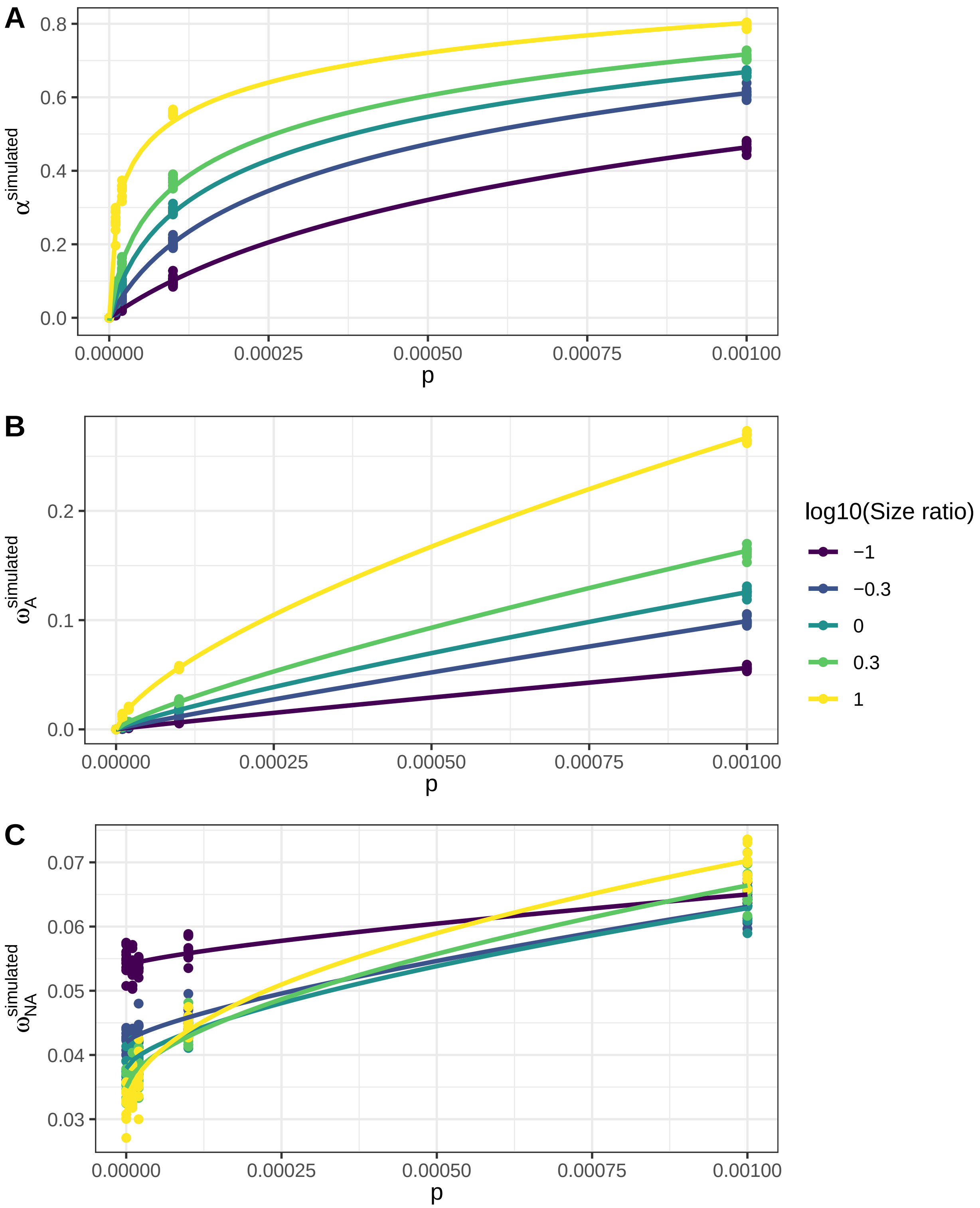


Figure 3: Real values as a function of $p$

```r
p.omegaNA <- ggplot(dat.exp.omegaNA,
  aes(x = PropPosSelFactor, y = OmegaNA-OmegaNA.simulated, color = NRatio)) +
  geom_point() + geom_abline(slope = 0) +
  geom_smooth(aes(x = PropPosSelOrder), method = "loess", se = FALSE) +
  facet_grid(~AlphaType) +
  ylab(expression(omega['NA'] - omega['NA']^simulated)) +
  xlab("Proportion of positively selected mutations") +
  guides(color = guide_legend("log10(Size ratio)")) +
  theme_bw()

combinePlots(p.alpha, p.omegaA, p.omegaNA)
```

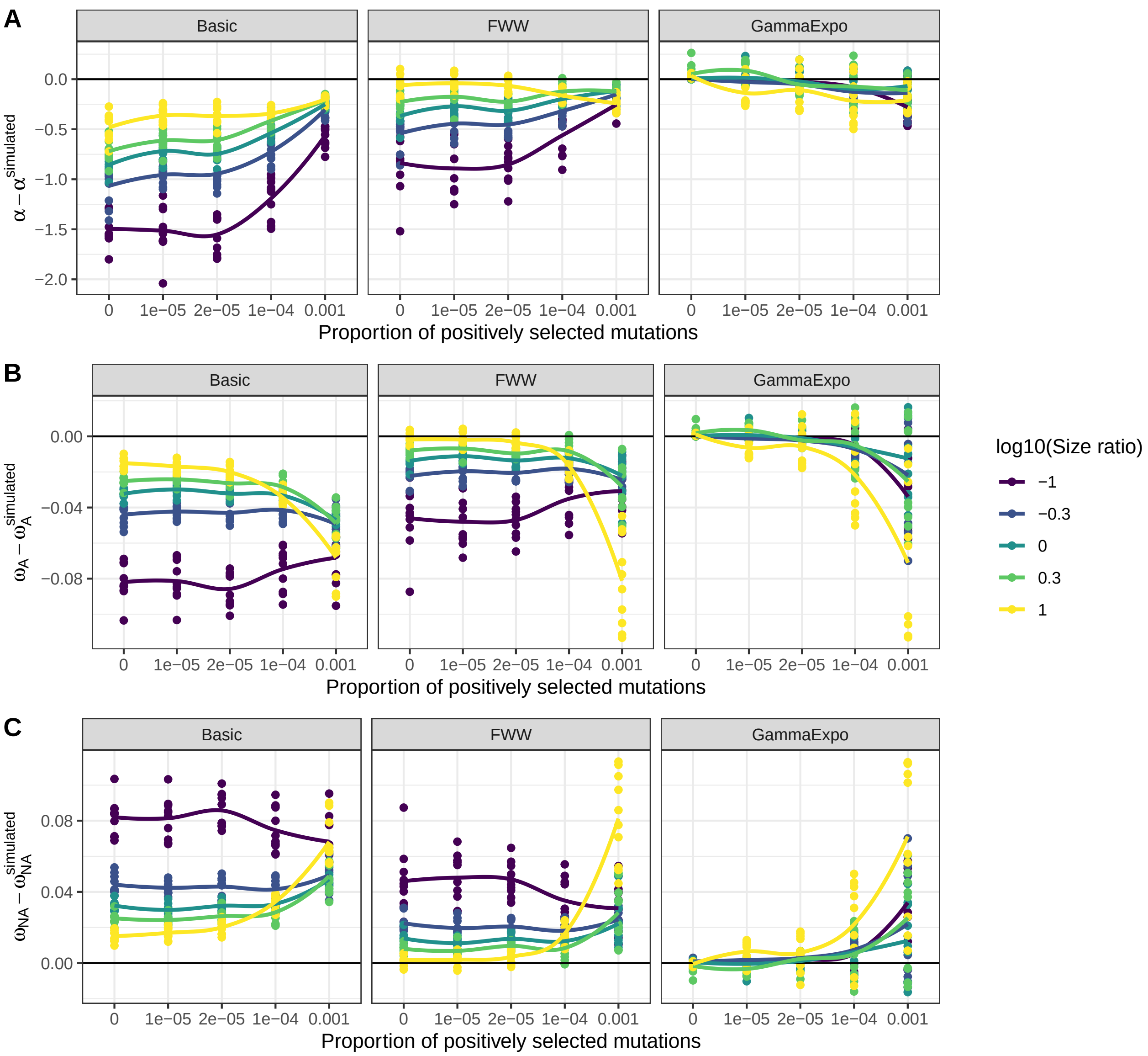


Figure 4: Exponential growth / shrinkage

The effect of the $N$ ratio on the inference of the various parameters is complex (Figure 4):

**Estimation of $\alpha$:**

For the basic and FWW estimates, the $N$ ratio has an increasing effect when the proportion of adaptive mutations is low. Using a model-based inference results in much less biased estimates. A small underestimation bias is still observed for large values of $p$. The under-estimation bias is comparable to or smaller than the one observed with the basic and FWW estimates for large values. This suggests that model-inference does not create biases but rather is better at correcting the estimation for small values.

**Estimation of $\omega_A$ and $\omega_{NA}$:**

The model-based inference recovers the true values when $p$ is small. For large $p$, a biased is observed when $N$ is very different from 1: $\omega_A$ is underestimated and $\omega_{NA}$ is overestimated.

#### Population structure: two demes

Two demes of 10,000 individuals exchange individuals with a certain probability at each generation. More precisely, a new individual has a probability $p_{admixt}$ to have a parent coming from the other deme. Migration is symetric between the two populations. The data sampling is done in one deme only.

```
dat$PropAdmixt[is.na(dat$PropAdmixt)] <- 0

dat.2de.alpha <- subset(dat, Scenario %in% c("Panmictic", "TwoDemes")) %>% pivot_longer(
  cols = c("Alpha.basic", "Alpha.FWW", "Alpha.inferred"),
  names_to = "AlphaType", values_to = "Alpha")
dat.2de.alpha$AlphaType <- factor(dat.2de.alpha$AlphaType,
  levels = c("Alpha.basic", "Alpha.FWW", "Alpha.inferred"),
  labels = c("Basic", "FWW", "GammaExpo"))

dat.2de.omegaA <- subset(dat, Scenario %in% c("Panmictic", "TwoDemes")) %>% pivot_longer(
  cols = c("OmegaA.basic", "OmegaA.FWW", "OmegaA.inferred"),
  names_to = "AlphaType", values_to = "OmegaA")
dat.2de.omegaA$AlphaType <- factor(dat.2de.omegaA$AlphaType,
  levels = c("OmegaA.basic", "OmegaA.FWW", "OmegaA.inferred"),
  labels = c("Basic", "FWW", "GammaExpo"))

dat.2de.omegaNA <- subset(dat, Scenario %in% c("Panmictic", "TwoDemes")) %>% pivot_longer(
  cols = c("OmegaNA.basic", "OmegaNA.FWW", "OmegaNA.inferred"),
  names_to = "AlphaType", values_to = "OmegaNA")
dat.2de.omegaNA$AlphaType <- factor(dat.2de.omegaNA$AlphaType,
  levels = c("OmegaNA.basic", "OmegaNA.FWW", "OmegaNA.inferred"),
  labels = c("Basic", "FWW", "GammaExpo"))
```

We first plot the results for when $p > 0$ (Figure 5):

```
p.alpha <- ggplot(subset(dat.2de.alpha, PropPosSel > 0),
  aes(x = PropPosSelFactor, y = Alpha-Alpha.simulated,
      col = as.ordered(PropAdmixt), linetype = Scenario)) +
  geom_point() + geom_abline(slope = 0) +
  geom_smooth(aes(x = PropPosSelOrder - 1), method = "loess", se = FALSE) +
  facet_grid(~AlphaType) +
  ylab(expression(alpha - alpha^simulated)) +
  xlab("Proportion of positively selected mutations") +
  guides(color = guide_legend("Admixture\nprobability")) +
  theme_bw()

p.omegaA <- ggplot(subset(dat.2de.omegaA, PropPosSel > 0),
```

```
  aes(x = PropPosSelFactor, y = OmegaA-OmegaA.simulated,
      col = as.ordered(PropAdmixt), linetype = Scenario)) +
  geom_point() + geom_abline(slope = 0) +
  geom_smooth(aes(x = PropPosSelOrder - 1), method = "loess", se = FALSE) +
  facet_grid(~AlphaType) +
  ylab(expression(omega[A] - omega[A]^simulated)) +
  xlab("Proportion of positively selected mutations") +
  guides(color = guide_legend("Admixture\nprobability")) +
  theme_bw()

p.omegaNA <- ggplot(subset(dat.2de.omegaNA, PropPosSel > 0),
  aes(x = PropPosSelFactor, y = OmegaNA-OmegaNA.simulated,
      col = as.ordered(PropAdmixt), linetype = Scenario)) +
  geom_point() + geom_abline(slope = 0) +
  geom_smooth(aes(x = PropPosSelOrder - 1), method = "loess", se = FALSE) +
  facet_grid(~AlphaType) +
  ylab(expression(omega['NA'] - omega['NA']^simulated)) +
  xlab("Proportion of positively selected mutations") +
  guides(color = guide_legend("Admixture\nprobability")) +
  theme_bw()

combinePlots(p.alpha, p.omegaA, p.omegaNA)
```

Globally, the graphs are very similar to what we obtain with a panmictic population. Using the GammaExpo model, we see a small underestimation of $\omega_A$ and $\alpha$, and a low overestimation of $\omega_{NA}$ when $p$ is low ($< 1e-6$).

We then plot separately when the real proportion of adaptive mutation is 0 (Figure 6):

```
p.alpha <- ggplot(subset(dat.2de.alpha, PropPosSel == 0),
  aes(x = as.ordered(PropAdmixt), y = Alpha,
      col = as.ordered(PropAdmixt), linetype = Scenario)) +
  geom_point() + facet_grid(~AlphaType) +
  ylab(expression(alpha)) +
  xlab(expression("Admixture probability")) +
  guides(color = guide_legend("Admixture\nprobability")) +
  theme_bw()

p.omegaA <- ggplot(subset(dat.2de.omegaA, PropPosSel == 0),
  aes(x = as.ordered(PropAdmixt), y = OmegaA,
      col = as.ordered(PropAdmixt), linetype = Scenario)) +
  geom_point() + facet_grid(~AlphaType) +
  ylab(expression(omega[A])) +
  xlab(expression("Admixture probability")) +
  guides(color = guide_legend("Admixture\nprobability")) +
  theme_bw()

p.omegaNA <- ggplot(subset(dat.2de.omegaNA, PropPosSel > 0),
  aes(x = as.ordered(PropAdmixt), y = OmegaNA,
      col = as.ordered(PropAdmixt), linetype = Scenario)) +
  geom_point() + facet_grid(~AlphaType) +
  ylab(expression(omega['NA'])) +
  xlab(expression("Admixture probability")) +
  guides(color = guide_legend("Admixture\nprobability")) +
  theme_bw()
```

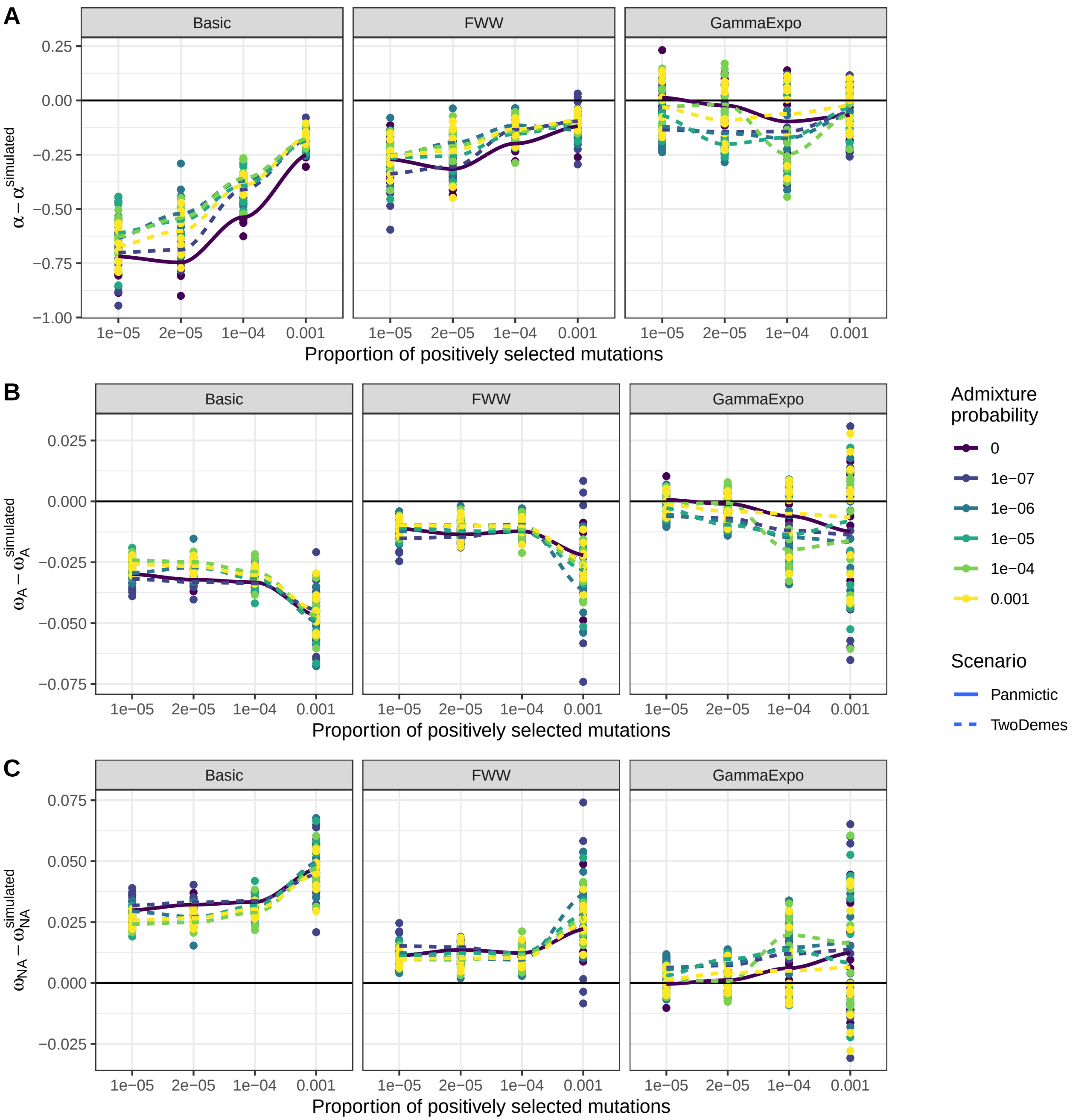


Figure 5: Population structure: two demes

```
combinePlots(p.alpha, p.omegaA, p.omegaNA)
```

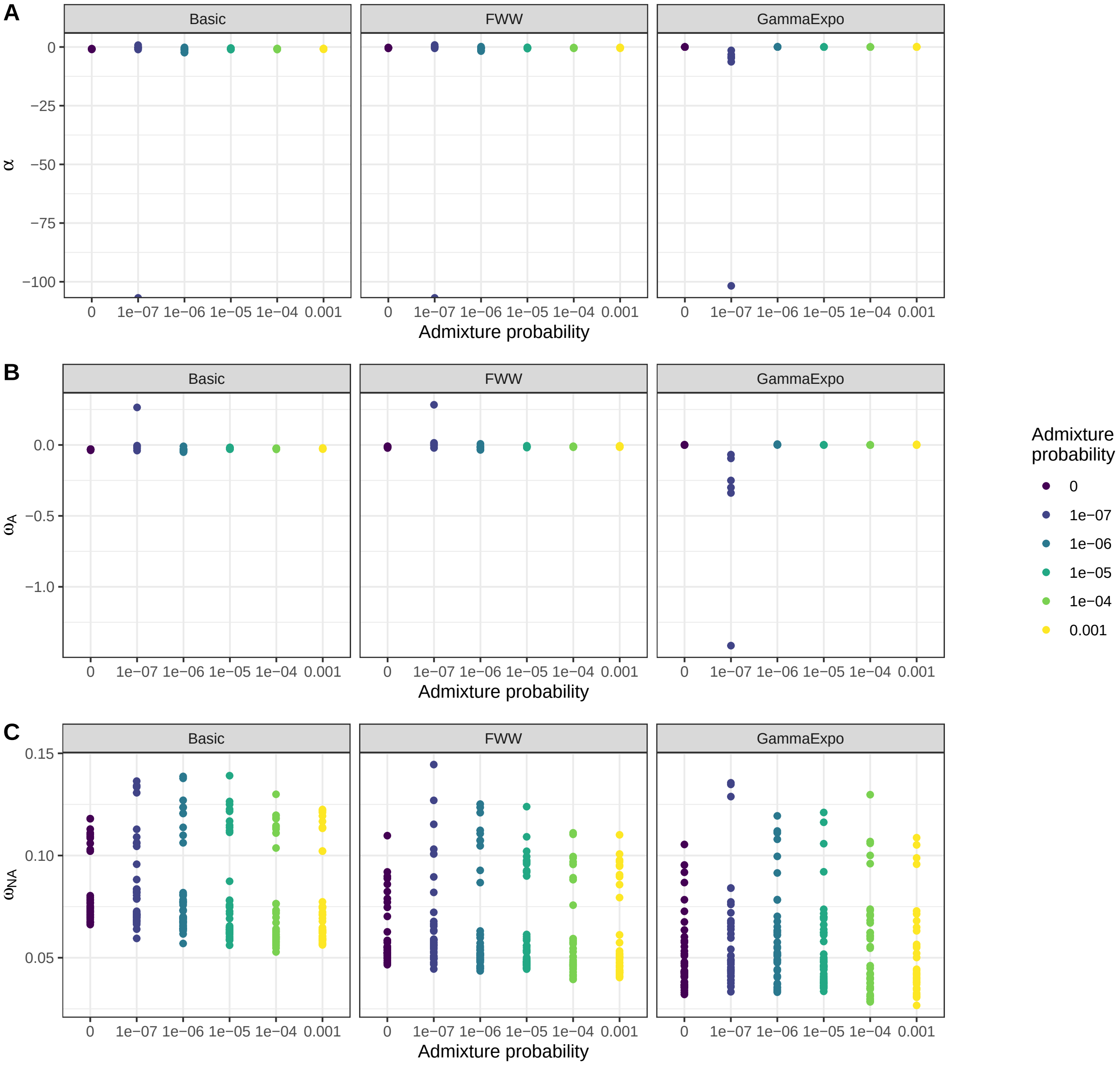


Figure 6: Population structure: two demes, no adaptive mutations

Strong admixture (p < 1e-6) leads to negative $\omega_A$ and $\alpha$ when there is no adaptive mutations. In all other cases, the GammaExpo model provides good estimates.

#### Population structure: two merging populations

In this model, we have two demes exchanging migrants with a fixed proportion of 1e-6. After a certain time $t$, the two populations merge into a panmictic population, which is sampled when a total of 100,000 generations have occurred (Figure 7).

```
dat$MergeTime[is.na(dat$MergeTime)] <- 0

dat.2dm.alpha <- subset(dat, Scenario %in% c("Panmictic", "TwoDemesMerge")) %>% pivot_longer(
```

```
  cols = c("Alpha.basic", "Alpha.FWW", "Alpha.inferred"),
  names_to = "AlphaType", values_to = "Alpha")
dat.2dm.alpha$AlphaType <- factor(dat.2dm.alpha$AlphaType,
  levels = c("Alpha.basic", "Alpha.FWW", "Alpha.inferred"), labels = c("Basic", "FWW", "GammaExpo"))

dat.2dm.omegaA <- subset(dat, Scenario %in% c("Panmictic", "TwoDemesMerge")) %>% pivot_longer(
  cols = c("OmegaA.basic", "OmegaA.FWW", "OmegaA.inferred"),
  names_to = "AlphaType", values_to = "OmegaA")
dat.2dm.omegaA$AlphaType <- factor(dat.2dm.omegaA$AlphaType,
  levels = c("OmegaA.basic", "OmegaA.FWW", "OmegaA.inferred"),
  labels = c("Basic", "FWW", "GammaExpo"))

dat.2dm.omegaNA <- subset(dat, Scenario %in% c("Panmictic", "TwoDemesMerge")) %>% pivot_longer(
  cols = c("OmegaNA.basic", "OmegaNA.FWW", "OmegaNA.inferred"),
  names_to = "AlphaType", values_to = "OmegaNA")
dat.2dm.omegaNA$AlphaType <- factor(dat.2dm.omegaNA$AlphaType,
  levels = c("OmegaNA.basic", "OmegaNA.FWW", "OmegaNA.inferred"),
  labels = c("Basic", "FWW", "GammaExpo"))
```

```
p.alpha <- ggplot(dat.2dm.alpha,
  aes(x = PropPosSelFactor, y = Alpha-Alpha.simulated,
      col = as.ordered(MergeTime), linetype = Scenario)) +
  geom_point() + geom_abline(slope = 0) +
  geom_smooth(aes(x = PropPosSelOrder), method = "loess", se = FALSE) +
  facet_grid(~AlphaType) +
  ylab(expression(alpha - alpha^simulated)) +
  xlab("Proportion of positively selected mutations") +
  guides(color = guide_legend("Merging\ntime")) +
  theme_bw()

p.omegaA <- ggplot(dat.2dm.omegaA,
  aes(x = PropPosSelFactor, y = OmegaA-OmegaA.simulated,
      col = as.ordered(MergeTime), linetype = Scenario)) +
  geom_point() + geom_abline(slope = 0) +
  geom_smooth(aes(x = PropPosSelOrder), method = "loess", se = FALSE) +
  facet_grid(~AlphaType) +
  ylab(expression(omega[A] - omega[A]^simulated)) +
  xlab("Proportion of positively selected mutations") +
  guides(color = guide_legend("Merging\ntime")) +
  theme_bw()

p.omegaNA <- ggplot(dat.2dm.omegaNA,
  aes(x = PropPosSelFactor, y = OmegaNA-OmegaNA.simulated,
      col = as.ordered(MergeTime), linetype = Scenario)) +
  geom_point() + geom_abline(slope = 0) +
  geom_smooth(aes(x = PropPosSelOrder), method = "loess", se = FALSE) +
  facet_grid(~AlphaType) +
  ylab(expression(omega['NA'] - omega['NA']^simulated)) +
  xlab("Proportion of positively selected mutations") +
  guides(color = guide_legend("Merging\ntime")) +
  theme_bw()

combinePlots(p.alpha, p.omegaA, p.omegaNA)
```

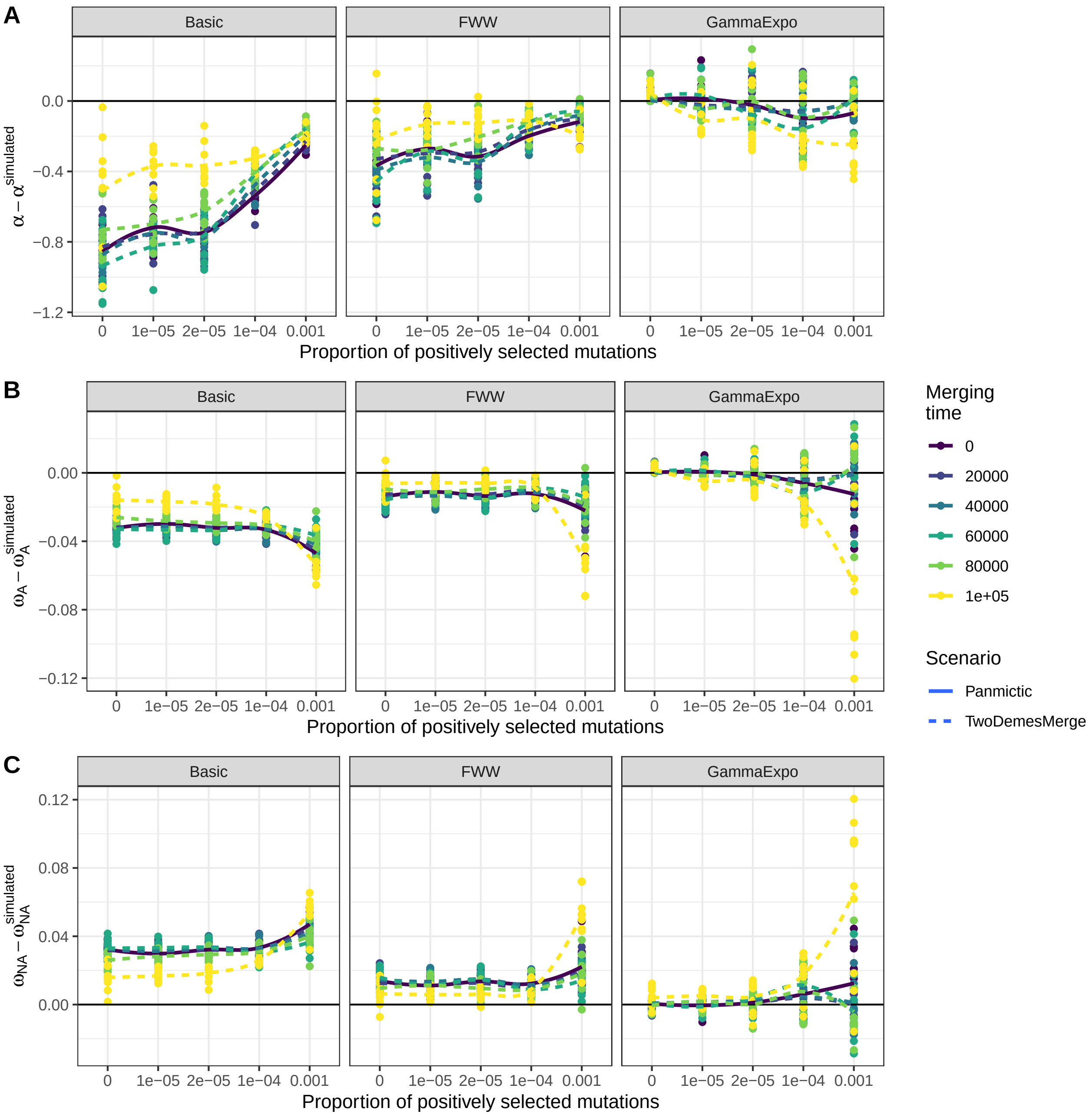


Figure 7: Population structure: two merging demes

We see virtually no effect of population structure. The exception is for a merging time equal to 1e5 and high values of $p$, in which case $\omega_A$ is more strongly underestimated (and $\omega_{NA}$ over-estimated). This scenario corresponds to the case where two populations are mixed in the sample, which we should be able to detect prior to inference.

#### Conclusion

The model-based inference using the GammaExpo* model is very robust to demography and population structure. Grapes will report negative estimates only when the true proportion of adaptive mutations is 0 and there is a strong population structure. Strong departure from a panmictic, constant population generally lead to an underestimation of $\alpha$ and $\omega_A$, so that the model-based estimates can be considered lower bounds of the true values.
